# The genetic architecture of phenotypic diversity in the betta fish (*Betta splendens*)

**DOI:** 10.1101/2021.05.10.443352

**Authors:** Wanchang Zhang, Hongru Wang, Débora Y. C. Brandt, Beijuan Hu, Junqing Sheng, Mengnan Wang, Haijiang Luo, Shujie Guo, Bin Sheng, Qi Zeng, Kou Peng, Daxian Zhao, Shaoqing Jian, Di Wu, Junhua Wang, Joep H. M. van Esch, Wentian Shi, Jun Ren, Rasmus Nielsen, Yijiang Hong

## Abstract

The Betta fish displays a remarkable variety of phenotypes selected during domestication. However, the genetic basis underlying these traits remain largely unexplored. Here, we report a high-quality genome assembly and re-sequencing of 727 individuals representing diverse morphologies of the betta fish. We show that current breeds have a complex domestication history with extensive introgression with wild species. Using GWAS, we identify the genetic basis of multiple traits, including several coloration phenotypes, sex-determination which we map to *DMRT1*, and the long-fin phenotype which maps to *KCNJ15*. We identify a polygenic signal related to aggression with many similarities to human psychiatric traits, involving genes such as *CACNB2* and *DISC1*. Our study provides a resource for developing the Betta fish as a genetic model for morphology and behavior in vertebrates.

## Introduction

The Betta fish (*Betta splendens*), is indigenous to central Thailand and the lower Mekong [1], but is mostly known for its domesticated forms appreciated as an ornamental fish originally bred for its use in gambling matches similar to cock fights [2]. Through captive breeding, a wide variety of behaviors and morphologies have emerged, including variation in aggressiveness, pigmentation, body size, and fin shape. *B. splendens* is easy to breed and maintain, and provides a useful resource for exploration of the genetic basis of behavior and morphology in vertebrates, due to the high degree of intraspecific variability and the vast number of characterized phenotypes. Although researchers have examined the inheritance of body color, fin length and sex in classic crosses in the 1930s to 1940s [3–9], relatively little was known until a recent study investigated the double tail, elephant ear, albino and fin spot phenotypes [10]. Here we report a high-quality chromosomal-level genome assembly of a female *B. splendens*, resequencing data of 727 domesticated individuals, and resequencing data of 59 individuals from six other species in the *B. splendens* complex. We examine the evolutionary relationship and origins among breeds and use association mapping to identify the genetic basis of a number of different traits including sex determination, fin morphology, coloration, body size, and aggressiveness and other behaviors.

## Results

### Genome assembly, annotation and quality assessment of the betta fish

We generated a high-quality chromosomal-level assembly of the betta fish using a multifaceted sequencing and assembling workflow, including PacBio reads, Illumina reads, Hi-C reads, 10X Genomics reads and BioNano optical mapping (see Materials and methods, Supplementary Text 2, Figs. S1, S2 and Tables S1-S7 for details). The final assembled genome totaled 451.29 Mb on 21 chromosomes with contig and scaffold N50 reaching 4.07 Mb and 19.63 Mb, respectively. CEGMA [11] (Core Eukaryotic Genes Mapping Approach) confirmed 239 of 248 (96.4%) complete core eukaryotic genes, while BUSCO [12] (Benchmarking Universal Single-Copy Orthologs) covered 2,521 of 2,586 (97.5%) single-copy orthologous genes (Table S1), indicating high completeness of the genome and gene annotation. These statistics suggest our assembly is a more complete genome than obtained in previous assemblies of *B. splendens* deposited in GenBank (Accession: GCF_900634795.2 and GCA_003650155.3) (Table S1). We compared the *B. splendens* genome to 13 other teleost genomes (Fig. 1A), and found a large expansion of the *SLC4A* gene family which is associated with bicarbonate transporters that regulate intracellular pH levels [13]. *B. splendens* is known for its adaptation to hypoxic and black (acidic) water conditions. These comparative genomic results are discussed in detail in Supplementary Text 3, Figs. S3-S4, Supplementary Data 3.

**Fig. 1.**
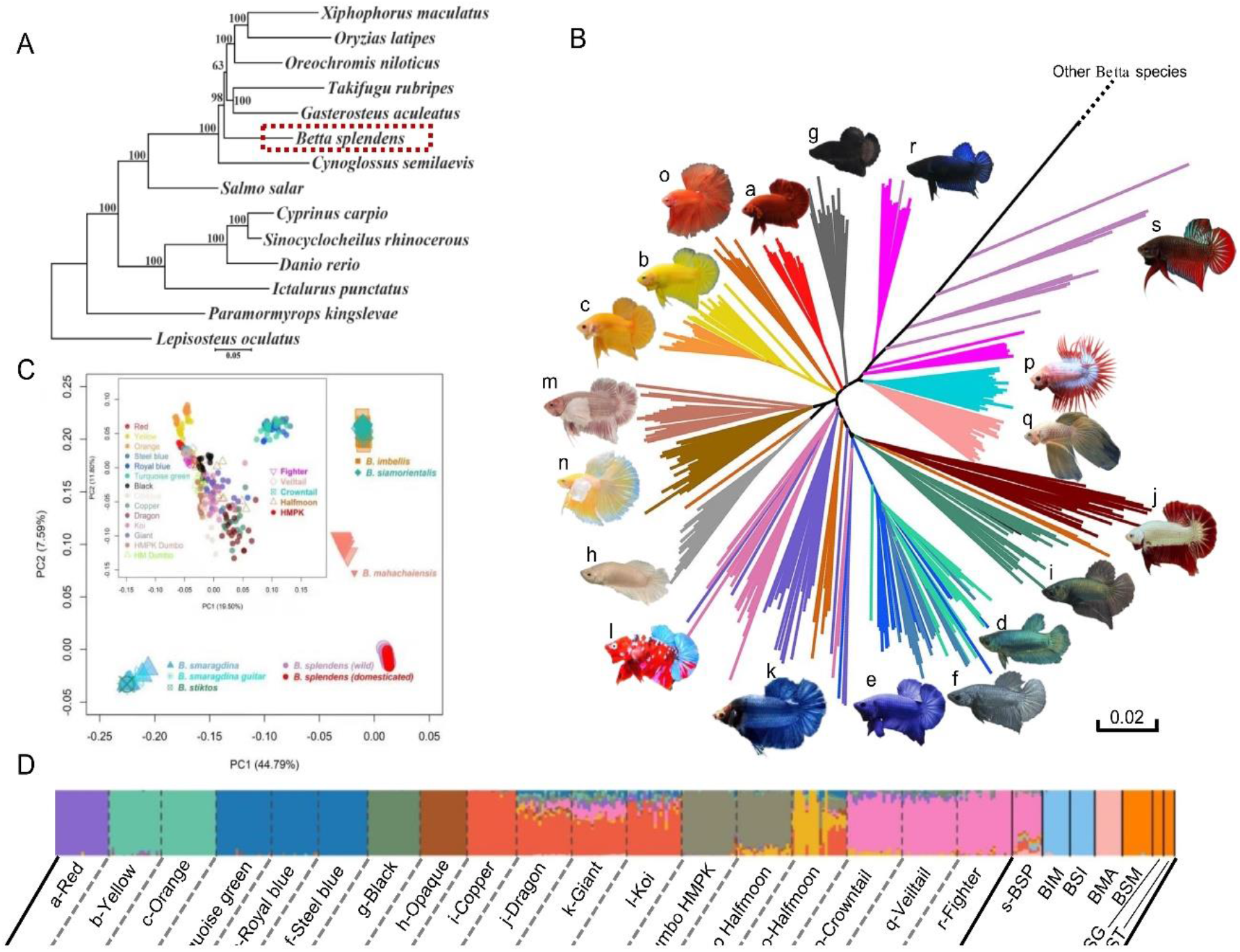
Phylogeny and population structure of the betta fish (*Betta splendens*). (A) The phylogeny of teleost including the betta fish constructed with single-copy genes across the genome. (B) Neighbor-joining tree of concatenated sequences, representing an average genomic tree, for domesticated and wild forms of *Betta splendens*. The tree is truncated at the branch connecting to individuals of other *Betta* species, and the untruncated tree can be found in Fig. S6. The letters by the fish photos correspond to the numbers in (D). (C) Principal components analysis (PCA) of the Betta species complex. The inset is the PCA for all *Betta splendens* individuals. (D) *ADMIXTURE* analysis of the *Betta splendens* breeds and their closely related wild species. *K* = 12 is presented here and the results with *K* varying from 2 to 15 are in Fig. S17. Acronyms for wild species are: BSP, *Betta splendens*; BIM, *Betta imbellis*; BSI, *Betta siamorientalis*; BMA, *Betta mahachaiensis*; BSM, *Betta smaragdina*; BSG, *Betta smaragdina guitar*, BST, *Betta stiktos*.

### Phenotypic diversity, genome-resequencing, and variant calling

We collected 14 breeds of Betta fish differing in tail type (Fig. 1B and S5; Table S8; Supplementary Text 1) and performed whole genome resequencing of the 727 individuals resulting in ~2.7 Tb clean sequencing data with depths ranging from 3X to 34X (average 6.7X). To elucidate the population history of the *B. splendens* complex and the origins of the domesticated Betta fish, we analyzed six wild species and 14 domesticated breeds using 59 individuals from the wild species and 20 random individuals from each breed. Principal component analysis (PCA) and a Neighbor-joining tree from concatenated sequences, representing the average genomic coalescent tree, based on 11.37 million variants (Materials and methods) confirmed that the domesticated breeds cluster together (Fig 1B and 1C). These results are compatible with the hypothesis that all current breeds of the Betta fishes were domesticated from the same group of wild *B. splendens. B. imbellis* and *B. siamorientalis* also cluster together in the tree (Fig. S8), while the lineages of *B. stiktos* and *B. smaragdina guitar* are interspersed and not monophyletic with respect to each other in the average genomic tree, suggesting that they all should be considered different varieties of the *B. smaragdina* species (Fig 1C and S6).

Treemix analyses (Fig. S7-S15) suggest extensive introgression between wild species in the *B. splendens* complex, in particular gene flow from *B. mahachaiensis* into the lineages of Royal-blue, Steel-blue and Turquoise-green breeds (Fig. S16). ABBA-BABA tests support this conclusion (Table S9), e.g., with a D(H1: Crowntail, H2: Royal; H3: Mahachaiensis, H4: Smaragdina) = 0.31 (Z score=21.51). These results show that other species in the *B. splendens* complex have contributed to the genomic make-up of some breeds of *Betta splendens*, suggesting that they may also have contributed to phenotypic variation in these breeds.

### Diversification of the Betta fish during domestication

Several clades of the Fighter breed fall as outgroups to the rest of the domesticated breeds in the average genomic tree suggesting that the Fighter breed represents an early domesticated form and that the earliest domesticated Betta fish in fact were breeds selected for fighting [2]. Other breeds falling towards the root of the tree include Veiltail and Crowntail. Population structure, as revealed by both *ADMIXTURE* analyses (Figs. 1D, S17) and average genomic trees (Fig. 1D), suggests that breeds defined by coloration and morphology generally clustered together, although we note that this conclusion might be affected by the sampling strategy employed here. The HMPK breeds, which are short-fin types distinct from the Fighter and wild *B. splendens* by a rounded tail shape and body shape, also cluster based on appearance, mostly related to colors (Fig. 1B). The Red, Yellow, Orange, Turquoise-green, Royal-blue and Steel-blue breeds show reduced nucleotide diversity (Figs. S18), suggesting that these strains experienced strong bottleneck effects in their domestication history.

### Regulatory variants in *DMRT1* mediate the male-heterogamety of *B. splendens*

Using GEMMA (Materials and methods) [14], we performed association mapping using all 727 individuals consisting of 590 males and 137 females to map the sex determination (SD) locus (Fig. 2A). A kinship matrix inferred by GEMMA and the first three PCs were used as covariates to minimize the effects of population stratification (Materials and methods). We find a highly significant map location at position 2.83-2.89 Mb on chromosome 2 (*P* = 4.80E-63) and no evidence of statistical inflation (Fig. S19). 530 out of 537 individuals with heterozygous genotypes of the lead variant (chr2:2,839,325) are phenotypically males, which strongly supports the hypothesis of male heterogamety (XY/XX) (Table S10).

**Fig. 2.**
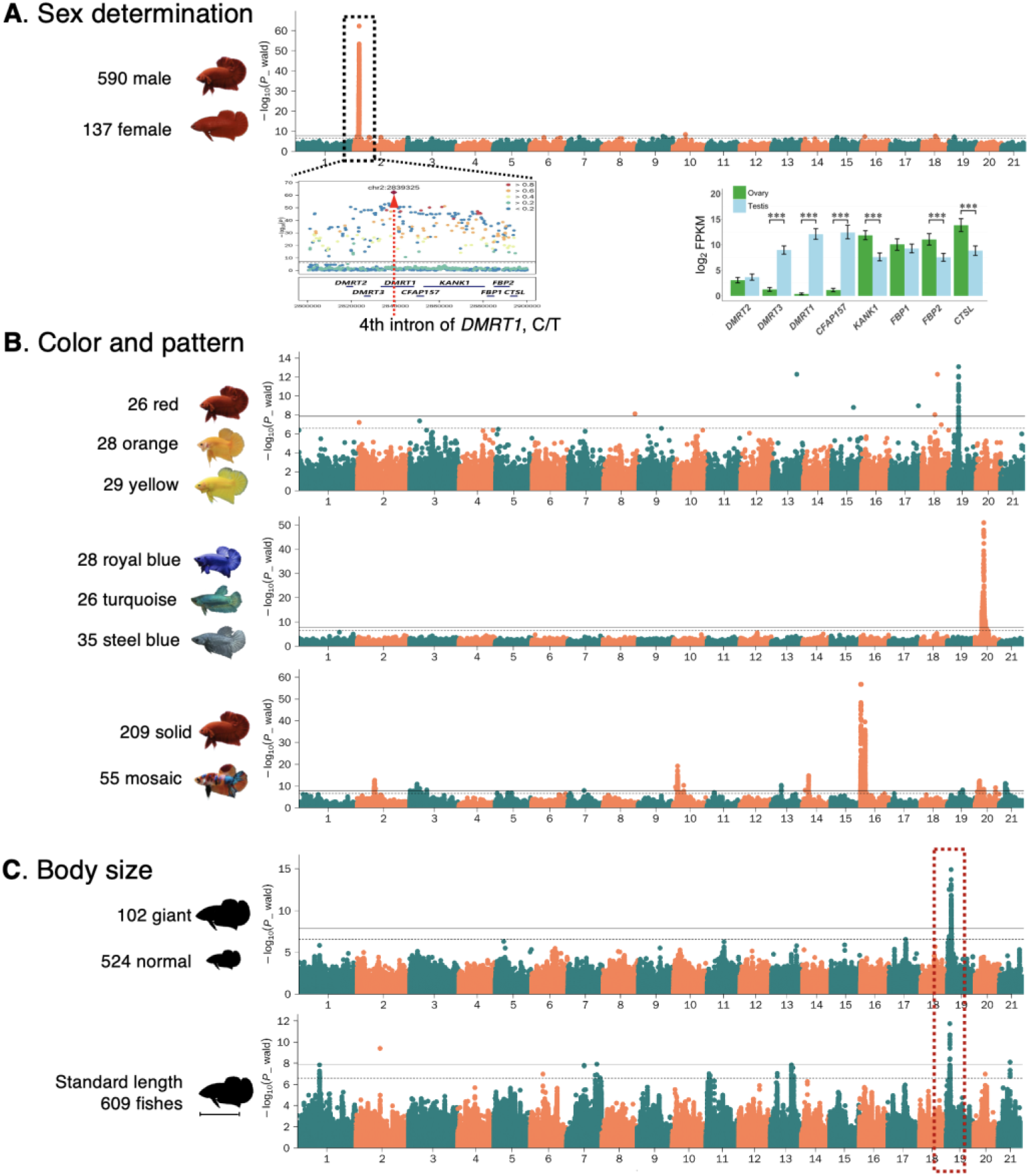
Genome-wide association studies and locus analysis of sex determination, skin color and pattern, and body size in *Betta splendens*. (A) GWAS of sex determination and expression profiles in testis and ovary for genes in the associated locus. A single genome-wide significant signal is identified, with a lead SNP located on the intron 4th of the *DMRT1* gene. (B) Manhattan plots for GWAS of the red, orange and yellow color phenotypes (upper), the royal blue, turquoise green, and steel blue color phenotypes (middle), the solid color and mosaic pattern phenotypes (lower). (C) Manhattan plot for GWAS of body size. The GWAS in the upper plot is conducted with case (giant) and control (non-giant), and the lower is conducted with the standard body length. The shared genome-wide significant signals are highlighted with the red box. The GWAS results of other body size indices are in Fig. S35 and S38. The horizontal lines on Manhattan plots represent genome-wide (solid line) and suggestive genome-wide (dash line) significance level, respectively.

The most strongly associated variants cluster in introns of *DMRT1* and *KANK1* (Fig. 2B). *KANK1* function is in cytoskeleton formation by regulating actin polymerization [15], making it a less like candidate for SD. *DMRT1*, in contrast, is a well-known gene contributing to the sex determination in fish [16], bird [17] and reptiles [18]. Also, mRNA-sequencing shows that *DMRT1* is predominantly expressed in testis (Fig. 2A). *DMRT1* in the Betta fish is evolutionarily close to the medaka fish *DMY* gene (Fig. S24), which was the first sex-determining gene identified in teleost [19]. The presence of males with homogametic female genotype (52/590) may suggest that environmental factors, such as temperature, also could play a role in sex determination similarly to many other fish [20].

### Gain of function of *KCNJ15* contributes to the overgrowth of all fins

The most striking morphological difference among the Betta fish are in the fins, especially the caudal fin. Veiltail, Crowntail, and Halfmoon show a remarkable outgrowth in dorsal, anal and caudal fins compared to the Fighter and HMPK breed (Fig. S23). A pattern of dominant inheritance in inter-breed crosses [21], suggests a shared genetic basis for long-fin growth. To map the underlying genetic variation, we used the same procedure as for SD, dividing specimens into long-fin individuals (N=201) and short-fin individuals (N=525) (detailed descriptions of all phenotypic measurements are in Supplementary Text 4). We notice that in all GWAS presented in this paper, we were comparing genetically differentiated breeds or morphotypes resulting in a substantial challenge in controlling for population structure. However, using standard methods for correction (Materials and methods), we obtained reasonable QQ-plots and small inflation factors (Fig. S24). Nonetheless, our design may not be able to distinguish between traits that have been selected in the exact same morphotypes/breeds during domestication, and in this sense resembles a classical F_ST_ scan for identifying adaptive genetic differences.

The GWAS for long-fin versus short-fin identified an extremely significant locus (*P* = 9.28E-159) centered at position 9.60 Mb on chromosome 17 (Fig. 3A). There are three lead variants in perfect LD: one synonymous (9,590,677 bp, exon 3, C1551T), one intronic (9,590,546 bp, intron 2, G>A) in *SMG8*, and one in the 3’UTR of *KCNJ15* (9,596,738 bp, G>A). All three variants perfectly distinguish between long-fin and short-fin, and are, therefore, considered candidate causal mutations. *SMG8* acts as a regulator of kinase activity involved in non-sense-mediated decay of mRNAs [22], and is an unlikely candidate. In contrast, *KCNJ15* encodes a potassium channel, which is a much better candidate, as genes encoding potassium channels have already been identified to cause various long-fin phenotypes in zebrafish, including *KCNK5B* [23], *KCNH2A* [24] and *KCNJ13* [25]. We explored the expression profiles between long caudal fin and short caudal fin by RNA-Seq, and there are no significant expression differences in *SMG8* (Fig. 3C). However, *KCNJ15* is highly expressed in long-fin breeds but has no intact transcripts detected in the short-fin breeds (corresponding to the wild *B. splendens* phenotype) (Fig. S25). RT-PCR (Fig. 3D and 3E) shows that this long-fin specific transcript was also present in paired fins (pectoral and pelvic fins), suggesting a developmental alteration in all appendages. More discussion of *KCNJ15*, including expression patterns, is provided in Supplementary Text 8.

**Fig. 3.**
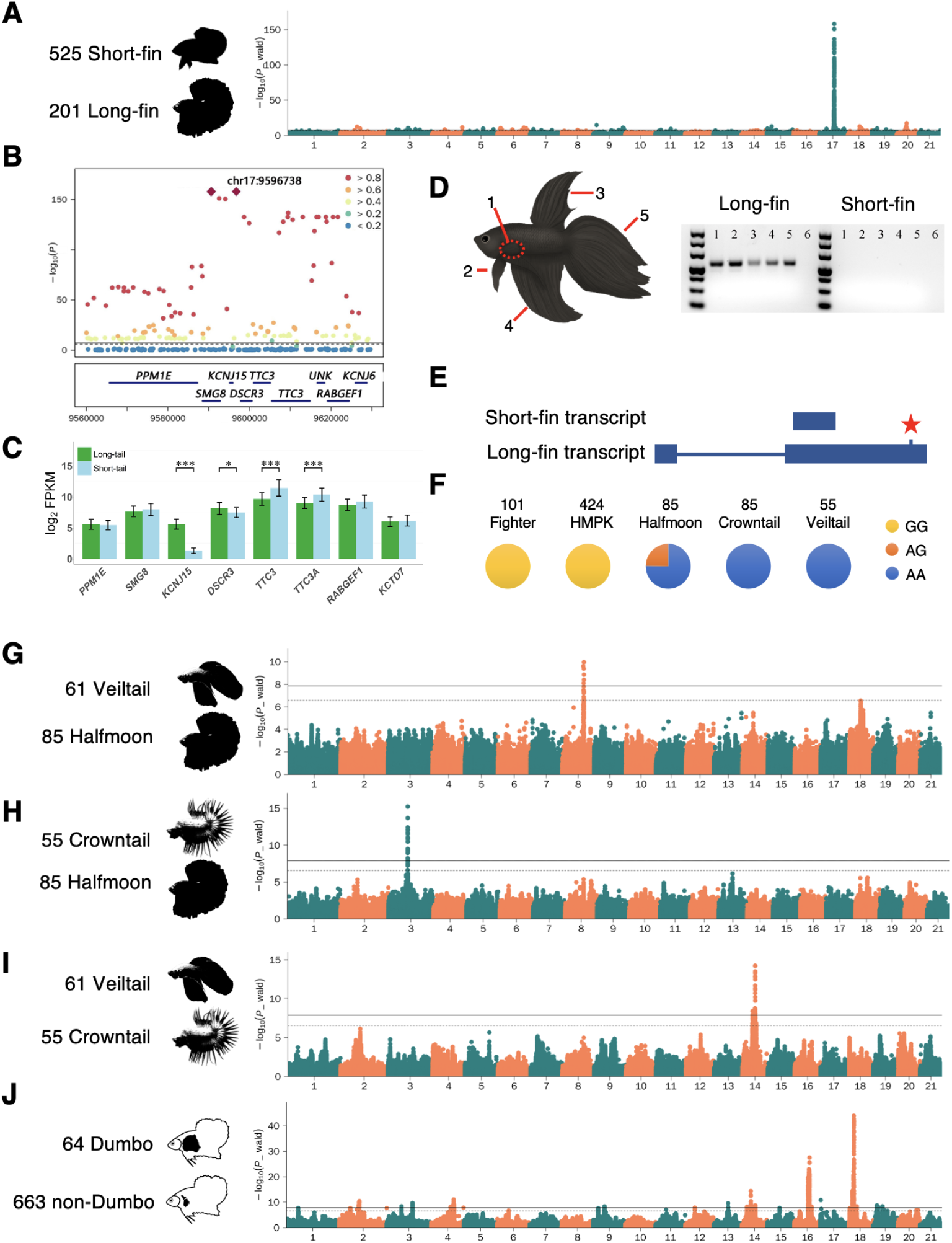
GWAS of fin morphology in *Betta splendens*. (A) Manhattan plot for the long-fin vs. short-fin morphology. (B) LocusZoom plot for the most significant genome-wide association signal. (C) The expression profiles in caudal fin for genes in the associated locus. (D) RT-PCR analysis of the *KCNJ15* gene in different fins of the fish (right), number 1-5 on the gel picture are fin codes shown on the left, and the 6th lane is the blank control. (E) The location of the peak SNP and gain of a new transcript in long-fin fishes. (F) Genotype frequencies at the peak SNP in five different *Betta splendens* breeds. (G-I) Manhattan plots of GWAS for different fin morphs among long-fin breeds. (J) Manhattan plots of GWAS for the Dumbo, i.e., outgrown pectoral fin phenotype.

### Veiltail, Crowntail and Halfmoon phenotypes

Within the breeds with long-fins, i.e., the Veiltail, Crowntail and Halfmoon, we identified three independent loci including 12.21 – 12.27 Mb on chromosome 3, 13.48 – 13.79 Mb on chromosome 8, and 8.25 – 8.40 Mb on chromosome 14, using one breed as case and another as control (Figs. 3G, 3H, 3I, S26-S28). Veiltail was the first long-fin breed recorded [9] and selected for full extension in caudal fin ray spreading as Halfmoon. The 300-kb long locus on chromosome 8, harboring 29 associated SNPs and four protein-coding genes, (Fig. S26) is associated with the phenotypic difference between Veiltail and Halfmoon. The locus at 12.21 – 12.27 Mb on chromosome 3 is associated with a reduction of webbing tissue in all fins comparing Halfmoon and Crowntail (Fig. S27) and includes FRMD6 involved in actomyosin structure organization and with loss of expression displaying epithelial-to-mesenchymal transition features [26]. In Supplementary text 8, we discussed the candidate genes in these peak regions and also association mappings for other in-depth phenotyping of fin morphology, including fin ray number and fin ray branching patterns.

### The ‘Dumbo’ phenotype

The pectoral fin in teleosts is homologous to the anterior appendages in amphibian, reptile and mammal [27]. One breed, the ‘Dumbo’, is characterized by the overgrowth of its paired pectoral fins, with more and elongated fin rays (Fig. 1B, S5). GWAS using the Dumbo as cases and other non-Dumbo as controls identified two strong signals (Fig. 3J), are located on chromosome 16 (8.87 – 9.72 Mb, *P* = 2.99E-28, Fig. S29) and 18 (4.25 – 4.47 Mb, *P* = 1.08E-44, Fig. S30) respectively. Wang et al. [10] investigated the same phenotype using F_ST_ scan and located a 1.3Mb region on chromosome 16 (chromosome 9 in their assembly) that contains our GWAS peak. They suggested, based on gene expression analyses, that the causal gene could be *kcnh8* [10]. However, this gene resides outside of the associated locus on chromosome 16 identified in this study (Fig. S29). Moreover, no differential expression was detected in the pectoral fin tissue for *kcnh8* when comparing Dumbo and non-Dumbo breeds (Fig. S31). The lead SNP in our study is in a region containing a cluster of eight genes from the *HOXA* gene family which is essential for forming fin skeleton and digits in teleost [28]. For the signal on chromosome 18, the lead SNP is located on the 3’UTR of *FBXL15* (F-box and leucine-rich repeat protein 15), a gene involved in dorsal/ventral pattern formation and bone bass maintenance [29], rendering this a strong candidate gene. More discussion can be found in Supplementary Text 8.

### The Giant phenotype

Body size exhibits polygenic inheritance in most organisms [30]. However, a Giant mutant in the Betta fish shows significant enlargement in total length, standard length, height and weight compared to other breeds (Fig. S34). A GWAS comparing Giant size vs. normal size and GWAS using quantitative traits (including total length, standard length, height and weight), identified a shared significant locus at position 2.03 – 2.26 Mb on chromosome 19, which explained 7.83-16.34% of phenotypic variance (Fig. 2C, S35). Conditional on the top SNP (chr19:2,130,270), all other associated variants in this locus lose their significance in all body size GWAS, suggesting a common genetic basis underlying the body enlargement (Fig. S36).

### Colors and color patterning in the Betta fish

In teleost, erythrophores and xanthophores produce red and yellow pigments, respectively, and orange color is generated with a mixture of red and yellow pigments [31]. In the Betta fish, solid red is inherited dominantly over solid yellow [32]. Association mapping of the three colors identifies a locus harboring 93 variants, all residing in the *RNF213*, at position 5.83 Mb of chromosome 19 (Fig. 2B, S44). *RNF213* encodes a large cytoplasmic protein that is involved in angiogenesis and the non-canonical Wnt signaling pathway in vascular development [33].

Mosaic color pattern is found elusively in several domesticated fishes, like Koi carp [34], medaka [35], and the Betta fish (Fig. 2B), however, the underlying molecular basis has never previously been investigated. Using a GWAS with a case-control design between solid (N=209) and mosaic colors (N=55), we identified two highly significant loci on chromosome 16 (0.07-0.43 Mb and 2.57-2.79 Mb; Fig. 2B) discussed in more detail in Supplementary Text 10, together with other GWAS results for coloration.

Crossing Royal-blue males with Royal-blue females produces Turquoise-green, Royal-blue and Steel-blue offspring, at a Mendelian ratio of 1:2:1 [5]. Previous studies show that these three phenotypes are based on structural coloring and that all breeds have erythrophores, xanthophores, melanophores and iridophores in the scales, except Steel-blue which lacks erythrophores on the upper layer of the scales [36]. A GWAS with the three colors identified a single locus (Fig 2B, S47) at 5.03 – 5.26 Mb on chromosome 20 and the same locus together with another locus on chromosome 7 were also found when using Copper and Steel-blue as cases and controls (Fig. S50). This locus on chromosome 20 contains 13 protein coding genes (Fig. 2B, Fig. S48) including *MTHFD1L* (methylenetetrahydrofolate dehydrogenase 1 like) involved in the synthesis of tetrahydrofolate [37], which is engaged in the *de novo* assemble of purines, a key component of iridophores. Other candidate loci are discussed in Supplementary Text 10.

### Aggression

Aggressive behavior is a complex phenotype involving genetics, endocrinology, neurophysiology and metabolism [38]. Male winners of one-vs-one fights have been selected by breeders for improved combat performance [39]. We performed a GWAS using Fighters (N=101) as cases and other breeds (N=626) as controls. A total of 346 polymorphisms including 26 exonic (8 nonsynonymous and 18 synonymous), 176 intronic and 144 intergenic within/near 82 protein-coding genes were identified (Supplementary Data 1). This result suggests a polygenic basis for the behavioral differences. The top signals show significant enrichment for genes involved in morphine addiction, circadian entrainment, oxytocin signaling, GABAergic synapse, estrogen signaling, vasopressin-regulated water reabsorption and axon guidance (Supplementary Data 5), which are also enriched in human and mouse aggression studies [40]. Furthermore, the top candidate genes include *CACNB2*, associated with several psychiatric disorders in humans [41], *DISC1* associated with schizophrenia, bipolar disorder, and recurrent major depression [42], and *AVPR2* with homologs that in knockout mice display reduced anxiety, impaired reciprocal social interaction and decreased social recognition [43, 44]. Supplementary Text 11 discusses possible candidate genes in more detail.

To further investigate behaviors associated with fighting, we measured 11 different behaviors displayed during simulated fighting (Figs. 4B-J, S52, Supplementary Text 11). Fish from the *Betta* genus, including the Betta fish, are mostly facultative air breathers [45]. We recorded the times of air breathing of each Betta fish (N=467) during fighting within one minute in our study and performed an association mapping on it (Figs. 4J, 4L, S53). Nine polymorphisms upstream of *SLC16A1* at around 4.79 Mb and three intronic variants of *CDH4* centered at 15.24 Mb on chromosome 7 were significantly associated with the frequency of air-breathing (Figs. 4L, S54). *SLC16A1* is a rapid monocarboxylate transporter which moves many monocarboxylates such as lactate and pyruvate across the plasma membrane [46], and mutations in this gene are associated with erythrocyte lactate transporter defect [47]. We hypothesize that the *SLC16A1* alleles are associated with higher frequency of air-breathing and may contribute to more efficient metabolism of the lactate-related substrates produced by muscle contraction during fighting.

**Fig. 4.**
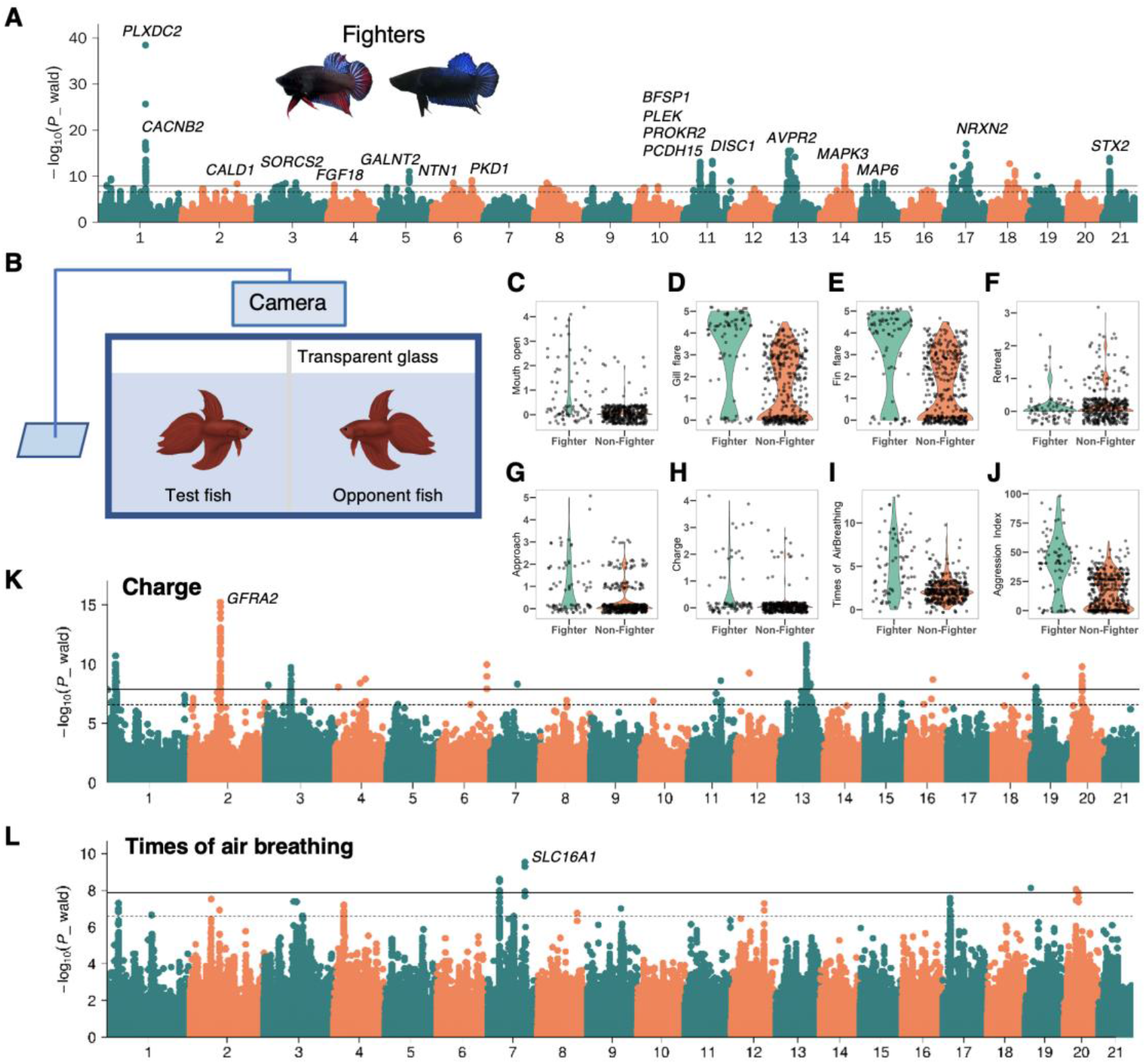
GWAS of aggression behaviors in *Betta splendens*. (A) Manhattan plot of GWAS of fighter versus non-fighter fishes. Two representative morphs of fighter fishes are shown as inset. (B) Experimental setup to quantify the aggressiveness of *Betta splendens* individuals. Aggressiveness indices of test fishes are recorded in the presence of an opponent fish. (C-J) Boxplot of eight aggressiveness phenotypes in fighter versus non-fighter fishes. (K-L) Manhattan plots of GWAS using the charge score and the times of air breathing, respectively.

A GWAS for the charging behavior during fighting (Figs. 4H, 4K) identifies multiple significant lead SNPs in/near *GFRA2*, a gene encoding a protein that plays a key role in the control of neuron survival and differentiation [48]. Additional association mapping results are described in Supplementary Text 11 for other behaviors.

## Discussion

The Betta fish has a complex domestication history involving introgression from other wild species and intensive selection on a variety of phenotypes. A number of traits seem to be likely monogenic, including sex-determination which we map to *DRMT1* and caudal fin size which we map to *KCNJ15*. Other traits show a more polygenic inheritance pattern, particularly the important aggression trait. Our results form a basis for introducing the betta fish as a potential model organism for understanding the genetic basis of a host of different traits in vertebrates, including coloration and patterning, skeletal development of fin/limbs, and behavioral traits such as aggression. As an easy breeder in captivity, the Betta fish has substantial potential as a new model system to augment existing vertebrate models such as zebrafish and mice.

