## Supplementary Data for "The genetic architecture of phenotypic diversity in the betta fish (*Betta splendens*)"

Science
AAAS

### Supplementary Materials for

#### The genetic architecture of phenotypic diversity in the betta fish (*Betta splendens*)

Wanchang Zhang1†, Hongru Wang2†, Débora Y. C. Brandt2, Beijuan Hu1, Junqing Sheng1, Mengnan Wang1, Haijiang Luo1, Shujie Guo1, Bin Sheng1, Qi Zeng1, Kou Peng1, Daxian Zhao1, Shaoqing Jian1, Di Wu1, Junhua Wang1, Joep H. M. van Esch6, Wentian Shi4, Jun Ren3, Rasmus Nielsen2, 5*, Yijiang Hong1*

**This PDF file includes:**

Materials and Methods

Supplementary Texts S1 to S11

Figs. S1 to S56

Tables S1 to S13

Captions for Data S1 to S5

**Other Supplementary Materials for this manuscript include the following:**

Data S1 to S5:

1. Results of genome-wide association studies of all the recorded traits in the Siamese fighting fish.
2. Results of ABBA-BABA tests.
3. Expanded and contracted genes and families in the Siamese fighting fish.
4. Differentially expressed genes in brain between Fighter and Non-Fighter.
5. Functional enrichment of the identified genes by GWAS between the Fighter and Non-Fighter breeds.

**Material and Methods** 7
1. Fish samples 7
2. DNA extraction, library construction and sequencing 7
3. Hi-C library preparation and sequencing 7
4. RNA extraction and sequencing 8
5. RNA-Seq data analysis 8
6. Genome assembly 8
7. Genome annotation 9
Repeat annotation 9
Non-coding RNA annotation 9
Gene prediction 9
Gene function annotation 10
8. Phylogeny of teleost 10
Identification of orthologs 10
Phylogenetic tree construction and divergence time estimation 10
Expansion and contraction of gene families 11
9. Population genetics 11
Read mapping 11
SNP and genotype calling 11
Principal Component Analysis 13
NJ tree 13
ADMIXTURE 13
Nucleotide diversity 13
TreeMix 13
ABBA-BABA 13
10. Phenotyping 13
11. GWAS 14
12. RT-PCR to quantify gene expression 14

**Supplementary Figures** 15
Fig. S1. Genomic landscape of annotated protein-coding genes and non-coding RNA elements in the Siamese fighting fish genome. 15
Fig. S2. Genomic landscape of repeat sequences in the Siamese fighting fish genome. 16
Fig. S3. Time-calibrated phylogeny of teleosts. 17
Fig. S4. A schematic illustration presenting the expanded and contracted genes related to muscular function in the Siamese fighting fish. 18
Fig. S5. Illustration of samples used in this study. 19
Fig. S6. Neighbor-joining tree of the Betta fish. 20

Fig. S7. Treemix plot of Betta fish without migration. 21
Fig. S8. Treemix plot of Betta fish with one migration edge allowed. 22
Fig. S9. Treemix plot of Betta fish with two migration edges. 23
Fig. S10. Treemix plot of Betta fish with three migration edges. 24
Fig. S11. Treemix plot of Betta fish with four migration edges. 25
Fig. S12. Treemix plot for Betta fish with five migration edges. 26
Fig. S13. Treemix plot for Betta fish with six migration edges. 27
Fig. S14. Treemix plot for Betta fish with seven migration edges allowed. 28
Fig. S15: Treemix plot for Betta fish with eight migration edges allowed. 29
Fig. S16: ABBA-BABA test for introgressions between B. mahachaiensis and domesticated B. splendens 30
Fig. S17: ADMIXTURE analysis varying K from 2 to 15 31
Fig. S18: Nucleotide diversity across breeds of domesticated Betta splendens. 32
Fig. S19: QQ plot for sex GWAS. 33
Fig. S20: Synteny at the sex determining locus containing DMRT1. 34
Fig. S21: Sequencing depth and GC content in the sex determining region in the vicinity of the DMRT1 gene of the Siamese fighting fish in this study. 35
Fig. S22: Neighbor-joining tree of putative Betta and Medaka sex determining genes. 36
Fig. S23: Caudal fin length in five breeds of the Siamese fighting fish. 37
Fig. S24: QQ plot for long vs. short fin GWAS. 38
Fig. S25: RNA-Seq reads mapped to the KCNJ15 gene region in short-fin (HMPK) and long-fin (Halfmoon) fishes. 39
Fig. S26: Regional plot of the locus with the strongest association in a GWAS between Veiltail and Halfmoon breeds. 40
Fig. S27: Regional plot of the locus with the strongest association in a GWAS between Crowntail and Halfmoon breeds. 41
Fig. S28: Regional plot of the locus with the strongest association in a GWAS between Veiltail and Crowntail breeds. 42
Fig. S29: Regional plot of the region on chromosome 16 associated with the Dumbo phenotype. 43
Fig. S30: Regional plot of the region on chromosome 18 associated with the Dumbo phenotype. 44
Fig. S31: Expression profile of genes in the locus associated with the Dumbo phenotype in the pectoral fins of Dumbo and non-Dumbo phenotypes. 45
Fig. S32: Manhattan and QQ plots for GWAS for traits related to fin rays in the Siamese fighting fish. 47
Fig. S33: Regional plot of the locus with the strongest association in a GWAS for maximum number of dorsal fin ray splitting in the Siamese fighting fish. 48
Fig. S34: Body size measurements in the Giant HMPK compared to other breeds. 49
Fig. S35: Manhattan plots for the body size GWAS. 50

Fig. S36: GWAS for body size conditioning on top SNP associated with body size. 51
Fig. S37: QQ-plot of the body size GWAS results. 52
Fig. S38: GWAS of other body size traits: BMI, Total length/Height and Weight/Height. 53
Fig. S39: Regional plot of the locus with the strongest association in a GWAS for standard length in the Siamese fighting fish. 54
Fig. S40: Expression in muscle and brain tissues of genes in the 119kb region associated with body size. 55
Fig. S41: GO terms enriched in genes differentially expressed in muscle between the Giant-HMPK and HMPK. 56
Fig. S42: GO terms enriched in genes differentially expressed in brain between the Giant-HMPK and HMPK. 57
Fig. S43: Synteny analysis of the locus associated with the Giant phenotype. 58
Fig. S44: GWAS for red, yellow and orange color breeds in the Siamese fighting fish. 59
Fig. S45: Regional plot of the locus with the strongest association in a GWAS between Orange, Yellow and Red breeds in the Siamese fighting fish. 60
Fig. S46: Regional plot of the locus with the strongest association in a GWAS between Orange, Yellow and Red breeds in the Siamese fighting fish. 61
Fig. S47: GWAS for Turquoise green, Royal blue and Steel blue color breeds in the Siamese fighting fish. 62
Fig. S48: Regional plot of the locus with the strongest association in a GWAS between Turquoise green, Royal blue and Steel blue breeds in the Siamese fighting fish. 63
Fig. S49: A schematic of the breeding of Copper from Steel blue. 64
Fig. S50: GWAS of Copper versus Steel blue breeds. 65
Fig. S51: GWAS results of eye color in the Siamese fighting fish. 66
Fig. S52: Boxplot for aggression related phenotypes composing the aggression index. 67
Fig. S53: Manhattan plot of GWAS for aggression score and the phenotypes composing it. 68
Fig. S54: Quantile-quantile (QQ) plot for GWAS of times of breathing air in one minute in the Siamese fighting fish. 69
Fig. S53: GWAS results when conditional on the top sex variant in the four breeds 70
Fig. S54: GWAS results using different PCs for Fighter vs Non-Fighter. 71

**Supplementary Tables** 72
Table S1. Statistics of three Siamese fighting fish genome assemblies. 72
Table S2. Sequencing platforms and data output 73
Table S3. Repeat annotation of the Siamese fighting fish genome 74
Table S4. Transposable element annotation of the Siamese fighting fish genome 75
Table S5. Gene annotation of the Siamese fighting fish genome 76
Table S6. Non-coding RNA annotation of the Siamese fighting fish genome 77
Table S7. Statistics of functional annotation of structural genes in the Siamese fighting fish genome 78
Table S8. Description of all samples included in this study 79

Table S9. ABBA BABA tests for introgression between B. mahachaiensis and Turquoise, Steel blue, Royal blue and Copper breeds of Betta splendens. The tests are in the form of D(W, X; mahachaiensis, outgroup) where the outgroup is a B. smaragdina individual and W are Turquoise, Steel blue, Royal blue and Copper individuals and X are individuals of all other breeds. 80
Table S10. Genotype frequencies at top sex-associated SNP (chr2:2839325) in males and females 80
Table S12 GWAS summary for Mosaic pattern 81
Table S13 miRNA binding to KCNJ15 using RNAhybrid and miRanda 82

**Supplementary Text** 83
Supplementary Text 1. Background on Betta fish 83
Supplementary Text 2: Genome assembly and genome annotation results 85
Genome assembly 85
Genome annotation 85
Supplementary Text 3: Phylogeny and expanded and contracted gene families. 87
Phylogeny for B. splendens 87
Expanded gene families related to muscular, neuronal and solute carrier functions 87
Gene loss related to ion exchange pathway 89
Supplementary Text 4: Phenotyping 90
Morphology 90
Aggressive behaviours 90
Supplementary Text 5: An overview of GWAS setups 92
Body size 92
Color 92
Fins 93
Supplementary Text 6. Population genetics of Betta splendens 94
Supplementary Text 7: GWAS of sex determination 96
Supplementary Text 8: Candidate loci in GWAS of fin morphology 98
The KCNJ15 gene associated with fin length 98
Associated genomic loci with Veiltail, Crowntail and Halfmoon caudal fin type 99
Associations with the “Dumbo” phenotype 100
Supplementary Text 9: Candidate loci in GWAS of body size 101
Supplementary Text 10: Candidate loci in GWAS of coloration 103
Supplementary Text 11: Candidate loci in GWAS of aggression behaviours 106
GWAS signals for Fighter vs non-Fighter breeds 106
GWAS on the quantitative indices of aggression 108

**Supplementary Data** 109

Supplementary Data 1 Results of genome-wide association studies of all the recorded traits in the Siamese fighting fish. 109

Supplementary Data 2 Results of ABBA-BABA tests. 109

Supplementary Data 3 Positive selected, expanded and contracted gene and families in the Siamese fighting fish. 109

Supplementary Data 4 Differentially expressed genes in brain between Fighter and Non-Fighter 109

#### Material and Methods

1. Fish samples

Two female HMPK Siamese fighting fishes used for de novo genome assembly came from an inbred solid red $F_2$ cross bred in-house. All the 727 domesticated individuals used for genome resequencing were brought from the Yuexiu pet market in Guangzhou (China), and the 59 wild individuals from *B. splendens* complex were brought from commercial exporters in Bangkok (Thailand). Fishes are anesthetized and then fin clips are collected and preserved in ethanol in a -80°C freezer until DNA extraction. All the management of fishes are under the approval of the Ethics Committee for Animal Experiments of Nanchang University, China (No. SYXK-2015-0001).

2. DNA extraction, library construction and sequencing

To generate DNA sequences for genome assembly, high molecular-weight DNA was isolated from the muscle tissue of the two selected samples using MagAttract HMW DNA kit (QIAGEN, Germany), obtaining DNA fragments with average length of ~100 kb. Specifically for the two fish individuals, one fish sample was used for generating PacBio, Illumina and 10X Genomics data, and the other fish for Hi-C and BioNano data. For PacBio Sequel sequencing, two 20-kb-insert-size SMRTbell libraries were prepared and sequenced on the PacBio Sequel platform. For 10X Genomics sequencing, a GEM reaction and library preparation were conducted using size-selected DNA with a length of approximately 50-kb. For Illumina long-range paired-end sequencing, libraries were barcoded and paired-end sequenced with the Rapid method on an Illumina HiSeq X Ten platform. For BioNano optimal map construction, the library was constructed using the BspQl enzyme (New England Biolabs) with an appropriate label density (14.5 labels per 100 kb) to digest long-range DNA fragments. The detailed information regarding insert sizes, data output and genome coverage is listed in Table S2.

For population genomic resequencing, genomic DNA was extracted from fin tissue using the proteinase-K/phenol-chloroform method. Sequencing libraries with insert size 350 bp were constructed and then sequenced on a Illumina HiSeq X Ten platform to generate ~5X coverage data for each fish with 150bp paired-end reads. All the library preparation and sequencing procedures described here were conducted by Novogene Bioinformatics Institute (China).

3. Hi-C library preparation and sequencing

The library was constructed following standard protocols(*1*). Fish muscle tissue was used and the Hi-C sequencing libraries were amplified by PCR for 12-14 cycles before sequenced on Illumina HiSeq X Ten platform (pair-end 150 bp).

4. RNA extraction and sequencing

Four independent RNA libraries were prepared: (1) library of pooled RNA samples of 11 tissues (brain, liver, muscle, eye, skin, scale, fin, intestine, testis, ovary, embryo) of one HMPK individual was constructed and subjected to both full-length transcriptome sequencing using the PacBio Sequel platform and short reads sequencing with the Illumina Hiseq 2500 platform. (2) Libraries of RNA samples of caudal fin tissues from HMPK (n=5) and Halfmoon (n=5) individuals were prepared and sequenced on the Illumina Hiseq 2500 platform. (3) Libraries of RNA samples of muscle and brain tissues from giant size (n=5) and normal size (n=5) individuals were prepared and sequenced in the Illumina Hiseq 2500 platform. (4) microRNA libraries were prepared for caudal fin tissues of HMPK (n=5) and Halfmoon (n=5) individuals and sequenced in Illumina Hiseq 2500 platform to explore microRNA binding to the 3'UTR of *KCNJ15* (Supplementary Text 8).

5. RNA-Seq data analysis

The raw sequencing data were first processed by removing adapters and low-quality reads. The resulting clean paired-end reads were mapped to our assembly using Hisat2 (v.2.0.5) (*2*). Then the reads number mapped to each gene was counted using featureCounts (v1.5.0-p3) (*3*) before the FPKM (expected number of Fragments Per Kilobase of transcript sequence per Millions base pairs) was calculated. Differential gene expression analysis was performed with the edgeR package (3.18.1) (*4*). The P values were adjusted for multiple testing using the Benjamini & Hochberg method. Genes with adjusted P value < 0.05 and fold change greater than 2 were considered as differentially expressed.

6. Genome assembly

There were five steps involved in the genome assembly: (1) De novo assembly using PacBio long reads; (2) error correction with Illumina short reads and PacBio reads; (3) remove heterozygous haplo-contigs; (4) pre-scaffolding with 10X linked reads; (5) scaffolding with BioNano optimal map and (6) super-scaffolding to chromosome level with Hi-C.

(1) We sequenced the female solid red HMPK individual using PacBio Sequel SMRTbell platform with an insertion of 20 kb and obtained 53.27 Gb sequencing data. We used a in-house modified Falcon (v0.3.0) (https://github.com/PacificBiosciences/FALCON/)(*5*) to preliminary assemble the PacBio reads into 1,029 contigs with total and N50 length reaching 492.05 Mb and 3.98 Mb, respectively. Main parameters in Falcon are "--max_diff 100 --max_cov 100 --min_cov 2 --bestn 10 --min_len 6000". (2) Quiver5 (*6*) was employed to correct the sequencing errors of the PacBio consensus sequences. We used 53.05 Gb sequencing data of the same sample from Illumina HiSeq X Ten to correct the errors of the assembled contigs under the default parameters of Pilon (v1.18) (*7*). (3) Then, purge_haplotigs (*8*)was applied to remove the heterozygous contigs under parameters "-l 25 -m 70 -h 125 -a 75" and resulted in 273 contigs with total and N50 length reaching 448.89 and 4.99 Mb, respectively. (4) We used fragScaff (version2.1, -m 3000 -q 30 -E

30000 -o 60000 -C 5 -j 1.5) (*9*)to associate the 273 contigs into scaffolds based on 114.98 Gb data of linked reads generated from 10X genomics strategy. A total of 227 scaffolds were obtained with total, N50 and max length reaching 449.22 Mb, 6.57 Mb and 19.28 Mb. (5) BioNano Solve (v3.3) (*10*) was adopted to retrieve the optical maps generated by the Saphyr system (BioNano Inc.) onto the scaffolds, and this updated the total, N50 and max scaffold length to 451.33, 12.16 and 27.50 Mb. (6) The Lachesis (v-201701) (*11*) software was used to improve the genome assembly to the chromosome level with the 52.58 G Hi-C data, resulting in a genome assembly with scaffold N50 of 19,630,546 bp. Genome assembly results are presented in Supplementary Text 2 and Tables S1 and S2.

7. Genome annotation

Repeat annotation

Tandem repeats were extracted with Tandem Repeats Finder (TRF, v4.09) (*12*) by *ab initio* prediction. For other repeats, we employed both *de novo*-based and homology-based approaches. For the *de novo*-based approach, we used RepeatModeler (v2.0.1) (*13*), LTR_FINDER (v1.07) (*14*)and RepeatScout (v1.0.5) (*15*) to build the de novo repeat library. For the homology-based approach, we used RepeatMasker (4.1.0) (*16*) against the Repbase TE library (*17*) and RepeatProtein Mask (4.1.0) (*18*) against the TE protein database (*16*).

Non-coding RNA annotation

Four types of non-coding RNAs including tRNA, rRNA, miRNA and snRNA were annotated in the assembly. tRNAscan-SE (v1.4) (*19*) (http://lowe1ab.ucsc.edu/tRNAscan-SE/, default options) was used to find the tRNA sequences according to its structural features. We use the rRNA sequences of related species as the references and find the highly conserved rRNA sequences in our genome assembly through Blast (v2.2.26). Using the covariance model of Rfam family, the Infernal ("INFERence of RNA ALignment") (http://infernal.janelia.org/) software (v14.1) [infernal] (*20*) was applied to predict the sequences of miRNAs and snRNAs under default parameters.

Gene prediction

De novo predictions, homolog-based predictions and RNA-Seq based predictions were employed to annotate the protein coding genes in the Siamese fighting fish genome. Five ab initio gene prediction programs were used to predict genes, including Augustus (v3.0.2)(*21*), Genescan (v1.0)(*22*), Geneid (v1.4) (*23*), GlimmerHMM (v3.0.2) (*24*), and SNAP (v2006-07-28) (*25*). Protein sequences of 13 homologous species were downloaded from Ensembl or NCBI. Homologous sequences were aligned against the repeat-marked *Betta splendens* genome using TBLASTN (v2.2.26, e-value ≤ 1e-05). Genewise (v2.4.1, "-tfor-genesf") (26) was employed to predict gene models based on the alignment sequences. Trinity (v2.1.1, "--normalize_reads --

full_cleanup --min_glue 2 --min_kmer_cov 2 --KMER_SIZE 25 --no_distributed_trinity_exec") (*27*) and SMRTLink (v6.0, https://www.pacb.com/support/software-downloads/, default parameters) were used to assemble the RNA-Seq data, and then PASA software (http://pasapipeline.github.io/) (*28*) was used to improve the gene structures. A weighted and non-redundant gene set was generated by EVidenceModeler (v1.1.1) (*29*), which merged all gene models predicted by the above three approaches. Combining with transcript assembly, PASA adjusted the gene models generated by EVM. The final reference gene set obtained 25,104 protein coding genes (Table S5).

Gene function annotation

The longest transcript of each gene was retained for gene annotation. The translated protein sequence of the transcript was used to 'blastp' search the Swissprot database (http://www.uniprot.org) (*30*)with the cutoff "E-value <10-5", then the gene annotations and gene ontology terms from the closest hit were extracted for gene prediction. The conserved protein domains and functional motifs of the transcripts were identified with InterproScan (*31*) with default settings. The gene pathway annotation was obtained by mapping to the KEGG database (http://www.genome.jp/kegg/) (*32*). Finally, 22,788 protein coding genes (90.77% of total genes) were funcional annotated.

8. Phylogeny of teleost

Identification of orthologs

Gene families were identified by OrthoMCL (v1.4) (*33*). First, nucleotide and protein data of 13 species representative of different teleost families (*Cyprinus carpio*, *Danio rerio*, *Sinocyclocheilus rhinocerous*, *Cynoglossus semilaevis*, *Oryzias latipes*, *Salmo salar*, *Gasterosteus aculeatus*, *Oreochromis niloticus*, *Takifugu rubripes*, *Paramormyrops kingsleyae*, *Ictalurus punctatus*, *Xiphophorus maculatus* and *Lepisosteus oculatus*) were downloaded from Ensembl (Release 70) and NCBI to co-analyze with the *Betta splendens* genome assembly. The longest transcript of a gene was retained among the different alternative splicing transcripts, genes with ≤ 30 amino acids were discarded, and then an "all against all" BLASTP comparison was performed followed by filtering using "E-value ≤ 1E-07" cutoff. The blastp alignments were clustered using OrthoMCL (v1.4) (*33*) with a 1.5 inflation index. After clustering, 24,159 gene clusters and 465 single-copy orthologs were detected across the 14 teleost species including *B. splendens*.

Phylogenetic tree construction and divergence time estimation

The aforementioned 465 shared single-copy orthologs were utilized to estimate a teleost phylogeny. Coding sequences (CDS) of these orthologs were aligned by MUSCLE (v3.7, "-maxiters 2") (*34*). With these CDS alignments, a maximum-likelihood phylogenetic tree was constructed using

RAxML (v7.2.3, "-m GTRGAMMA -p 12345 -x 12345 -f ad") (*35*). Then, the program MCMCTree of PAML (v4.5)(*36*) (http://abacus.gene.ucl.ac.uk/software/paml.html) was applied to estimate divergence times among 14 species with parameters "burn-in=100,000, sample-number=100,000, and sample-frequency=2". Seven calibration points were selected from the TimeTree website (http://www.timetree.org) as normal priors to restrain the age of the nodes, including 76-111 Mya between *Xiphophorus maculatus* and *Oryzias latipes*, 87-151 Mya between *Oreochromis niloticus* and *Oryzias latipes*, 101-136 Mya between *Takifugu rubripes* and *Gasterosteus aculeatus*, 88-114 Mya between *Cynoglossus semilaevis* and *Gasterosteus aculeatus*, 186-227 Mya between *Betta splendens* and *Salmo salar*, 17-51 Mya between *Cyprinus carpio* and *Sinocyclocheilus rhinocerous*, and 87.4-124.7 Mya *between Danio rerio* and *Sinocyclocheilus rhinocerous.*

Expansion and contraction of gene families

The aforementioned gene clustering file from the OrthoMCL and time-calibrated phylogeny were used to detect expanded and contracted gene clusters with CAFE (v3.1, "load -p 0.05 -t 4 -r 10000; lamda -s") (*37*), which uses a birth-death model to infer the birth-death rate of gene families in a given tree and infers ancestral states of gene family sizes in the internal nodes of the tree. CAFE then infers the rate of change in gene family size along each branch, and tests the null hypothesis of no change in gene family expansion rate using a likelihood ratio test. We used this test and a 5% significance level to determine significance.

#### 9. Population genetics

##### Read mapping

Clean reads were mapped onto the newly generated assembly in this study with the Burrows-Wheeler Aligner (v0.7.8)(*38*). Duplicated reads were marked using the MarkDuplicates tool from the Picard software package (*39*) with default options. Local realignment around indels was performed using the IndelRealigner tool from the GATK software package (v3.3.0)(*40*).

##### SNP and genotype calling

The procedure followed the "reference SNP generation"→"Base Quality Score Recalibration (BQSR)"→"SNP and genotype calling"→"Variant Quality Score Recalibration (VQSR)" pipeline.

Two SNP panels were generated in this study, denoted as "3.77 M" and "11.90 M" respectively based on the number of SNPs in these panels. Different samples were used for calling these SNPs and different filtering was applied. The "3.77M" panel is used for GWAS and the "11.90M" is used for population genetic analysis.

(a) For both SNP sets, an initial SNP reference panel was constructed using two independent SNP detection pipelines involving GATK (version 4.0)(*41*) and Platypus (v0.1.5) (*42*) were employed respectively. The union set of the detected SNPs were taken as the final reference panel. For the GATK pipeline, the UnifiedGenotyper model with the parameters "genotyping_mode DISCOVERY glm BOTH" were used and the VCF was filtered with "DP<4, QD<4.0, FS>200, ReadRosRankSum<-20.0, InbreedingCoeff<-0.8". For the Platypus pipeline, the parameters "minPosterior 5, subthreshold 0.01, abThreshold 0.01" were used, followed by filtering with "miss 0.05, maf 0.1".

(b) BQSR: the SNP reference panel from (a) was used as known sites for calibrating base quality scores in the BAM files. The BaseRecalibrator program from the GATK (version 4.0) package(*41*), with default parameter settings, was used.

(c) SNP and genotype calling: The base-quality recalibrated BAM files from (b) were used as input for SNP and genotype calling. First, individual GVCF files were generated using HaplotypeCaller from GATK (version 4.0) (*41*), followed by joint genotype calling with GenotypeGVCF for the populations.

(d) VQSR: First, using the SNP panel generated in (a), the "VariantRecalibrator" program was run with parameters "known=false, training=true, truth=true, prior=12.0 -an DP -an QD -an FS -an SOR -an ReadPosRankSum -an MQRankSum -mode SNP -tranche 100.0 -tranche 99.9 -tranche 99.0 -tranche 90.0" to generate "snps.recal" and "snps.tranches" files which were used for "ApplyVQSR" to calibrate variant quality scores with the filter set to 99.

*"3.77 M" panel*

In this panel, 727 domesticated *Betta splendens* individuals were included. An initial VCF with 12.65M SNPs was filtered using VCFtools options "--maf 0.03 --minqQ 30" and PLINK option "--geno 0.05". After also filtering to keep only bi-allelic sites, a panel of 3.77M SNPs was obtained. Next, this panel was imputed and phased using Beagle (version 4.1) (*43*). Annovar was used to annotate the variant dataset functionally(*44*).

*"11.90 M" panel*

In this panel, 410 individuals, including ~20 randomly picked individuals from each breed of the Siamese fighting fish, and all the 59 individuals from six wild species of the *B. splendens* complex were used for SNP detection. SNP calling was performed with the pipeline described above, followed by hard filtering in GATK (version 4.0)(*41*) with default parameters. Sites with minor allele frequency lower than 0.03, non-biallelic, with missing rate greater than 50% were also removed. The final dataset contains 11.90M SNPs with 2.68% missing data. The panel was LD-pruned using PLINK (1.07)(*45*) with parameter "–indep-pairwise 50 5 0.2", obtaining a smaller dataset containing 721K SNPs. Both the 11.90 M and the 721K SNPs dataset were used for following population genetic analysis.

Principal Component Analysis

Principal component analysis (PCA) was conducted using smartpca from EIGENSOFT (6.1.4)(*46*) on the 721k LD-pruned SNPs from all wild and domesticated Betta fishes. We also performed an independent PCA including only the domesticated breeds, for which 351 domesticated individuals were extracted from the VCF file of the 11.90 M dataset, re-filtered using PLINK options "-maf 0.03 -geno 0.8", and variants on unplaced scaffold were also removed.

NJ tree

The Neighbor-Joining tree was built using PHYLIP (v3.697) (*47*)from the identical-by-state distance matrix computed by PLINK (v1.07) (*45*)from the 11.90 M dataset. The tree was visualized using FigTree (v.1.4.3)(*48*).

ADMIXTURE

Unsupervised structure analysis was performed using *ADMIXTURE* (v1.3.0) (*49*) on the 721k SNPs dataset, varying the number of clusters (*K)* from 2 to 15.

Nucleotide diversity

Nucleotide diversity in each breed was estimated by VCFtools (v0.1.13) (*50*) on the 11.90 M dataset.

TreeMix

The TreeMix (v1.13)(*51*) analysis was performed on the 721K dataset with "-bootstrap 1000 -k 500" parameters for the Betta species complex with *Betta smaragdina* set as the outgroup.

ABBA-BABA

The ABBA-BABA tests were conducted using ANGSD (v0.928) (*52*)with the "-doAbbababa" argument on mapped bam files with block size of 500 kb. This procedure is based on sampling single reads and is not subject to genotype calling errors. We randomly sample three individuals from each breed or species for the tests, and only conclude a pattern when it is consistent across different individuals tested.

10. Phenotyping

Individuals were phenotyped for GWAS of traits related to body size, color, fin shape and rays, aggressive behaviour and sex. A detailed description of how each phenotype was defined is available in Supplementary Text 4.

11. GWAS

Genome-wide association mapping was conducted using GEMMA (v.0.96) (*53*) on the 3.77M SNP dataset. A detailed description of the phenotypic measurements across breeds is in Supplementary Text 4 and a discussion of GWAS setups of various scenarios are specified in Supplementary Text 5. The Wald test is used to determine the association strength (*P*-values) and Bonferroni correction was used to define the genome-wide significance level (cutoff = -log10(0.05/number of variants) = 8.46). The effect of population stratification was corrected with a kinship matrix and the first 3, 10 or 20 PCs covariates. Only subtle differences were observed and the GWAS results presented are based on three PCs. We note here that the GWAS design used for most phenotypes is based on a case-control design comparing different sets of breeds. While we find that the p-values generally appear to be well-calibrated by standard methods as determined by inflation factors and inspection of qq-plots (see Figs. S19, S24, S32, S37, S38, S44), we note that this design is unlikely to be able to distinguish between phenotypic differences caused by differential selection during domestication on a phenotype different from the focal phenotype, if this phenotype has been selected in the exact same breeds. As such, our GWAS design shares some similarities with classical $F_{ST}$ scans for identifying selection.

12. RT-PCR to quantify gene expression

RNA molecules were extracted from all five different fin tissues of one Halfmoon and one HMPK individual respectively using TRIzol reagent (Life technologies). Reverse transcription was performed using PrimeScript RT reagent Kit with gDNA Eraser (TAKARA, Japan). The primers used for RT-PCR of *KCNJ15* were FP (5’-CACACTGCGTGACAAGTGAA-3’) and RP (5’-TACAGCCGGACCATACCTTC-3’). The amplified product was electrophoresed on a 1.5% agarose gel.

### Supplementary Figures

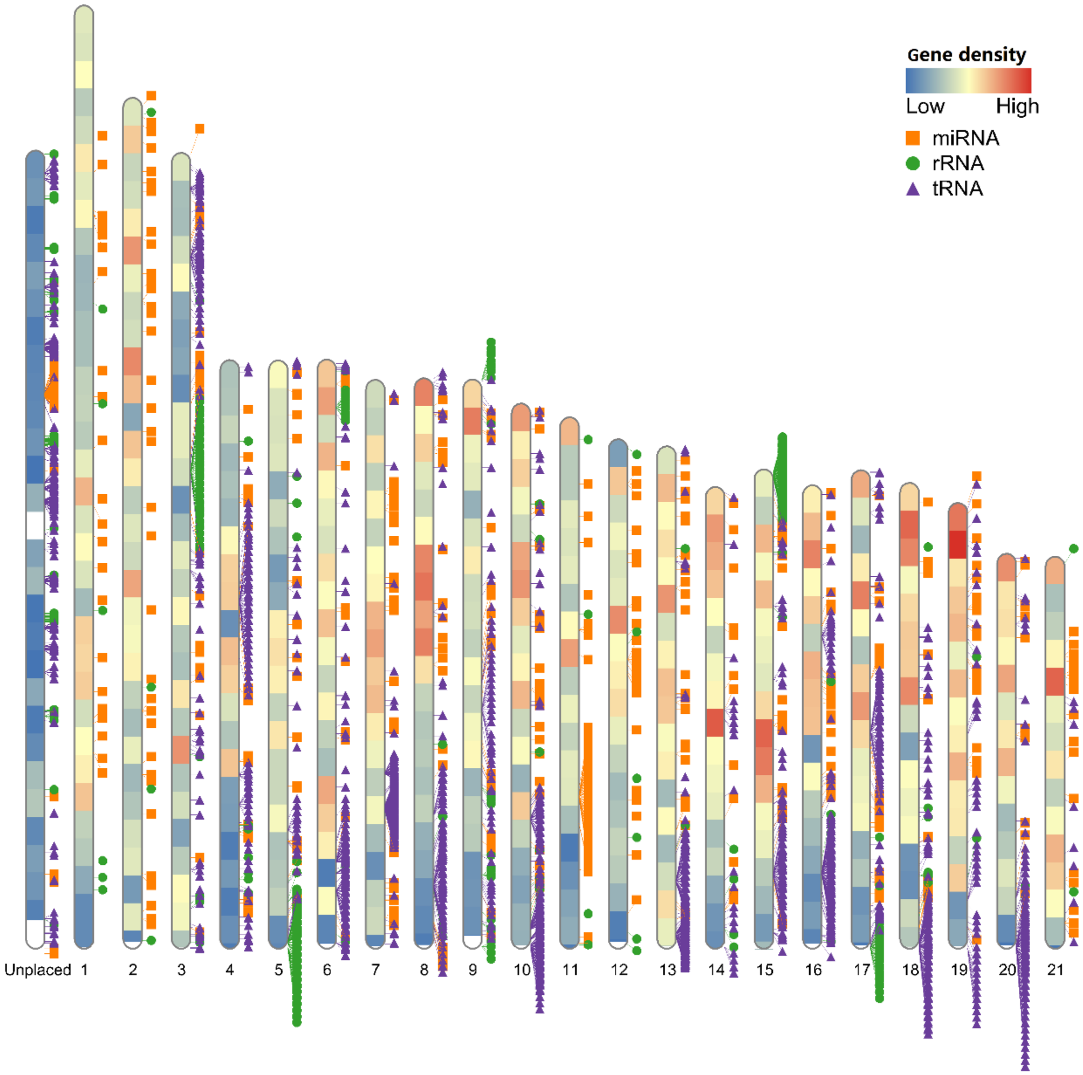


**Fig. S1. Genomic landscape of annotated protein-coding genes and non-coding RNA elements in the Siamese fighting fish genome.**

The color of the chromosome fragments corresponds to the gene density (shades of red represent higher density and shades of blue, lower density), and the miRNA, tRNA and rRNA are denoted as solid rectangles, circles and triangles, respectively. All the unplaced scaffolds are stacked together on the left "Unplaced" block. Note the gene hotspot close to the beginning of chromosome 19. All the elements are anchored into the genome using "RIdeogram" package (*54*)in the R program.

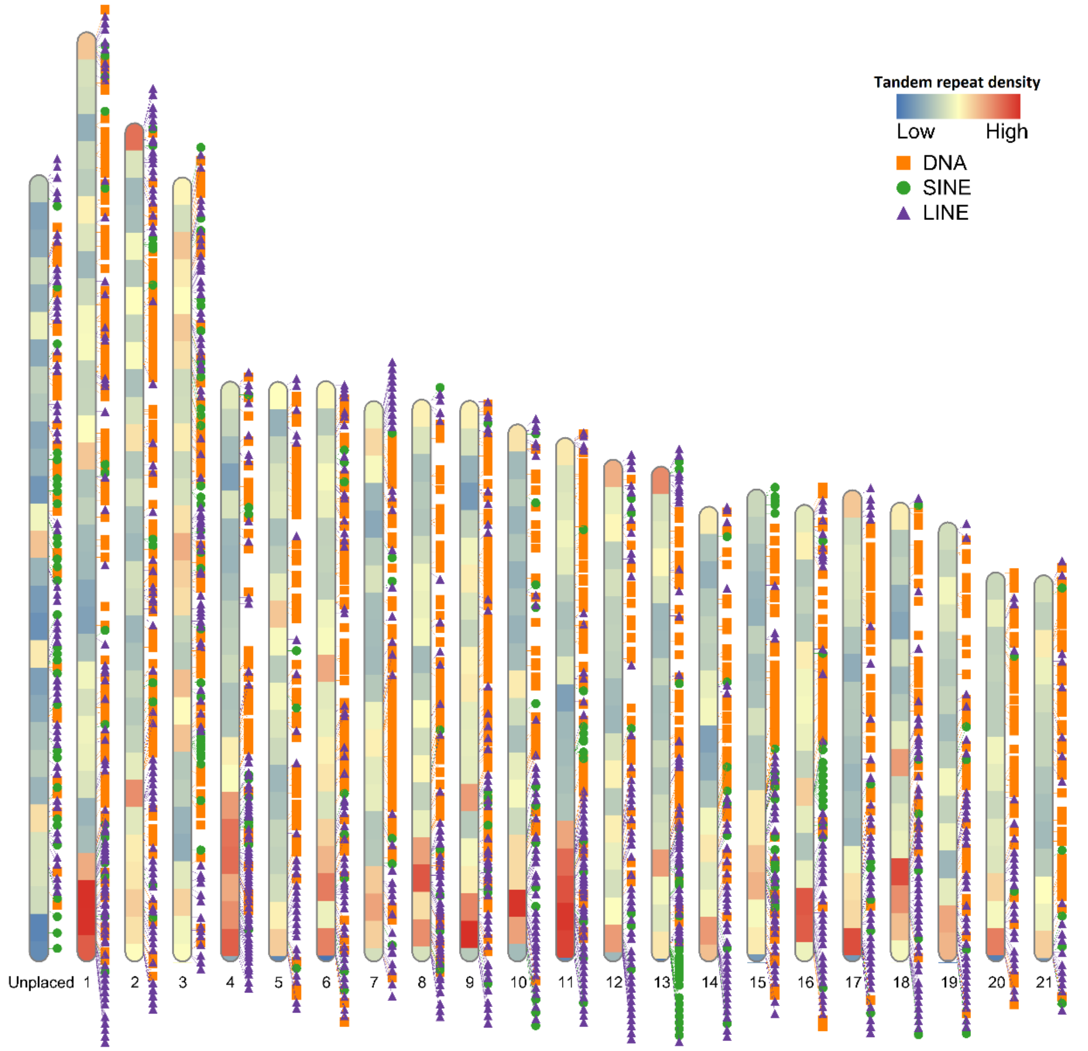


**Fig. S2. Genomic landscape of repeat sequences in the Siamese fighting fish genome.**

The color of the chromosome fragments corresponds to the repeat sequence density (shades of red represent higher density and shades of blue, lower density), and the DNA transposons, SINEs and LINEs are denoted as solid rectangles, circles and triangles, respectively. All the unplaced scaffolds are stacked together on the left "Unplaced" block. All the elements are anchored into the genome using the "RIdeogram" package (*54*) in the R program.

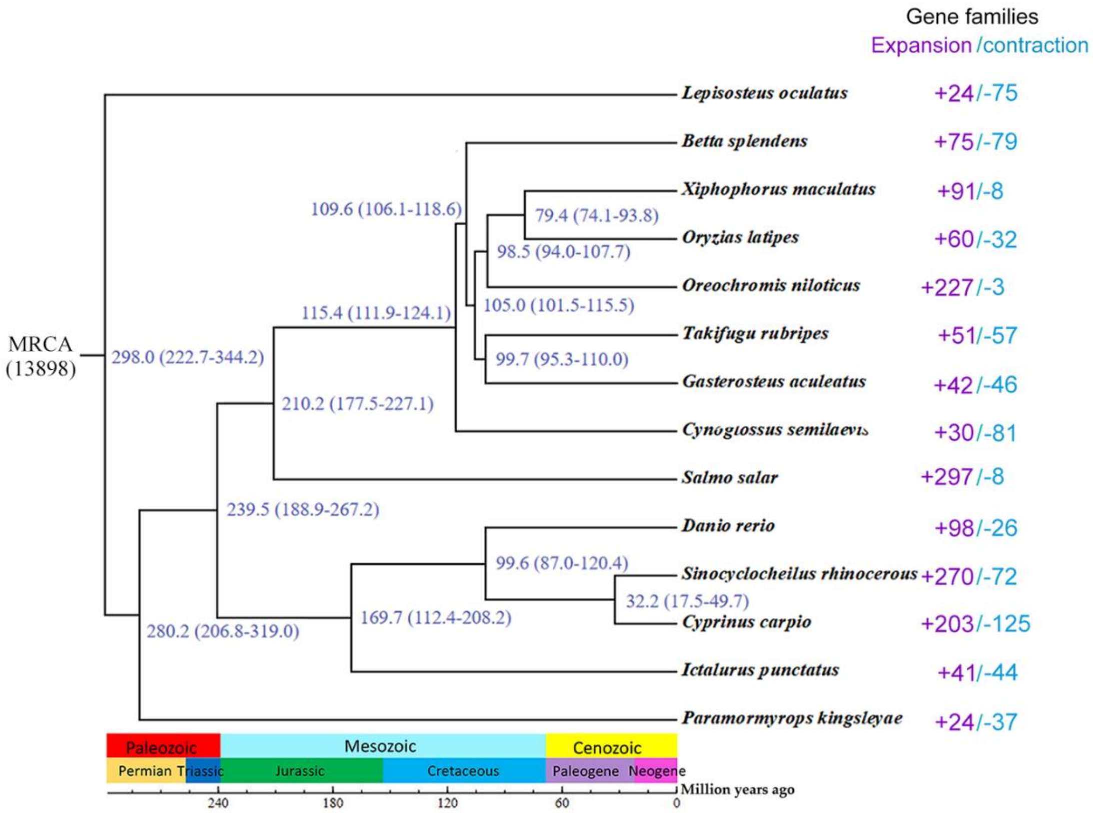


**Fig. S3. Time-calibrated phylogeny of teleosts.**

The phylogeny is constructed with genome wide single-copy orthologous genes from 14 teleost species including the Siamese fighting fish (*Betta splendens*). The numbers close to the nodes of the tree are the estimated divergence times (in units of millions of years ago). The numbers to the right of the phylogeny are the number of expanded and contracted gene families specific to each fish lineage (discussed in Supplementary Text 3).

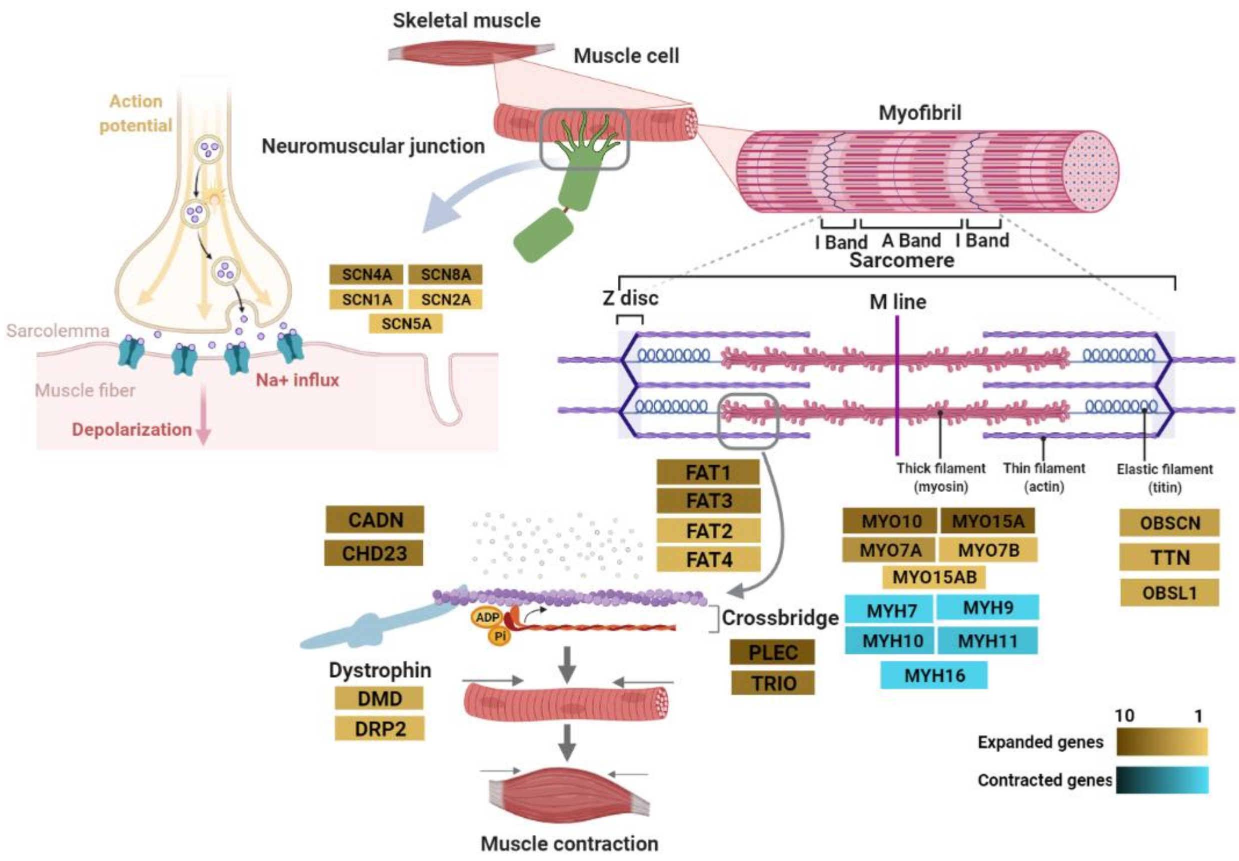


**Fig. S4. A schematic illustration presenting the expanded and contracted genes related to muscular function in the Siamese fighting fish.**

Gene families are labeled with a highlighted background with intensity ranging from 1 to 10 representing the number of copies of genes in each gene family. Expanded gene families are highlighted in yellowish-brown and contracted gene families are highlighted in skyblue. As discussed in Supplementary Text 3, several of the gene families expanded in the lineage of *Betta splendens* are related to the neuromuscular junction, muscle contraction and relaxation activities. These include proteins in the sarcomere (myosin encoded by the *MYO* genes, plectin encoded by *PLEC* and associated to Z-discs, titin encoded by *TTN* and obscurin encoded by *OBS* genes), cadherins (*FAT*, *CADN* and *CHD23*) involved in cell adhesion, *TRIO* involved in cytoskeleton reorganization, dystrophins (*DMD* and *DRP2*) and sodium channels encoded by *SCN* genes and related to the neuromuscular junction. Additionally, one of the family of genes encoding myosin heavy chains (*MYH*) is contracted in the *B. splendens* lineage.

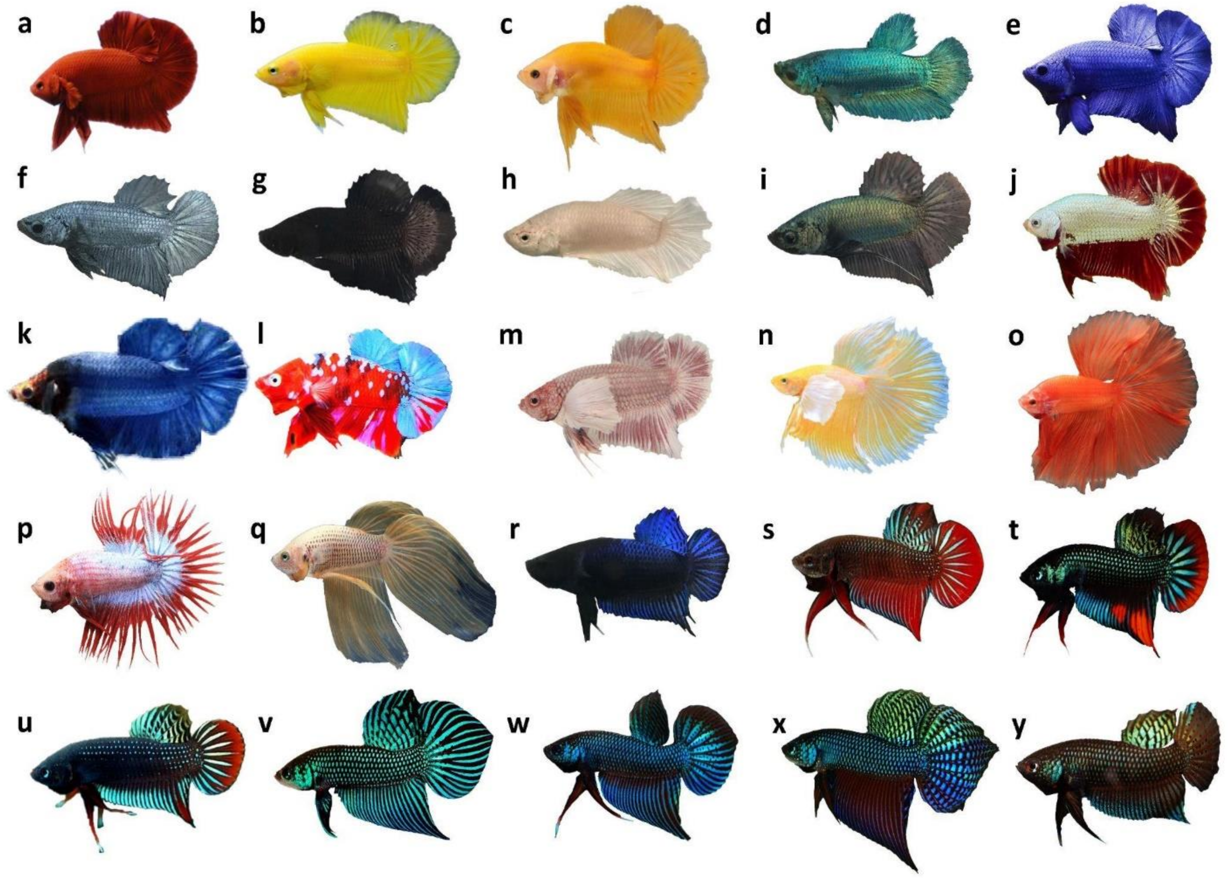


**Fig. S5. Illustration of samples used in this study.**

(a-r) are different domesticated breeds of the Siamese fighting fish and (s-y) are different wild Betta species. a: Solid Red Halfmoon Plakat (HMPK); b: Solid Yellow HMPK; c: Solid Orange HMPK; d: Turquoise-green HMPK; e: Royal-blue HMPK; f: Steel-blue HMPK; g: Solid Black HMPK; h: Solid Opaque HMPK; i: Copper HMPK; j: Dragon HMPK; k: Giant HMPK; l: Koi HMPK; m: HMPK Dumbo; n: Halfmoon Dumbo; o: Halfmoon; p: Crowntail; q: Veiltail; r: Fighter; s: wild *Betta splendens*; t: *Betta imbellis*; u: *Betta siamorientalis*; v: *Betta mahachaiensis*; w: *Betta smaragdina*; x: *Betta smaragdina guitar*; y: *Betta stiktos.*

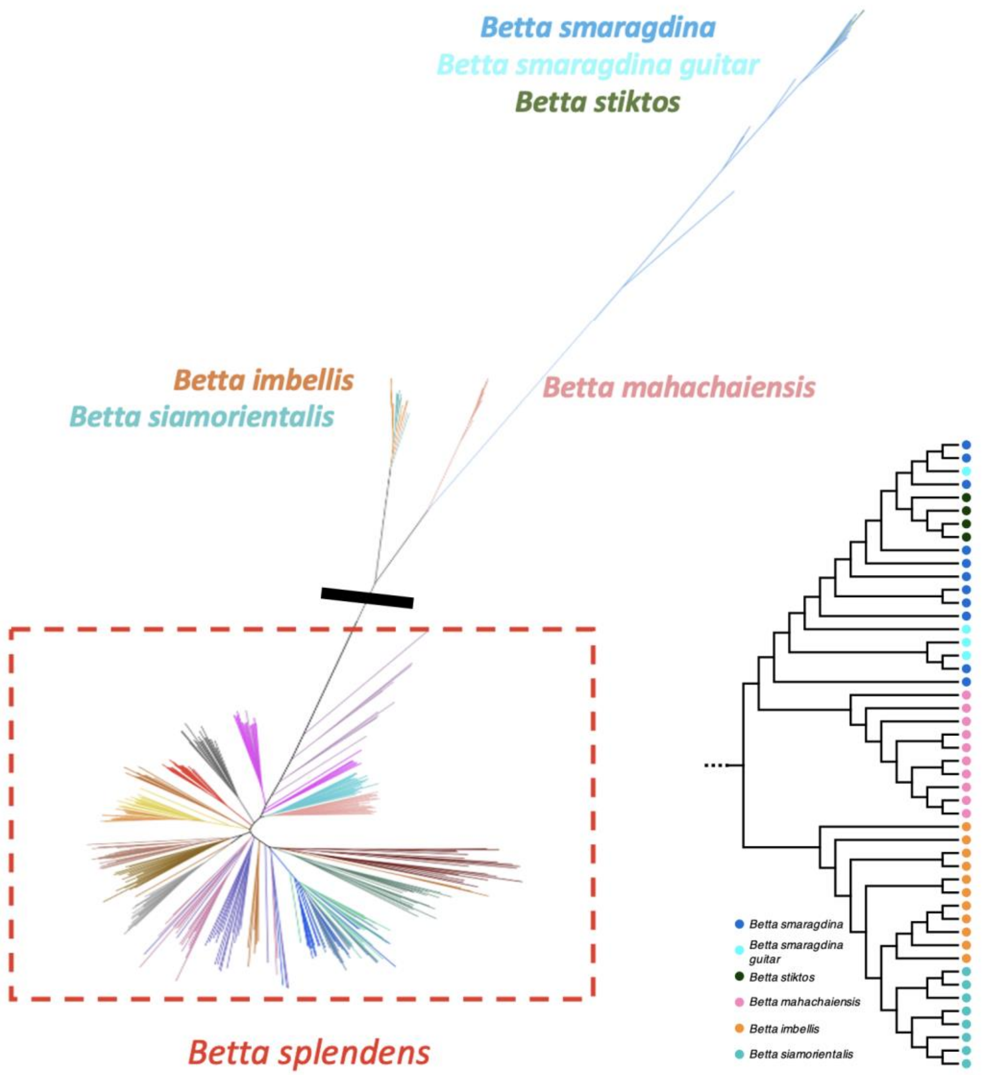


**Fig. S6. Neighbor-joining tree of the Betta fish.**

The tree is constructed using concatenated sequences at 11.90 M genome-wide SNP sites. The red box highlights the *Betta splendens* clade which is presented in detail in Fig. 1B. The inset to the bottom right is the subtree containing other wild species, which show that *B. imbellis* and *B. siamorientalis* cluster together in the tree, and the lineages of *B. stiktos* and *B. smaragdina guitar* are interspersed in the tree. The black line on the left NJ tree shows where the subtree was truncated.

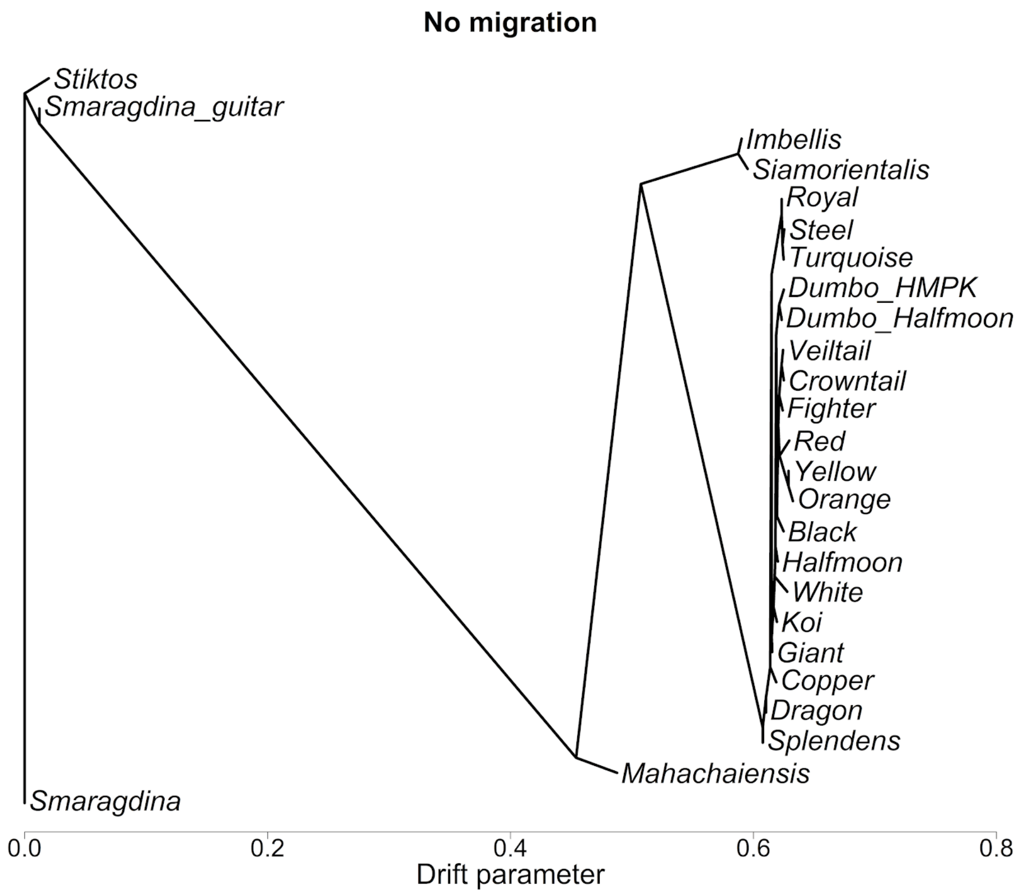


**Fig. S7. Treemix plot of Betta fish without migration.**

Similar to the Neighbor-joining tree, all domesticated *Betta splendens* breeds cluster together in Treemix, and domesticated HMPK breeds cluster internally by general morphology (*e.g.,* blue and green cluster together, as well as red, orange and yellow breeds).

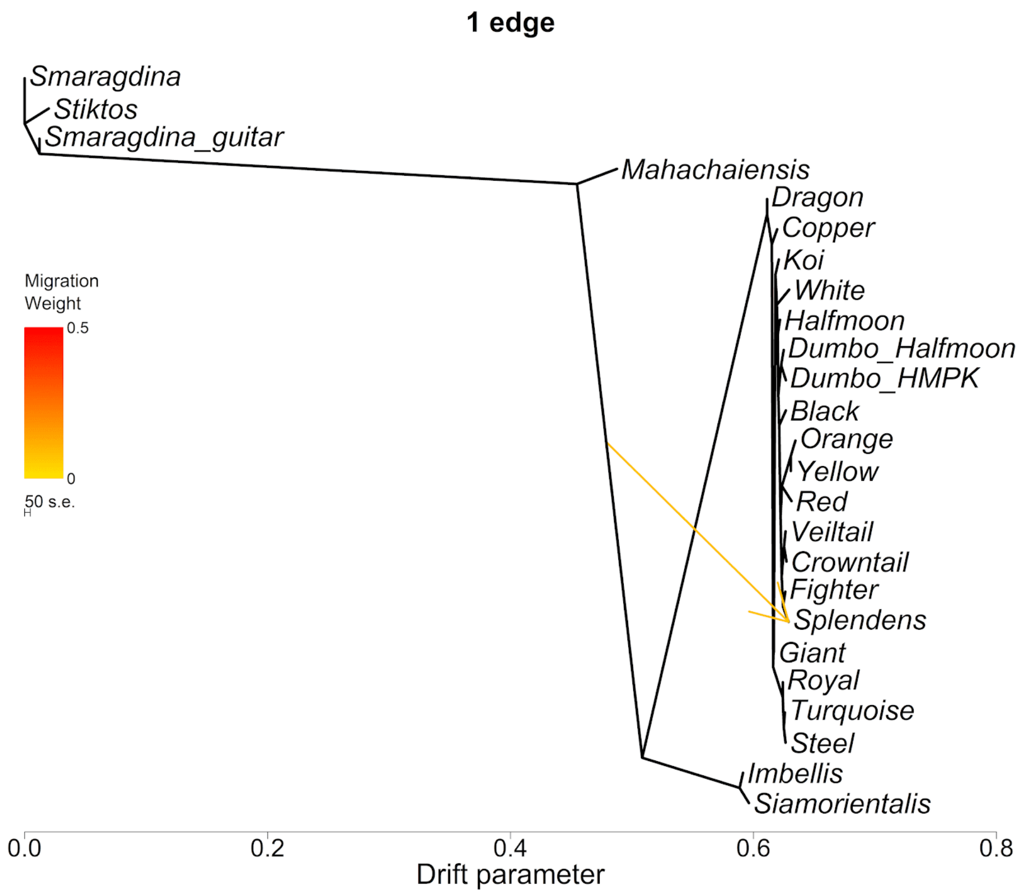


**Fig. S8. Treemix plot of Betta fish with one migration edge allowed.**

With one migration edge allowed, Treemix fits an edge connecting the ancestor of *B. imbellis*, *B. siamorientalis* and *B. splendens* to the wild *B. splendens*.

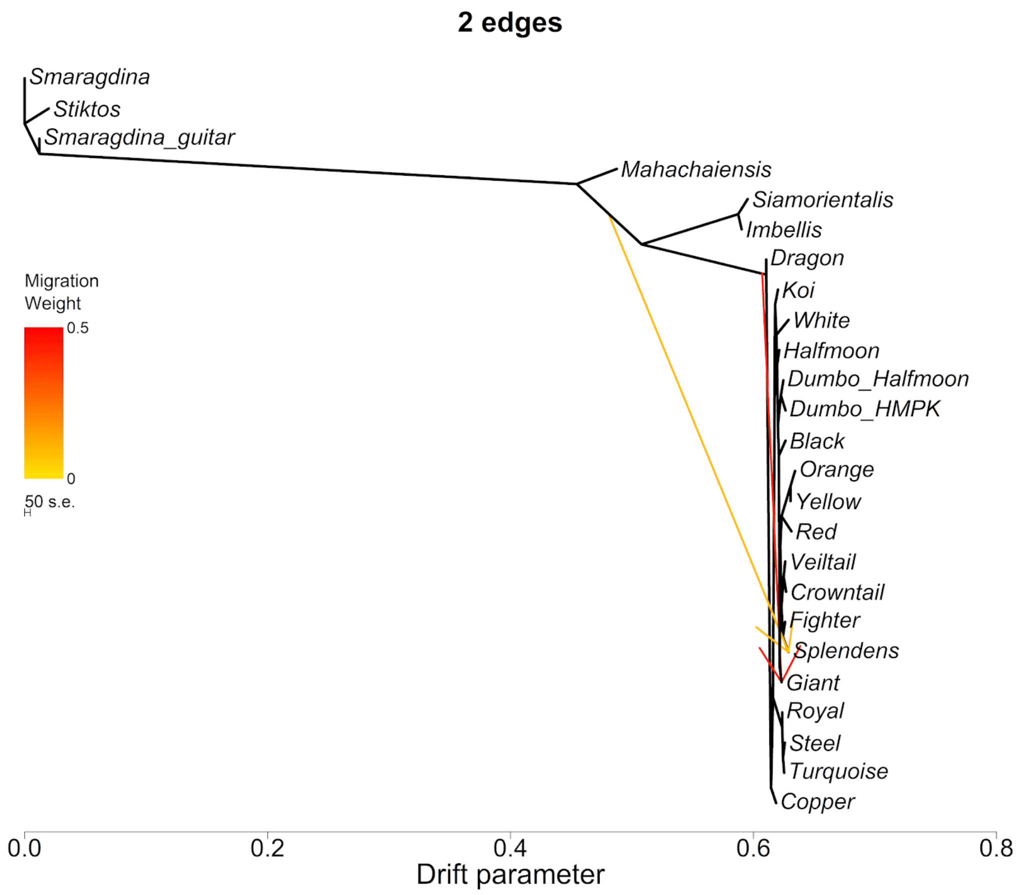


**Fig. S9. Treemix plot of Betta fish with two migration edges.**

With two migration edges allowed, Treemix fits the same edge as with one edge allowed, as well as a migration edge from the base of the *B. splendens* clade into the Giant breed.

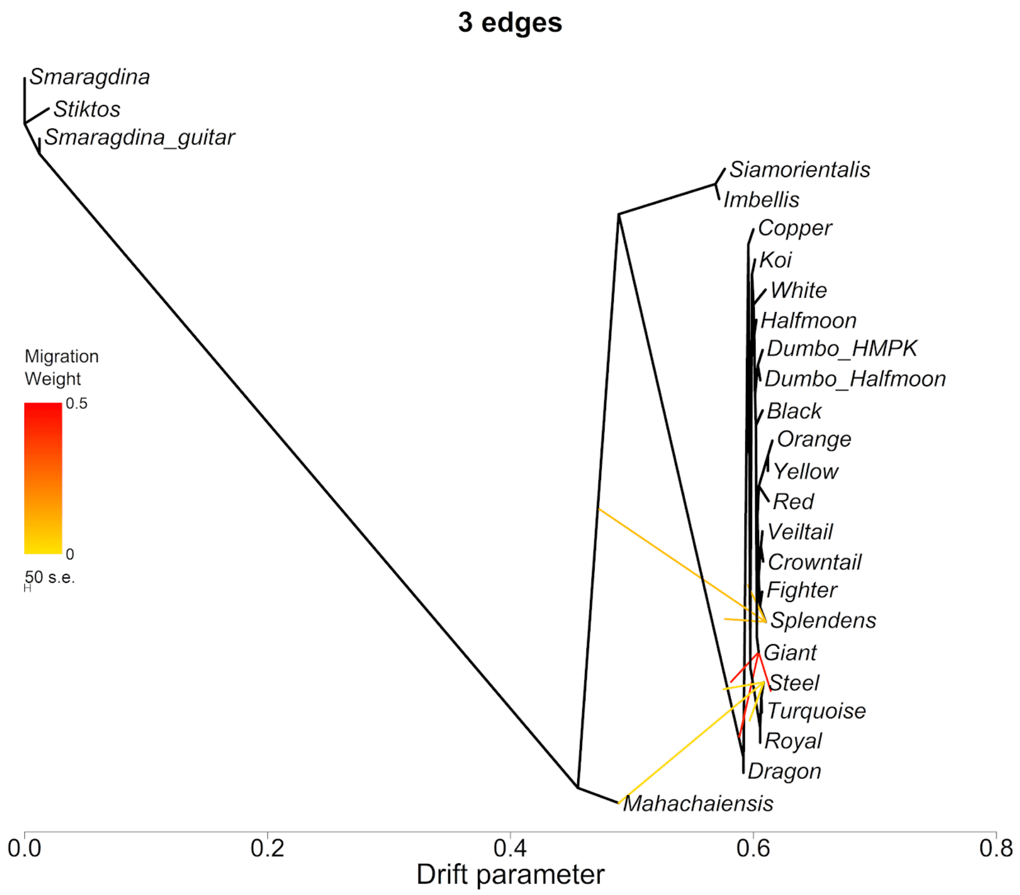


**Fig. S10. Treemix plot of Betta fish with three migration edges.**

With three migration edges allowed, Treemix fits the same edges as with two edges allied, as well as one edge from *B. mahachaiensis* into the Steel blue breed.

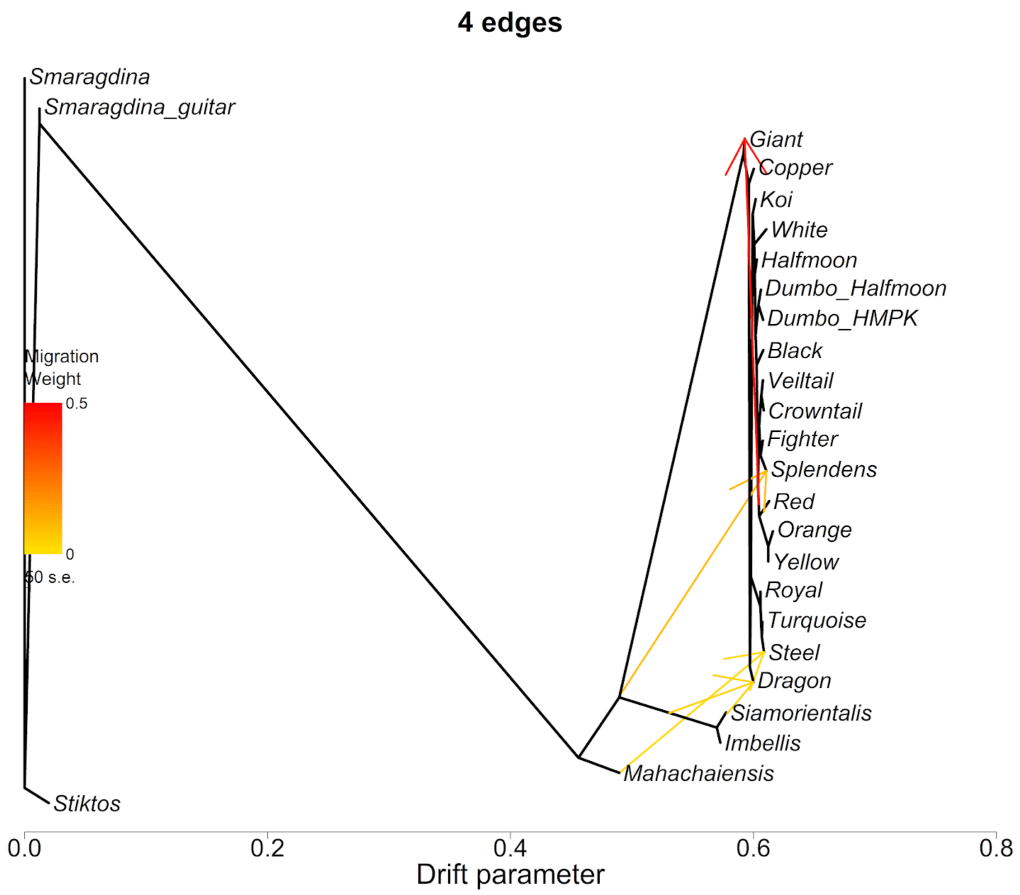


**Fig. S11. Treemix plot of Betta fish with four migration edges.**

With four migration edges allowed, Treemix fits two of the same edges with 3 edges allowed, as well as one new migration edge from the ancestor of *B. siamorientalis* and *B. imbellis* into the Dragon breed, as well as a heavy edge from the ancestor of Red, Orange and Yellow breeds into Giant.

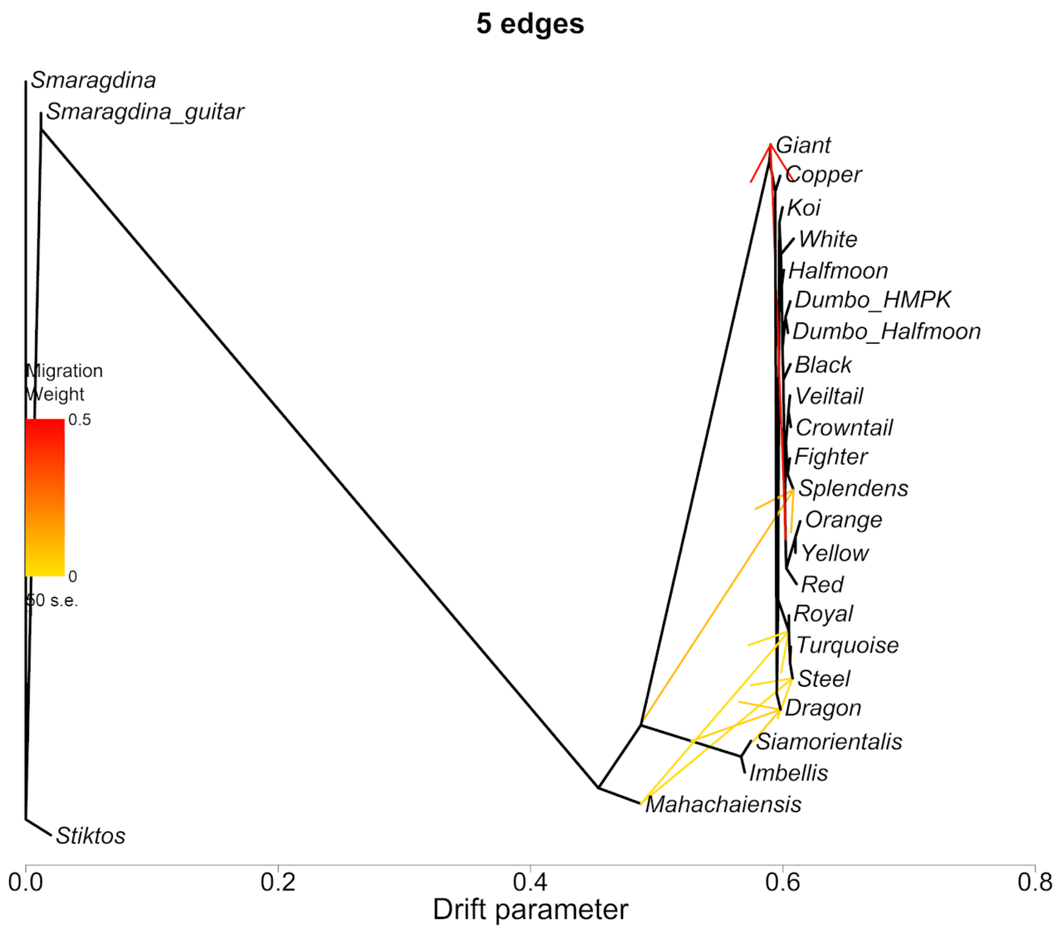


**Fig. S12. Treemix plot for Betta fish with five migration edges.**

With five migration edges allowed, Treemix fits an additional edge (relative to the model with four migrations allowed) from *B. mahachaiensis* into the blue-green clade.

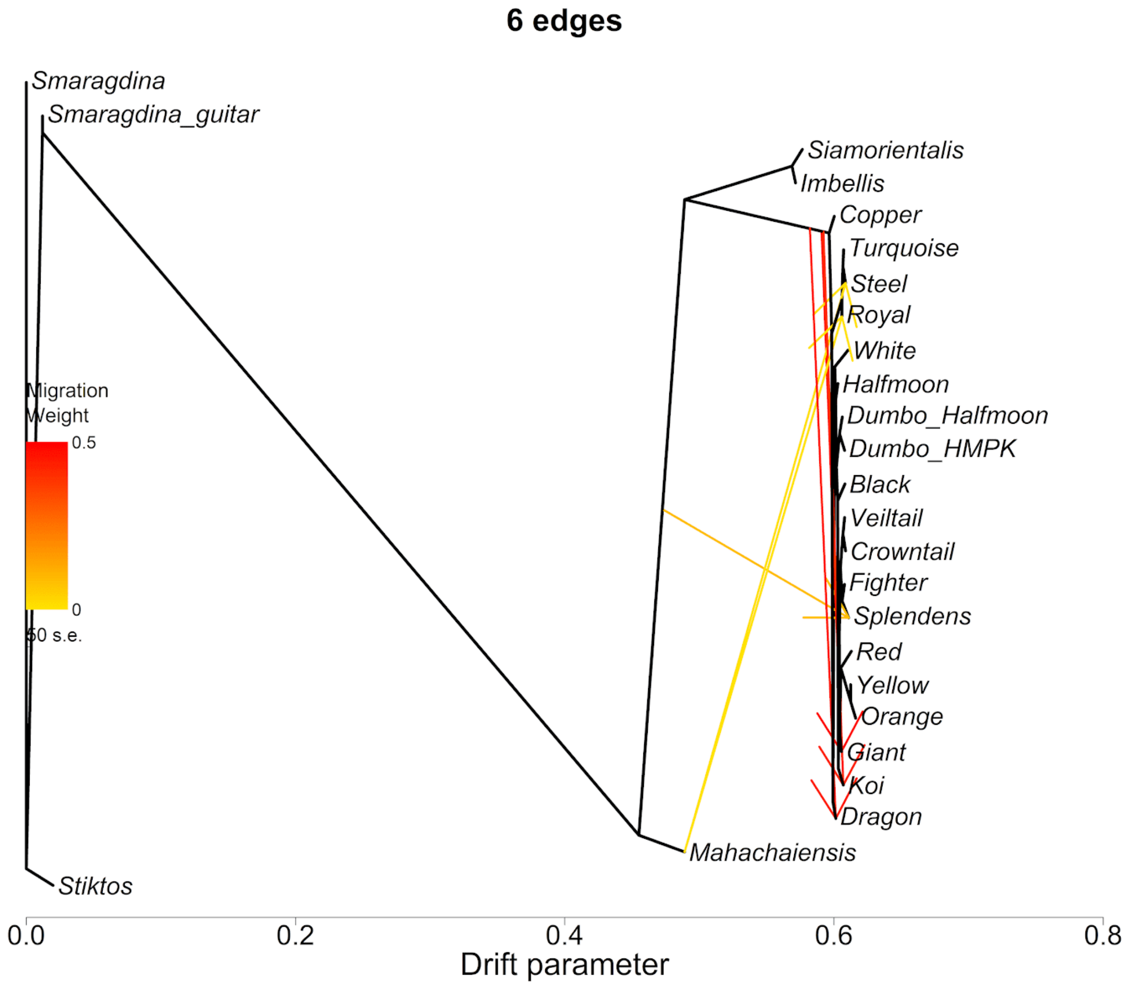


**Fig. S13. Treemix plot for Betta fish with six migration edges.**

With six migration edges allowed, Treemix fits additional edges (relative to the model with five migrations allowed) with high weight from the ancestor of *B. splendens* into the Giant, Koi and Dragon breeds.

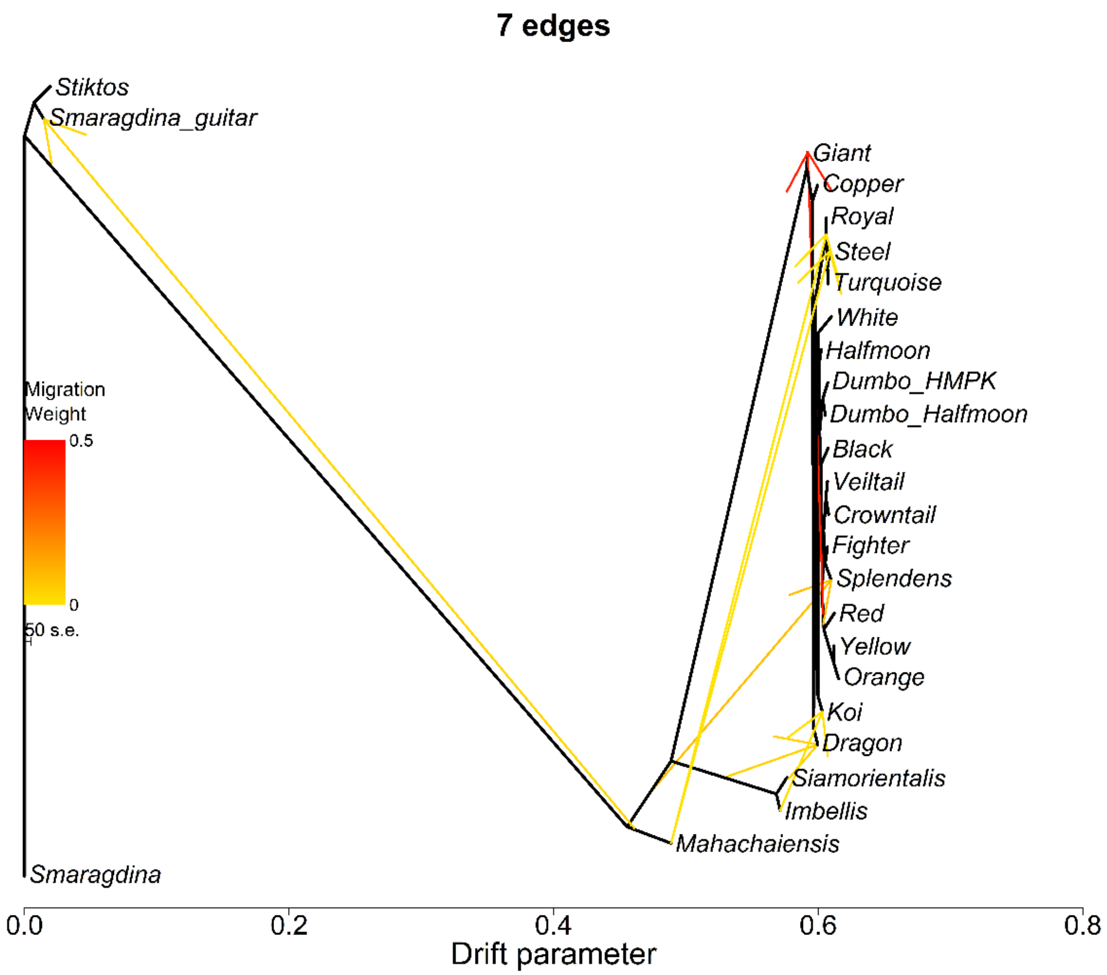


**Fig. S14. Treemix plot for Betta fish with seven migration edges allowed.**

With seven migration edges allowed, Treemix consistently (relative to models with fewer migration edges) fits migration edges from *B. mahachaiensis* into the blue-green clade, as well as an edge from the ancestor of *B. siamorientalis* and *B. imbellis* into Dragon. A heavy edge between the ancestors of Red, Yellow and Orange and Giant is also observed.

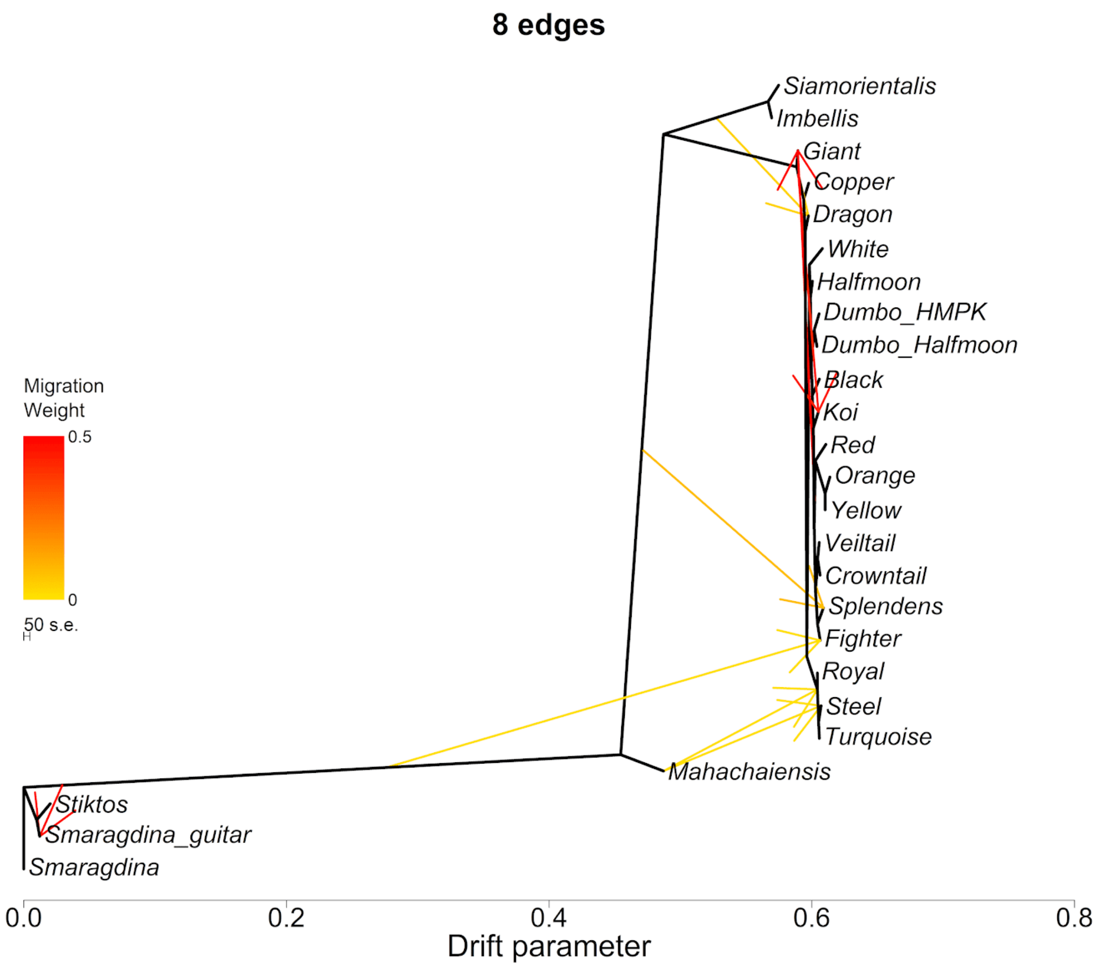


**Fig. S15. Treemix plot for Betta fish with eight migration edges allowed.**

With eight migration edges allowed, Treemix consistently (relative to models with fewer migration edges) fits migration edges from *B. mahachaiensis* into the blue-green clade, as well as an edge from the ancestor of *B. siamorientalis* and *B. imbellis* into Dragon. A heavy edge between the ancestors of Red, Yellow and Orange and Giant is also observed.

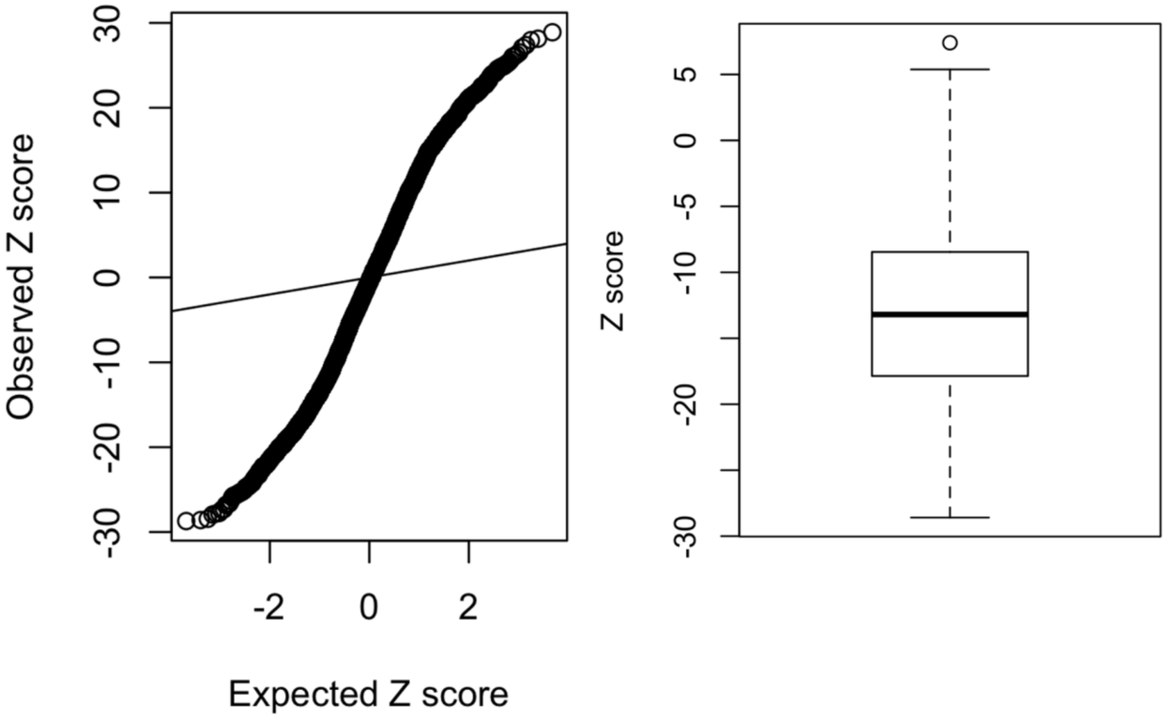


**Fig. S16. ABBA-BABA test for introgressions between *B. mahachaiensis* and domesticated *B. splendens***

The left panel shows the QQ plot testing D(dom, dom; mahachaiensis, outgroup) where the outgroup is a *B. smaragdina* individual. At H1 and H2 position, individuals of all different breeds were tested iteratively. For each breed or species, three randomly chosen individuals were used. The right panel shows the Z scores for D-statistic tests in the form of D(Turquoise/Royal/Copper/Steel, other breeds; mahachaiensis, outgroup). Most of the tests have negative value supporting gene flow between *B. mahachaiensis* and the Turquoise green, Royal blue, Copper or Steel blue breeds.

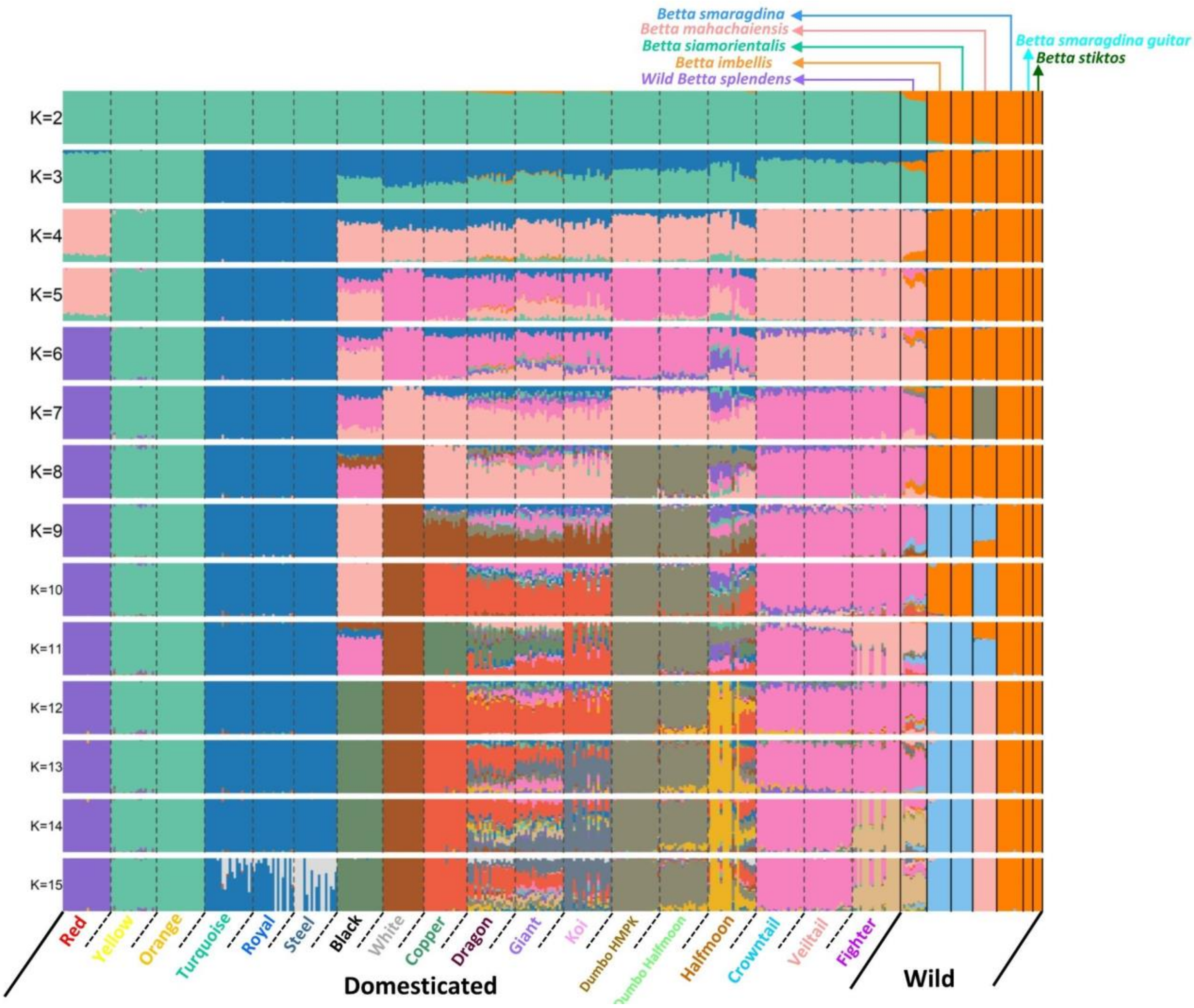


**Fig. S17.** ***ADMIXTURE* analysis varying *K* from 2 to 15**

The analysis was performed on the 721K SNP dataset containing 410 individuals. Individuals of the same domesticated breeds that are phenotypically defined by color and fin morphology (listed on the x axis) generally cluster together.

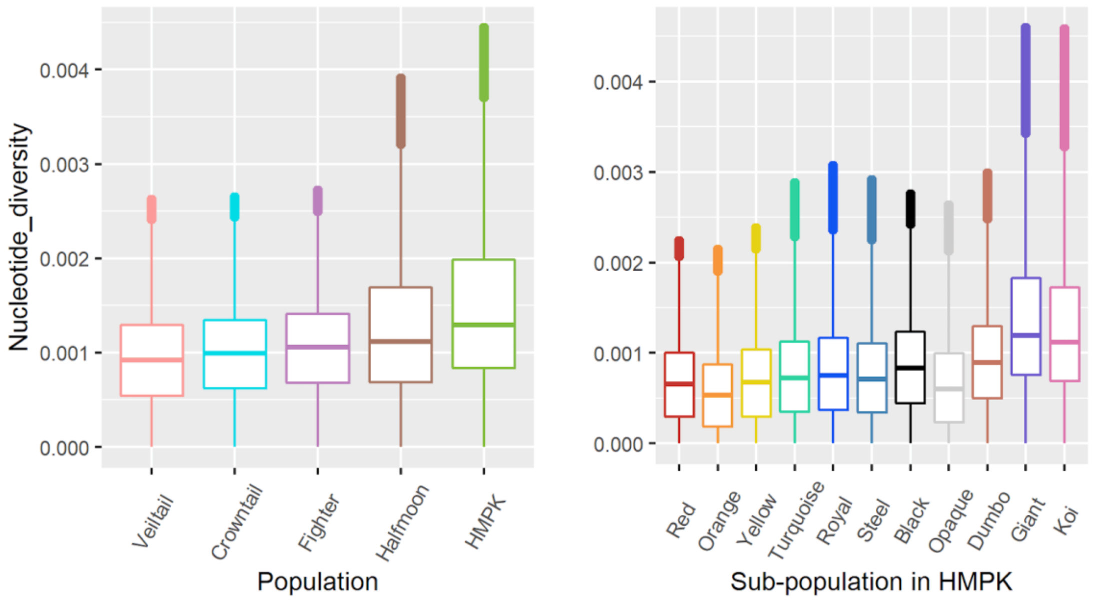


**Fig. S18. Nucleotide diversity across breeds of domesticated *Betta splendens*.**

Low nucleotide diversity in the domesticated breeds can be explained by bottlenecks during domestication. Among the HMPK breeds, breeds selected for color tend to have lower genetic diversity.

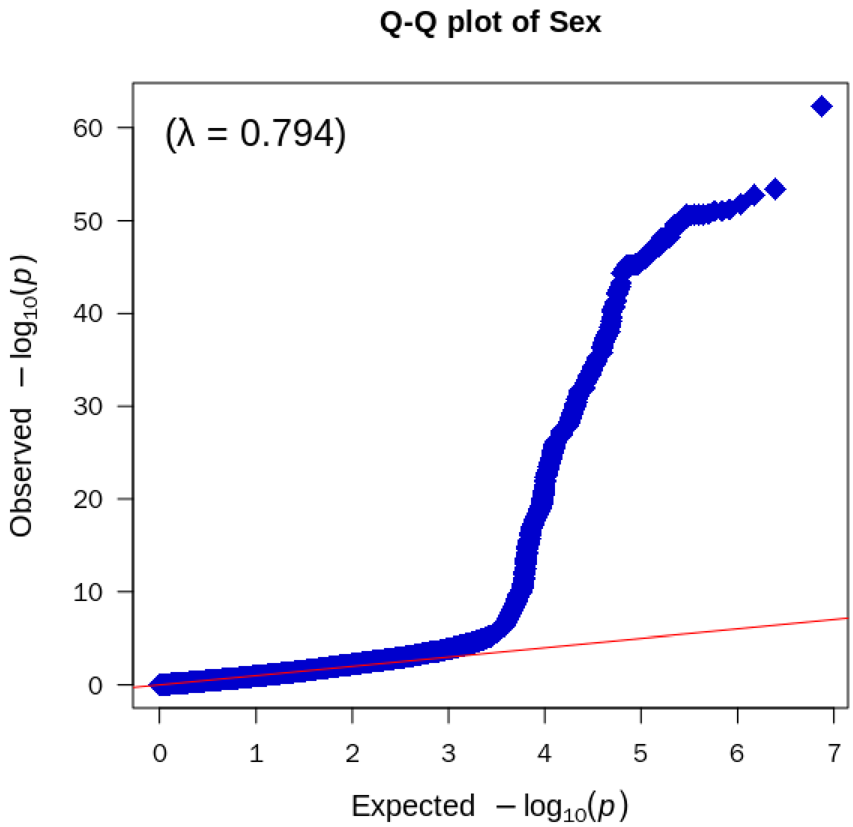


**Fig. S19. QQ plot for sex GWAS.**

QQ plot for the GWAS for sex described in main Fig. 2A. 590 males and 137 females were included. Red line shows expected P-values and blue dots show observed P-values.

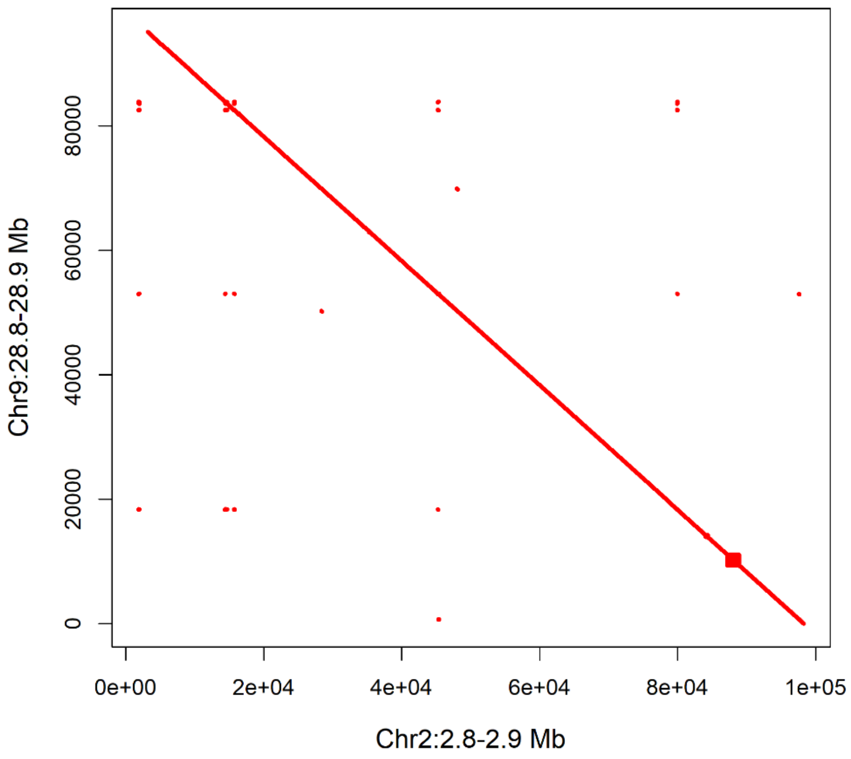


**Fig. S20. Synteny at the sex determining locus containing *DMRT1*.**

The plot is a comparison between our genome assembly of a female (x axis) of the Siamese fighting fish genome and a previously published assembly of a male (fBetSpl5.3, GenBank accession: GCA_003650155.3).

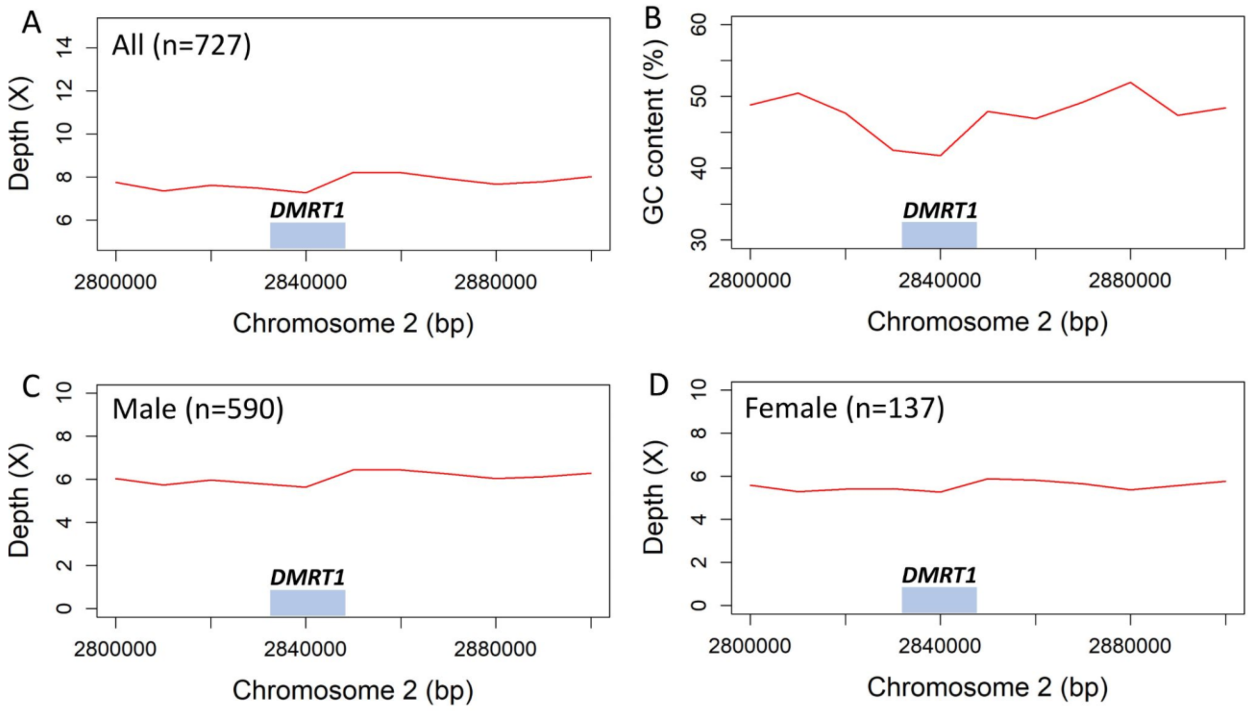


**Fig. S21. Sequencing depth and GC content in the sex determining region in the vicinity of the *DMRT1* gene of the Siamese fighting fish in this study.**

No differences in sequencing depth among males and females is observed, suggesting that sex chromosomes are not differentiated in this species.

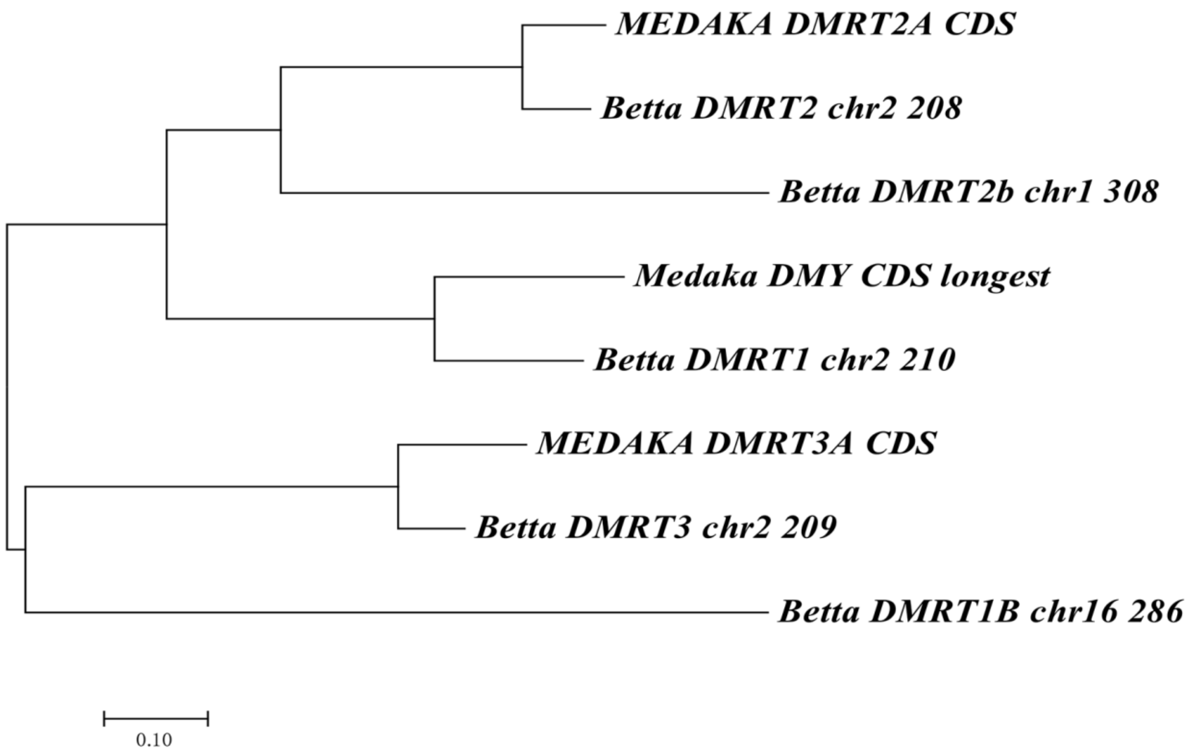


**Fig. S22. Neighbor-joining tree of putative Betta and Medaka sex determining genes.**

The Betta sex determining gene identified in this study (*DMRT1*) is most closely related to the Medaka *DMY* gene.

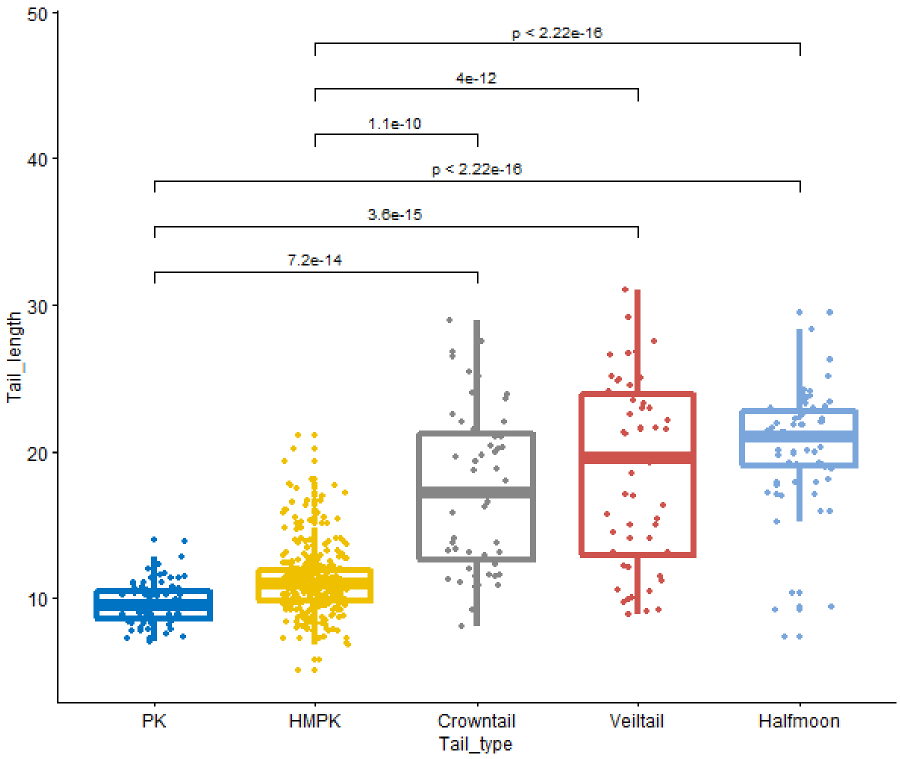


**Fig. S23. Caudal fin length in five breeds of the Siamese fighting fish.**

Caudal fins of the long-finned breeds (Crowntail, Veiltail and Halfmoon) are significantly longer than those of Plakat (PK) and Halfmoon Plakat (HMPK) breeds (p-values for a Wilcoxon test are shown).

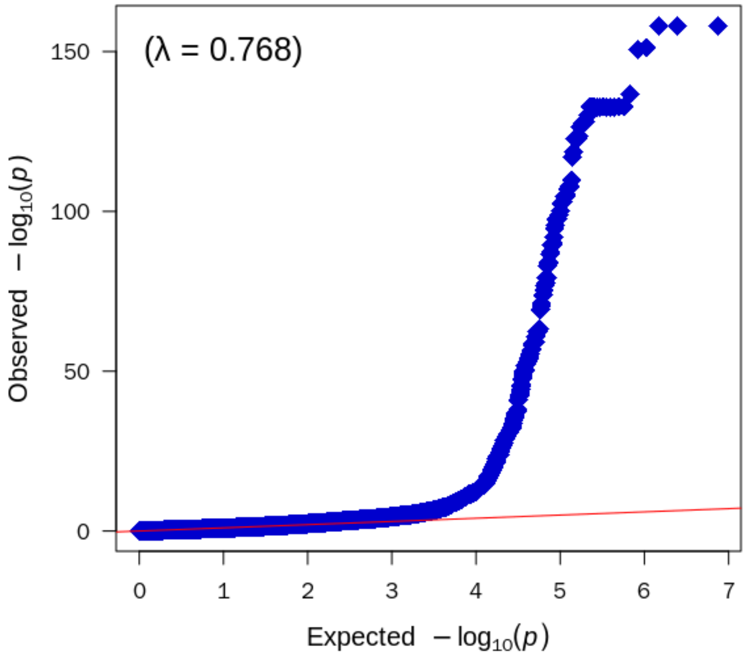


**Fig. S24. QQ plot for long vs. short fin GWAS.**

QQ plot for the GWAS for short-fin vs. long-fin described in main Fig. 3A. 525 short-fin individuals and 201 long-fin were included. Red line shows expected *P*-values and blue dots show observed *P*-values.

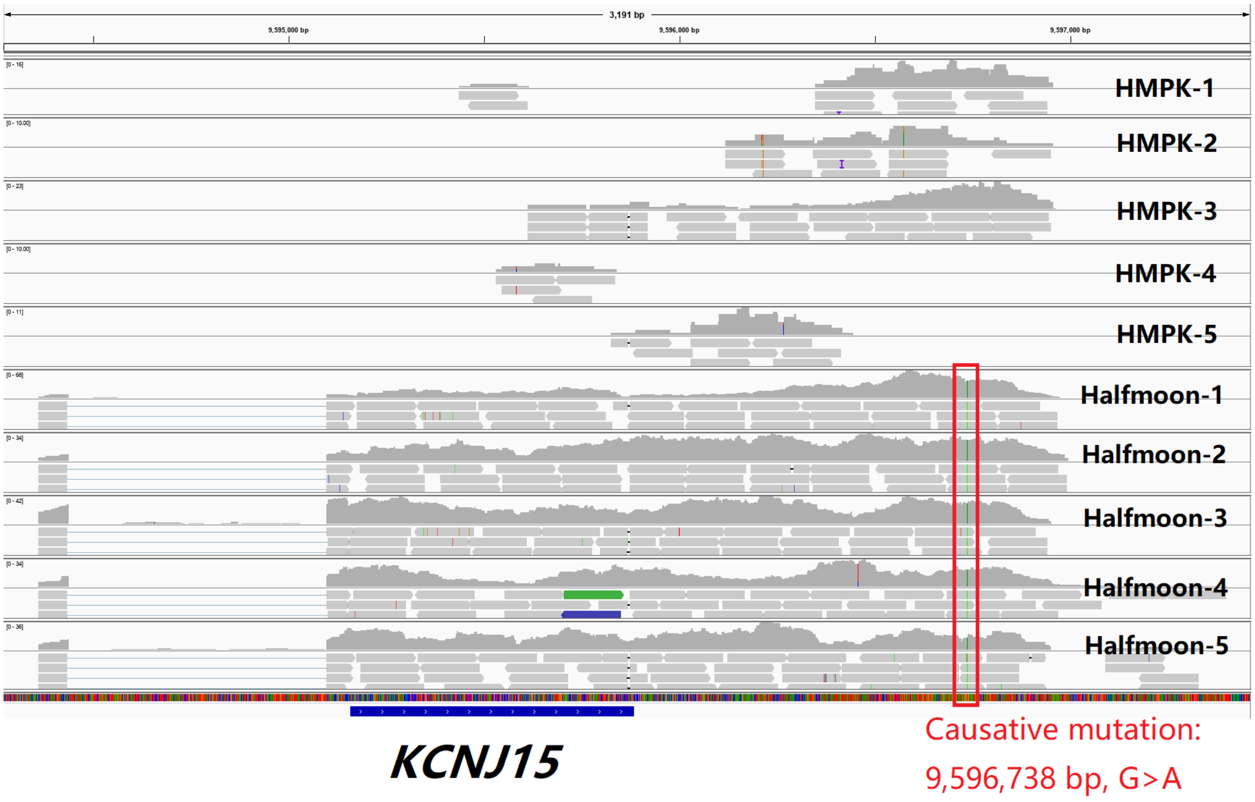


**Fig. S25. RNA-Seq reads mapped to the *KCNJ15* gene region in short-fin (HMPK) and long-fin (Halfmoon) fishes.**

Visualization of the alignment was done in IGV(*55*). The SNP with the strongest association to fin length (chr17:9,596,738, G>A) is highlighted in the red box.

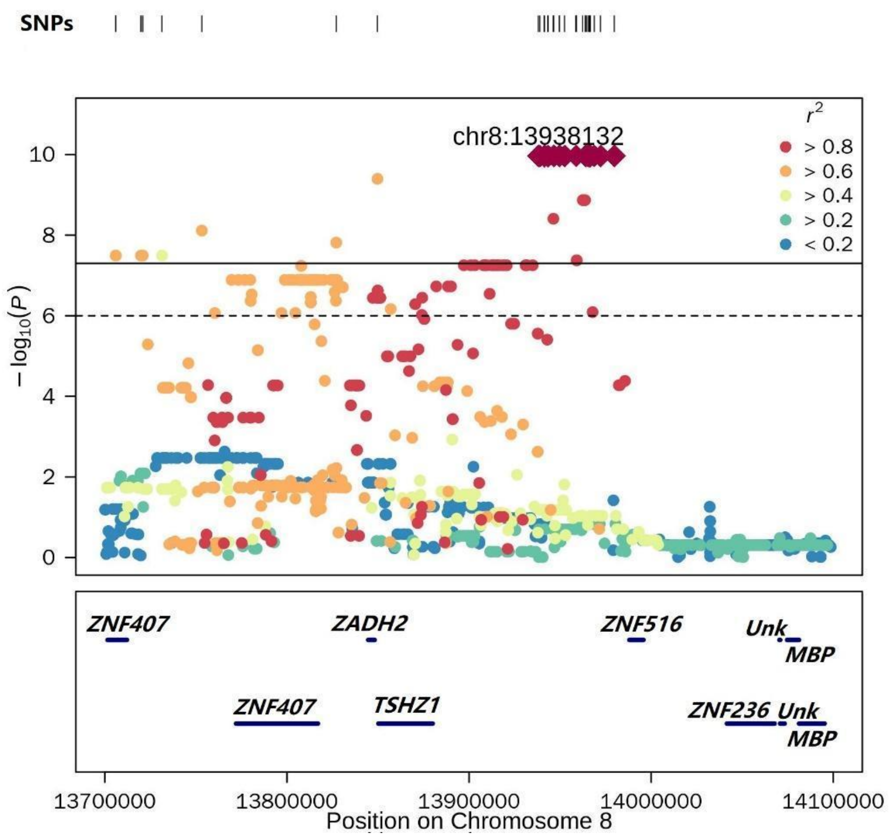


**Fig. S26. Regional plot of the locus with the strongest association in a GWAS between Veiltail and Halfmoon breeds.**

The solid and dashed lines represent genome-wide (0.05/Nsnp) and chromosomal level (1/Nsnp) significance thresholds, respectively. Colors represent linkage disequilibrium with the top variant, indicated by a diamond symbol. Bottom box shows the location of annotated genes in this region.

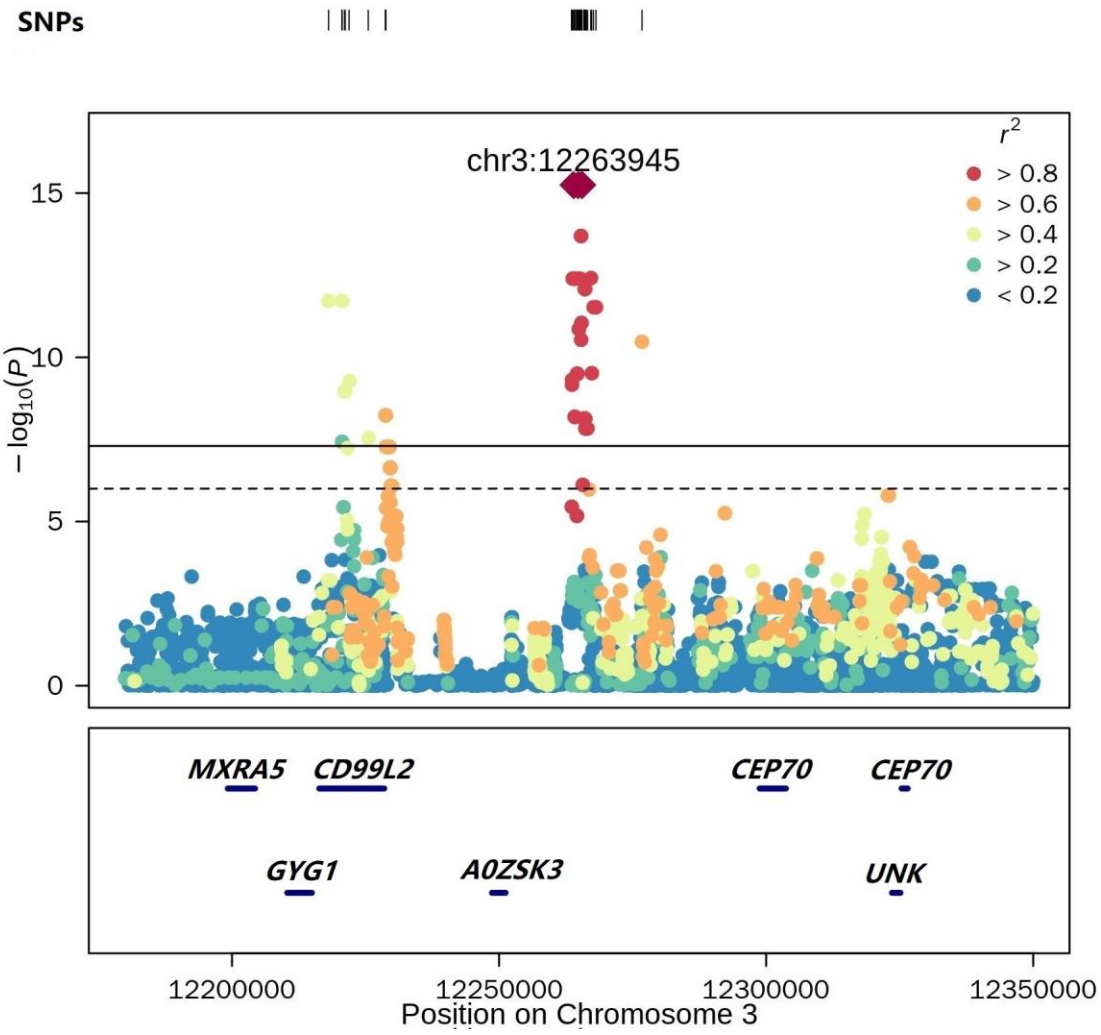


**Fig. S27. Regional plot of the locus with the strongest association in a GWAS between Crowntail and Halfmoon breeds.**

The solid and dashed lines represent genome-wide (0.05/Nsnp) and chromosomal level (1/Nsnp) significance thresholds, respectively. Colors represent linkage disequilibrium with the top variant, indicated by a diamond symbol. Bottom box shows the location of annotated genes in this region.

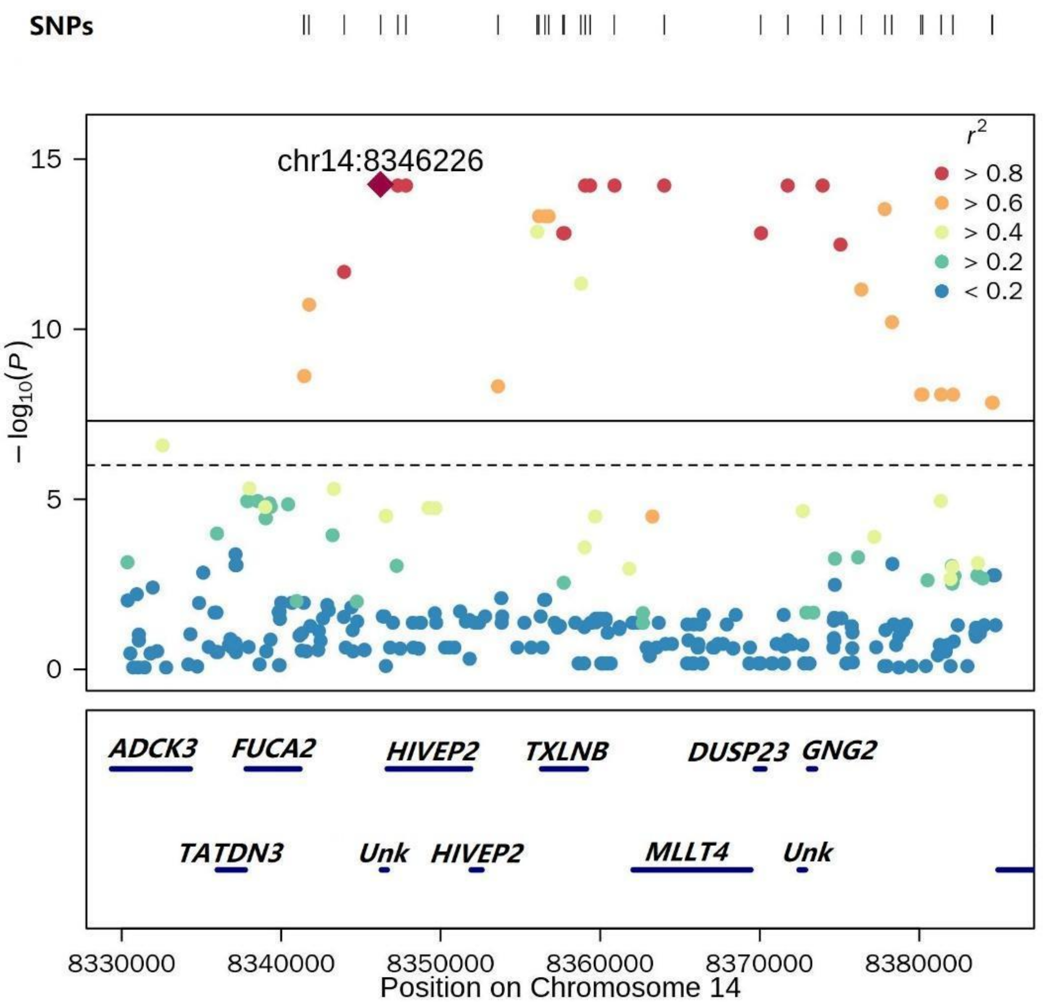


**Fig. S28. Regional plot of the locus with the strongest association in a GWAS between Veiltail and Crowntail breeds.**

The solid and dashed lines in the Manhattan plots represent genome-wide (0.05/Nsnp) and chromosomal level (1/Nsnp) significance thresholds, respectively. Colors represent linkage disequilibrium with the top variant, indicated by a diamond symbol. Bottom box shows the location of annotated genes in this region.

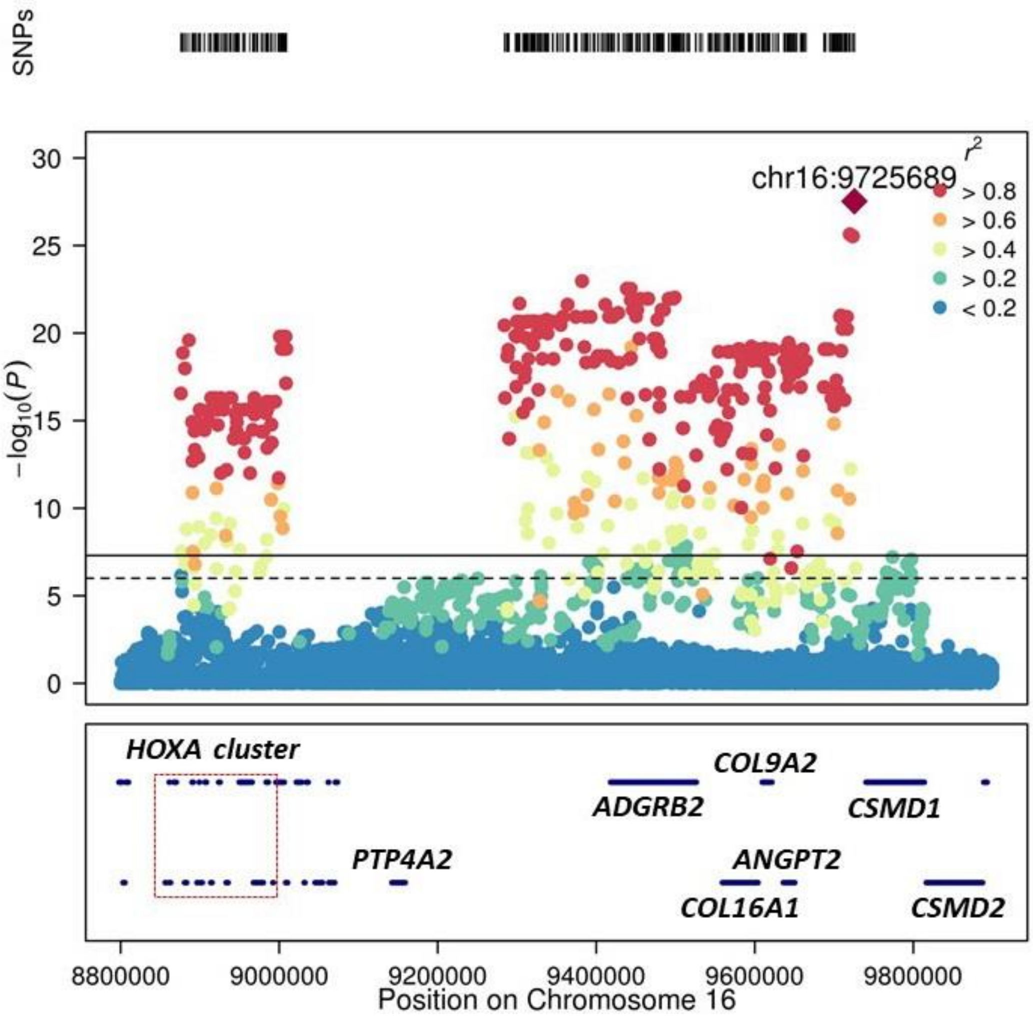


**Fig. S29. Regional plot of the region on chromosome 16 associated with the Dumbo phenotype.**

The solid and dashed lines in the top box represent genome-wide (0.05/Nsnp) and chromosomal level (1/Nsnp) significance thresholds, respectively. Colors represent linkage disequilibrium with the top variant, indicated by a diamond symbol. Bottom box shows the location of annotated genes in this region. Red dotted box shows the *HOXA* gene cluster.

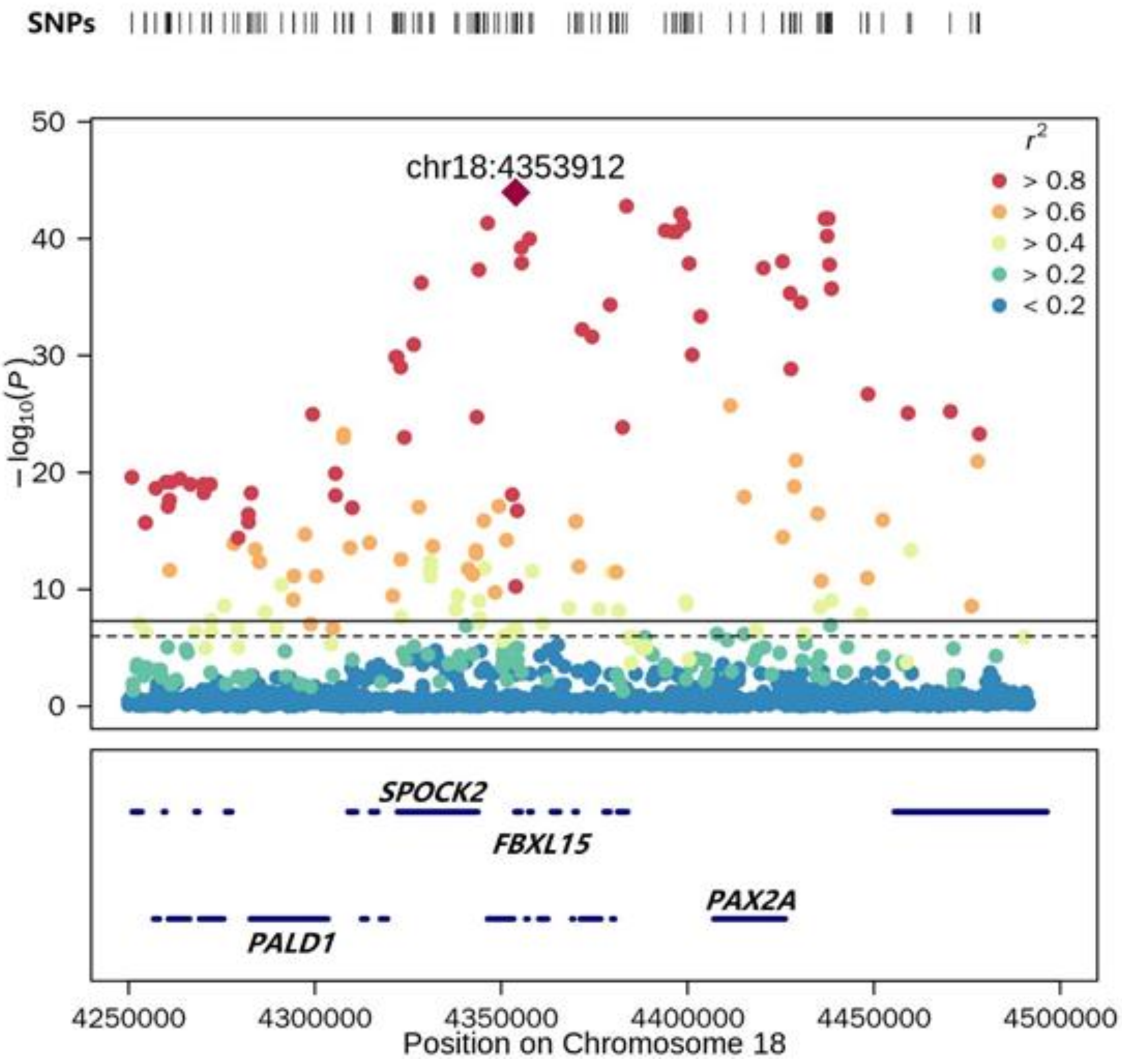


**Fig. S30. Regional plot of the region on chromosome 18 associated with the Dumbo phenotype.**

The solid and dashed lines in the top box represent genome-wide (0.05/Nsnp) and chromosomal level (1/Nsnp) significance thresholds, respectively. Colors represent linkage disequilibrium with the top variant, indicated by a diamond symbol. Bottom box shows the location of annotated genes in this region.

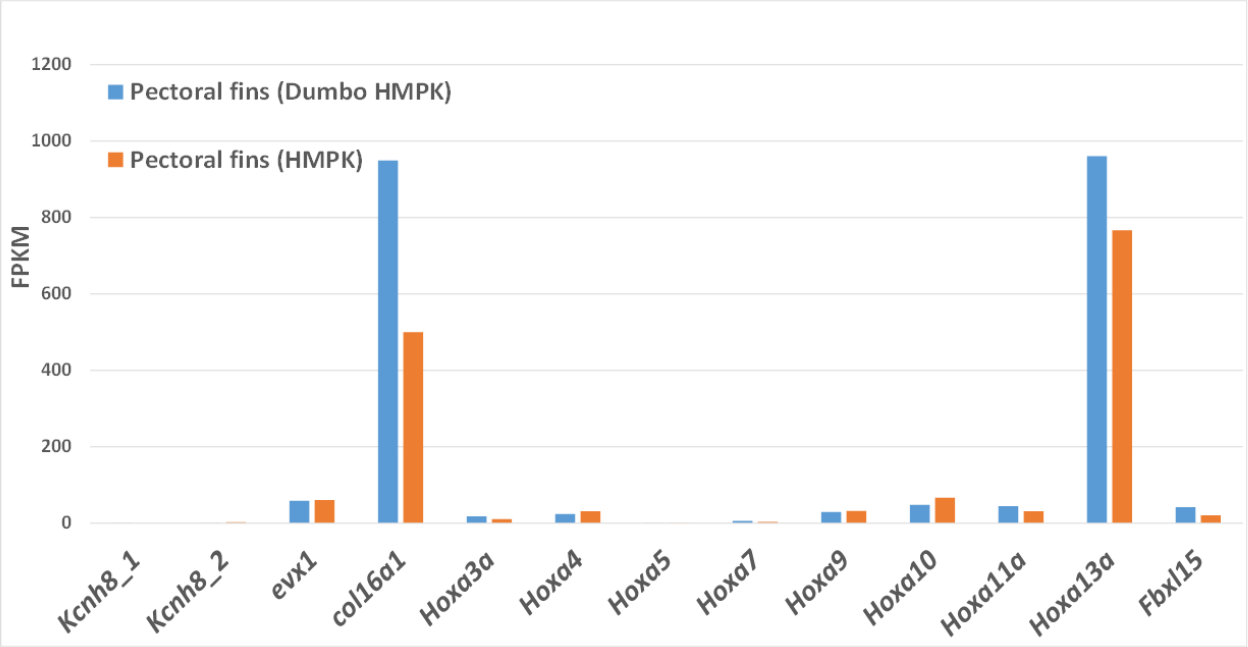


**Fig. S31. Expression profile of genes in the locus associated with the Dumbo phenotype in the pectoral fins of Dumbo and non-Dumbo phenotypes.**

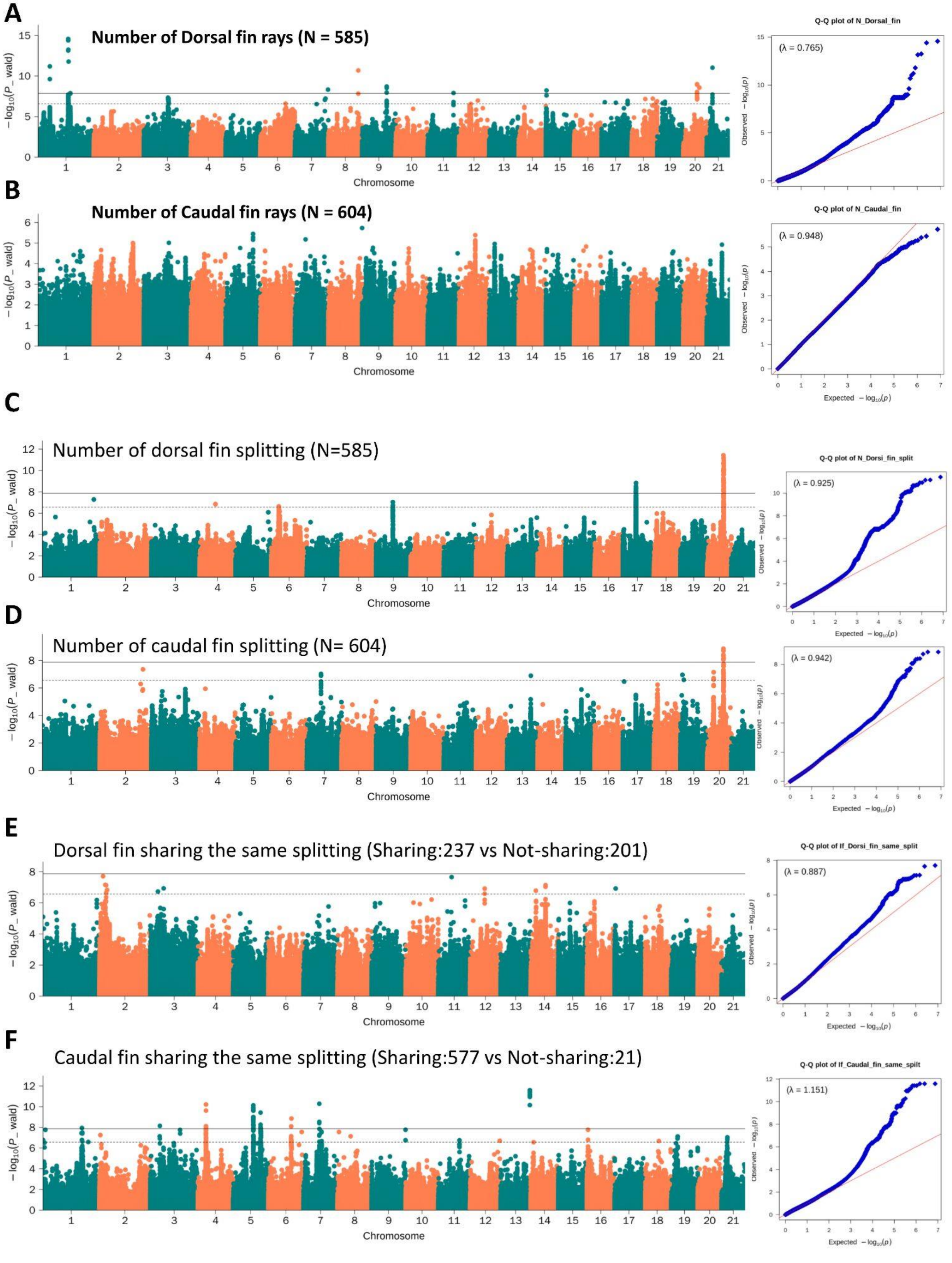

A
Number of Dorsal fin rays (N = 585)
Q-Q plot of N_Dorsal_fin
(λ = 0.765)
B
Number of Caudal fin rays (N = 604)
Q-Q plot of N_Caudal_fin
(λ = 0.948)
C
Number of dorsal fin splitting (N=585)
Q-Q plot of N_Dorsi_fin_split
(λ = 0.925)
D
Number of caudal fin splitting (N= 604)
(λ = 0.942)
E
Dorsal fin sharing the same splitting (Sharing:237 vs Not-sharing:201)
Q-Q plot of If_Dorsi_fin_same_split
(λ = 0.887)
F
Caudal fin sharing the same splitting (Sharing:577 vs Not-sharing:21)
Q-Q plot of If_Caudal_fin_same_spilt
(λ = 1.151)
−log10(P_wald)
Chromosome
Observed −log10(p)
Expected −log10(p)

**Fig. S32. Manhattan and QQ plots for GWAS for traits related to fin rays in the Siamese fighting fish.**

(A) Manhattan and QQ plot for number of dorsal fin rays GWAS (N=585). (B) Manhattan and QQ plot for number of caudal fin rays GWAS (N=604). (C) Manhattan and QQ plot for GWAS of number of dorsal fin ray splitting (N=585). (D) Manhattan and QQ plot for GWAS of number of caudal fin ray splitting (N=604). (E) Manhattan and QQ plot for GWAS of unanimous dorsal fin splitting (N=237 vs. 201). (F) Manhattan and QQ plot for GWAS of unanimous caudal fin splitting (N=577 vs. 21). The solid and dashed lines in the Manhattan plots represent genome-wide (0.05/Nsnp) and chromosomal level (1/Nsnp) significance thresholds, respectively. Red line in the QQ plots shows expected *P*-values and Blue dots show observed *P*-values. Traits are described in detail in Supplementary Text 4 and associated loci are discussed in Supplementary Text 8.

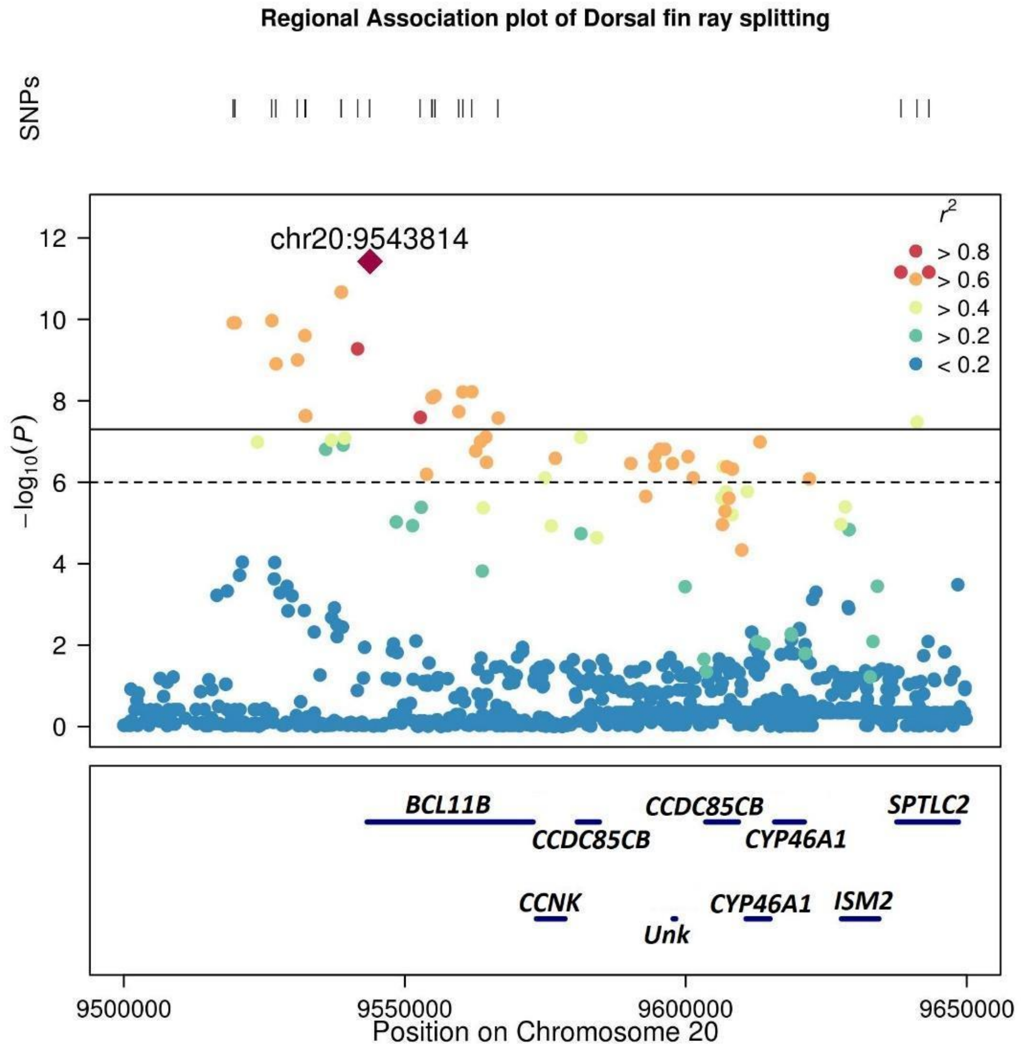


**Fig. S33. Regional plot of the locus with the strongest association in a GWAS for maximum number of dorsal fin ray splitting in the Siamese fighting fish.**

The solid and dashed lines represent genome-wide (0.05/Nsnp) and chromosomal level (1/Nsnp) significance thresholds, respectively. Colors represent linkage disequilibrium with the top variant, indicated by a diamond symbol. Bottom box shows the location of annotated genes in this region. Corresponding genome-wide Manhattan plot is shown in Fig. S32C.

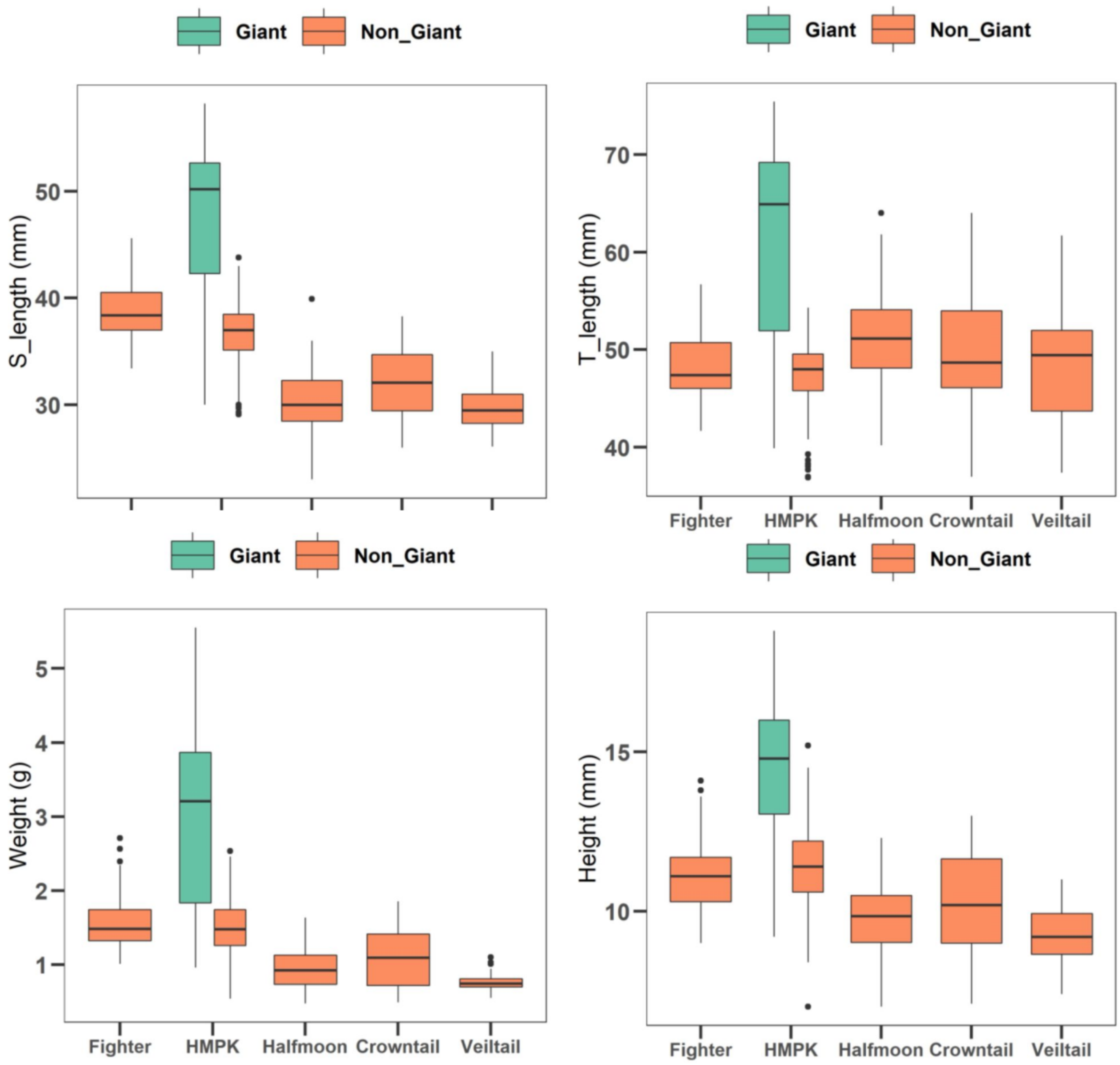


**Fig. S34. Body size measurements in the Giant HMPK compared to other breeds.**

The Giant HMPK Betta fish is larger than other breeds in all four body size measurements taken: Weight, Height, T_length (total length) and S_length (standard length).

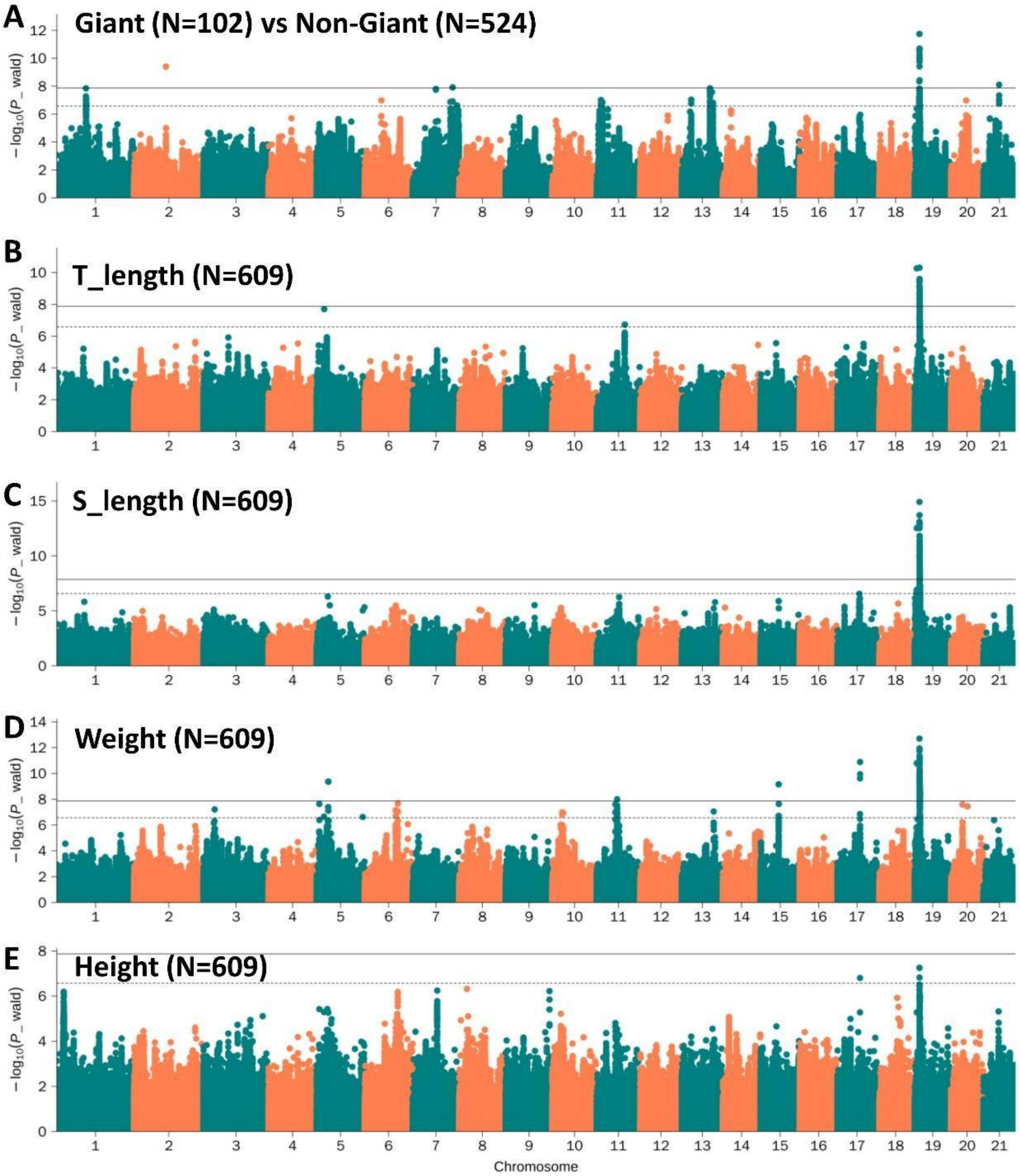


**Fig. S35. Manhattan plots for the body size GWAS.**

Manhattan plots for body size GWAS using different measurements of body size in the Siamese fighting fish. (A) Manhattan plot for Giant (N=102) vs. non-Giant phenotype (N=524) GWAS. (B) Manhattan plot for total length (T_length, N=609) GWAS. (C) Manhattan plot for standard length (S_length, N=609) GWAS. (D) Manhattan plot for Weight (N=609) GWAS. (E) Manhattan plot for Height (N=609) GWAS. The solid and dashed lines represent genome-wide (0.05/Nsnp) and chromosomal level (1/Nsnp) threshold, respectively. Corresponding QQ plots are in Fig. S37. A regional plot for the significantly associated variant on chromosome 19 is in Fig. S39.

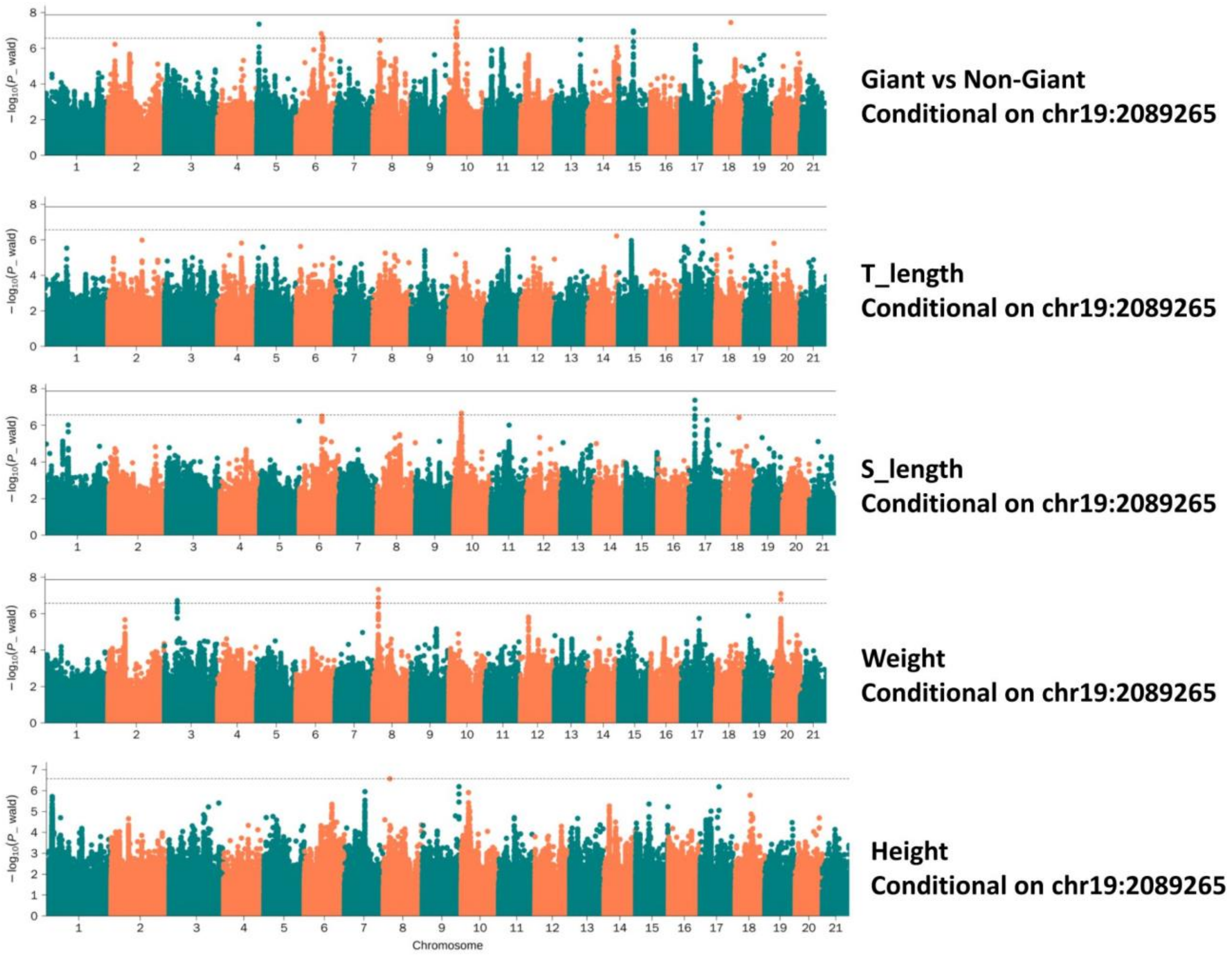


**Fig. S36. GWAS for body size conditioning on top SNP associated with body size.**

Manhattan plots of GWAS for body size measurements conditioning on the top SNP associated with body size (chr19:2089265). The solid and dashed lines represent genome-wide (0.05/Nsnp) and chromosomal level (1/Nsnp) significance thresholds, respectively.

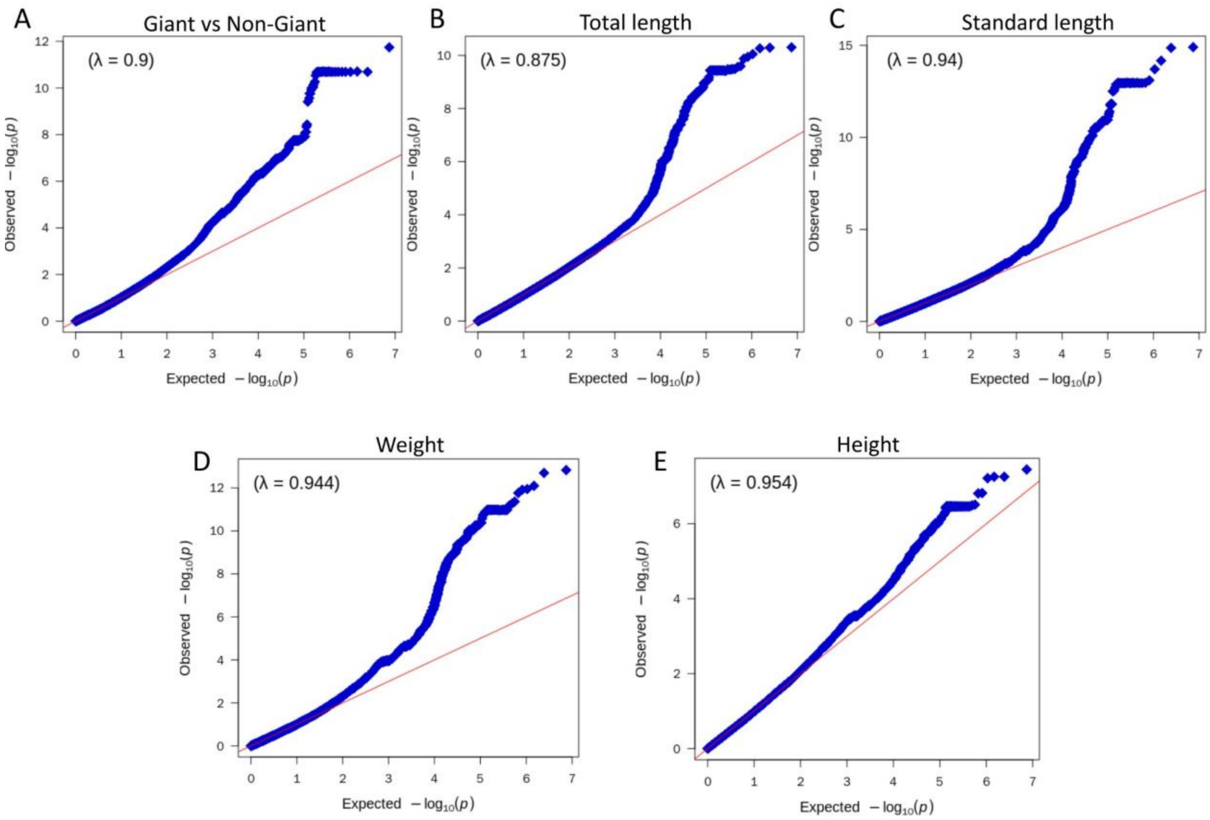


**Fig. S37. QQ-plot of the body size GWAS results.**

QQ plots for body size GWAS using different measurements of body size in the Siamese fighting fish. (A) QQ plot for Giant (N=102) vs. non-Giant phenotype (N=524) GWAS. (B) QQ plot for total length (N=609) GWAS. (C) QQ plot for standard length (N=609) GWAS. (D) QQ plot for Weight (N=609) GWAS. (E) QQ plot for Height (N=609) GWAS. Red line shows expected P-values and blue dots show observed P-values. Corresponding Manhattan plots are in Fig. S35.

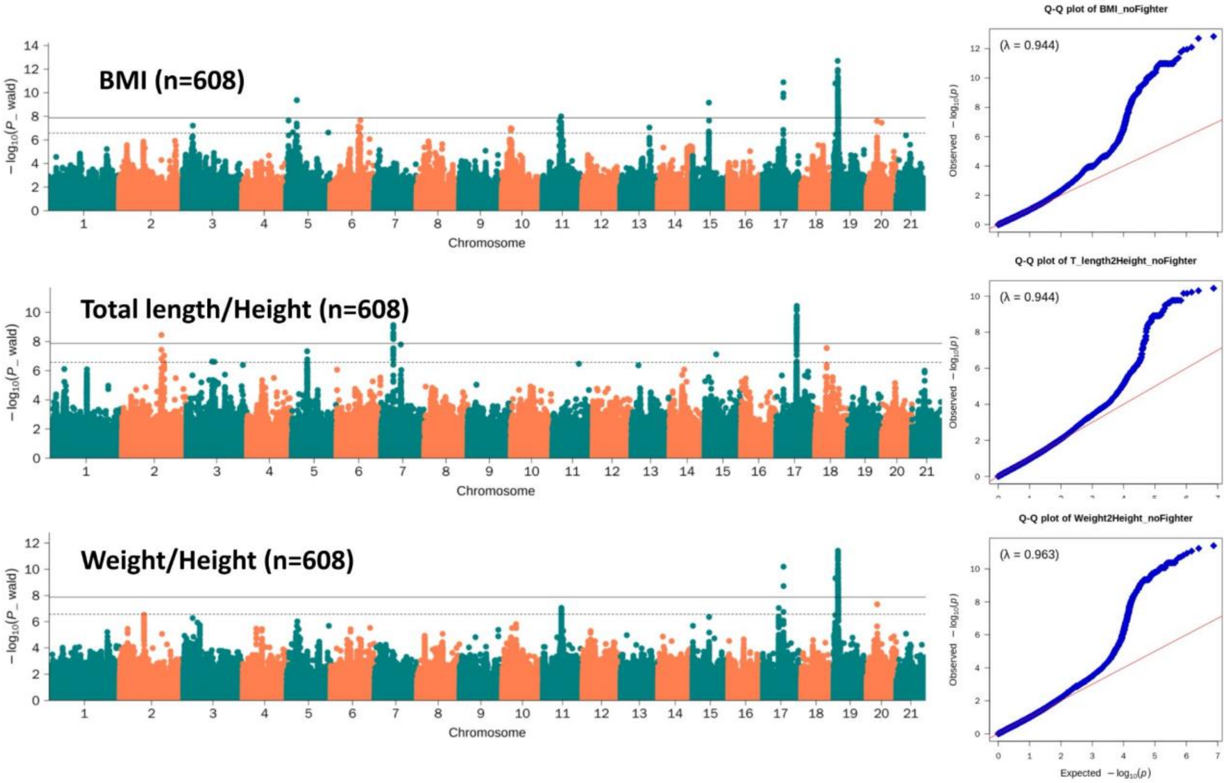


**Fig. S38. GWAS of other body size traits: BMI, Total length/Height and Weight/Height.**

BMI and Weight/Height are associated with the same locus on chromosome 19 as other body size traits. Total length/Height is associated with a locus on chromosome 17 also found to be associated with long-fin breeds. See discussion on Supplementary Text 9. The solid and dashed lines in the Manhattan plots represent genome-wide (0.05/Nsnp) and chromosomal level (1/Nsnp) significance thresholds, respectively. Red line in the QQ plots shows expected P-values and blue dots show observed P-values.

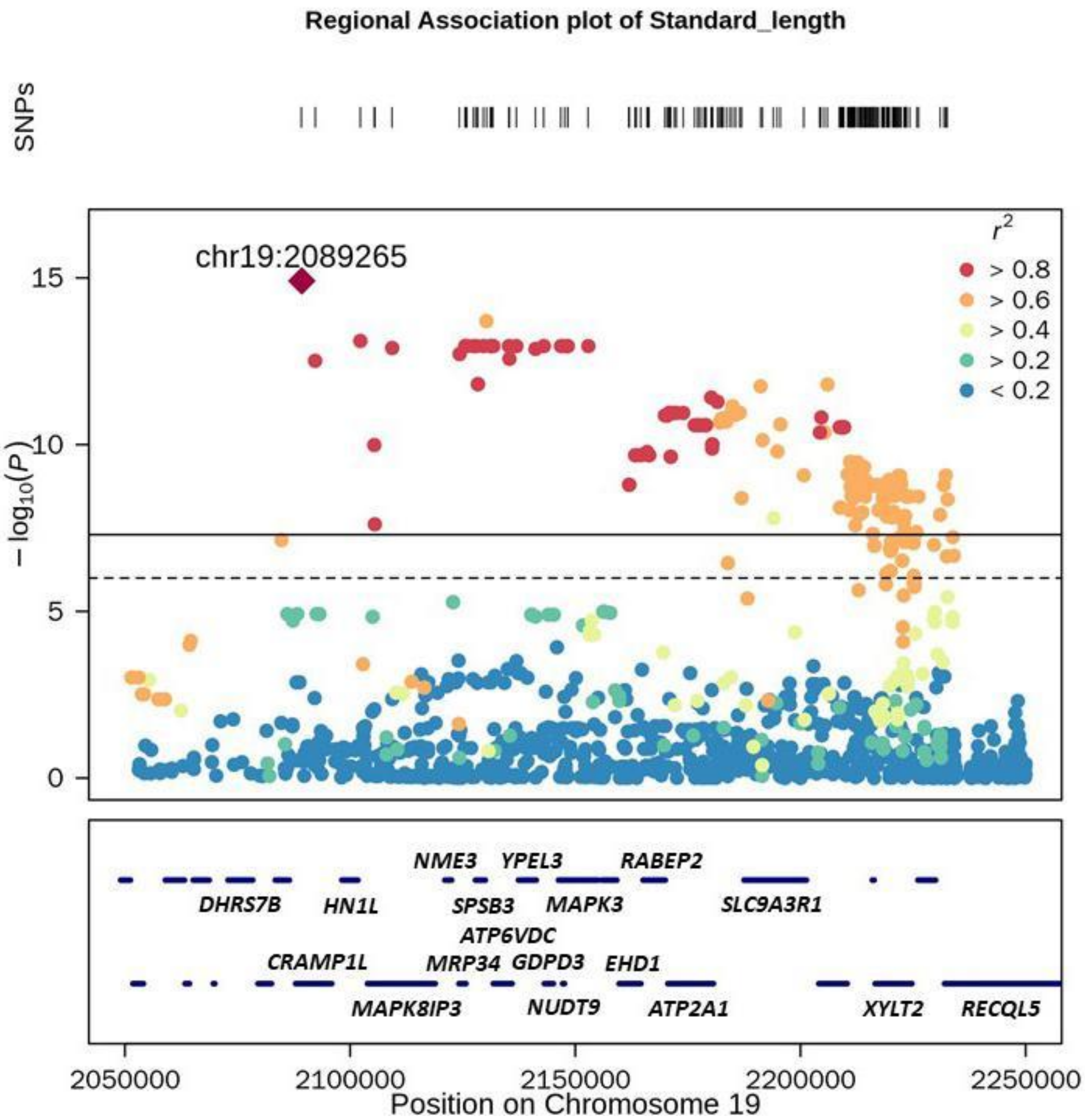


**Fig. S39. Regional plot of the locus with the strongest association in a GWAS for standard length in the Siamese fighting fish.**

This locus is also significantly associated with other body size measurements. Genome-wide Manhattan plots for all body size related phenotypes are shown in Figs. S35 and S38. The solid and dashed lines in the top box represent genome-wide (0.05/Nsnp) and chromosomal level (1/Nsnp) significance thresholds, respectively. Colors represent linkage disequilibrium with the top variant, indicated by a diamond symbol. Bottom box shows the location of annotated genes in this region.

| Gene_name | FRKM (HMPK_Brain) | FRKM (Giant_Brain) | log2FoldChange | padj | FRKM (HMPK_Muscle) | FRKM (Giant_Muscle) | log2FoldChange | padj |
|---|---|---|---|---|---|---|---|---|
| MYO15A | 8.2588 | 9.4303 | -0.1854 | 1.0000 | 0.3727 | 0.2472 | 0.3373 | 1.0000 |
| DRG2 | 101.2585 | 112.7125 | -0.1559 | 0.8709 | 485.1772 | 108.7030 | 2.1588 | 0.0002 |
| Gfer | 60.0241 | 58.5446 | 0.0346 | 0.9675 | 46.6983 | 10.8919 | 2.1097 | 0.0112 |
| SYNGR3 | 232.3504 | 155.3216 | 0.5819 | 0.1919 | 2.6663 | 1.8581 | 0.5511 | 0.8630 |
| znf598 | 16.2735 | 50.9340 | -1.6354 | 0.0118 | 27.0038 | 29.6491 | -0.1312 | 0.9700 |
| tmem11 | 57.5285 | 87.0772 | -0.6014 | 0.1623 | 63.2667 | 76.2405 | -0.2713 | 0.7201 |
| dhrs7b | 104.5097 | 198.7563 | -0.9293 | 0.0198 | 79.1274 | 42.0584 | 0.9030 | 0.3156 |
| CRAMP1L | 620.0873 | 578.7311 | 0.0989 | 0.6156 | 198.7876 | 182.2206 | 0.1242 | 0.8271 |
| HN1L | 358.3685 | 366.4853 | -0.0326 | 0.8849 | 2155.9693 | 1008.9539 | 1.0949 | 0.0026 |
| MAPK8IP3 | 7234.5069 | 6967.9599 | 0.0541 | 0.8741 | 190.4357 | 68.2380 | 1.4754 | 0.0001 |
| Nme3 | 132.9066 | 156.2153 | -0.2327 | 0.3766 | 20.2205 | 14.0957 | 0.5062 | 0.5524 |
| MRPS34 | 275.8321 | 336.6893 | -0.2882 | 0.0636 | 388.0349 | 538.5605 | -0.4742 | 0.3261 |
| Spsb3 | 331.7391 | 282.2561 | 0.2327 | 0.3372 | 87.0821 | 111.8205 | -0.3596 | 0.4979 |
| ATP6V0C | 7671.8765 | 10195.9468 | -0.4103 | 0.0000 | 512.5196 | 342.6398 | 0.5812 | 0.0335 |
| ypel3 | 1354.9371 | 1418.4085 | -0.0663 | 0.7693 | 522.3964 | 576.5228 | -0.1397 | 0.7743 |
| GDPD3 | 5.9768 | 12.2065 | -1.0298 | 1.0000 | 22.6721 | 20.6519 | 0.1399 | 0.9253 |
| MAPK3 | 3045.7249 | 2809.1412 | 0.1167 | 0.4902 | 579.7450 | 551.5371 | 0.0726 | 0.8683 |
| NUDT9 | 226.7370 | 211.9072 | 0.0976 | 0.6383 | 114.2462 | 86.8029 | 0.4071 | 0.4548 |
| EHD1 | 161.1696 | 173.2895 | -0.1049 | 0.7521 | 66.2119 | 72.3107 | -0.1279 | 0.8873 |
| RABEP2 | 156.9859 | 158.1046 | -0.0104 | 0.9765 | 337.2740 | 206.8134 | 0.7020 | 0.1754 |
| atp2a1 | 117.4310 | 119.6976 | -0.0278 | 0.9899 | 7780.6456 | 17926.5327 | -1.2041 | 0.0858 |
| Slc9a3r1 | 781.0012 | 864.6145 | -0.1468 | 0.5196 | 483.6753 | 492.3208 | -0.0248 | 0.9694 |
| RAB37 | 659.1946 | 877.1452 | -0.4122 | 0.1013 | 88.1456 | 71.2498 | 0.3132 | 0.7857 |
| XYLT2 | 283.3911 | 228.8644 | 0.3083 | 0.1353 | 264.3860 | 126.8385 | 1.0600 | 0.0872 |
| XYLT2_2 | 39.8910 | 55.8189 | -0.4888 | 0.2877 | 16.0931 | 19.1366 | -0.2381 | 0.8487 |
| Recql5 | 389.1490 | 226.5617 | 0.7801 | 0.0053 | 137.2944 | 85.0148 | 0.6949 | 0.1422 |

**Fig. S40. Expression in muscle and brain tissues of genes in the 119 kb region on chromosome 19 associated with body size.**

Candidate loci associated with body size, including Giant vs. non-Giant breeds are further discussed in Supplementary Text 9.

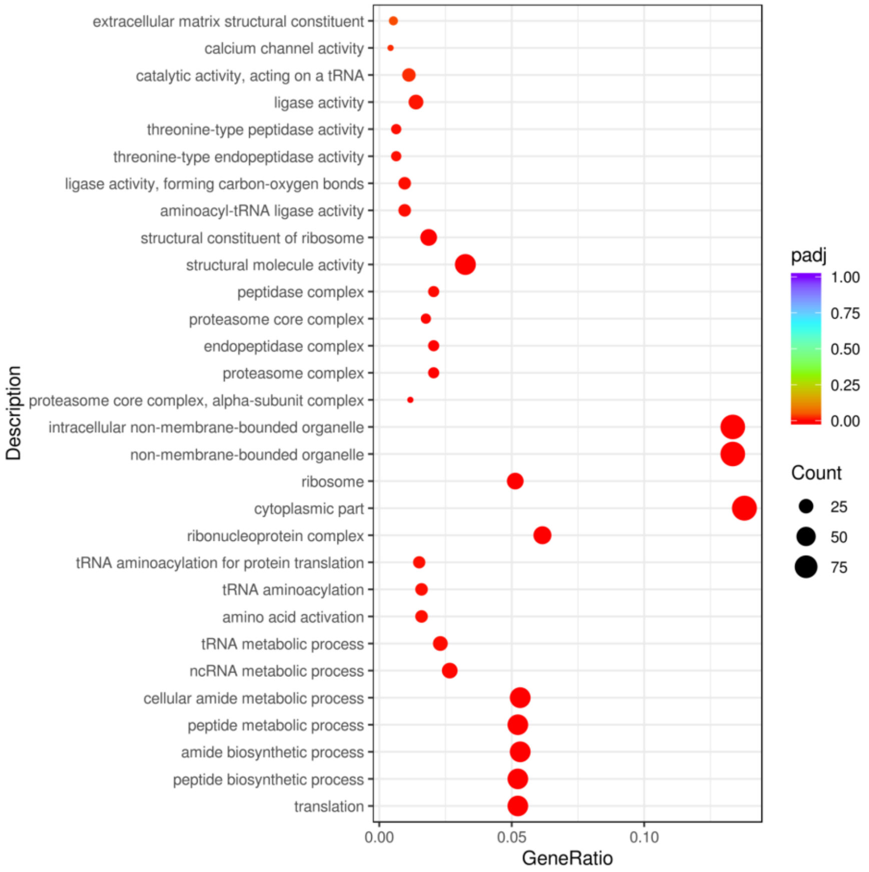


**Fig. S41. Gene Ontology (GO) terms enriched in genes differentially expressed in muscle between the Giant-HMPK and HMPK.**

Size of circles represent the number of genes, colors represent adjusted p-value from the differential expression analysis. GO categories are listed on the y-axis and Gene Ratio on the x-axis is the percentage of differentially expressed genes in the GO category.

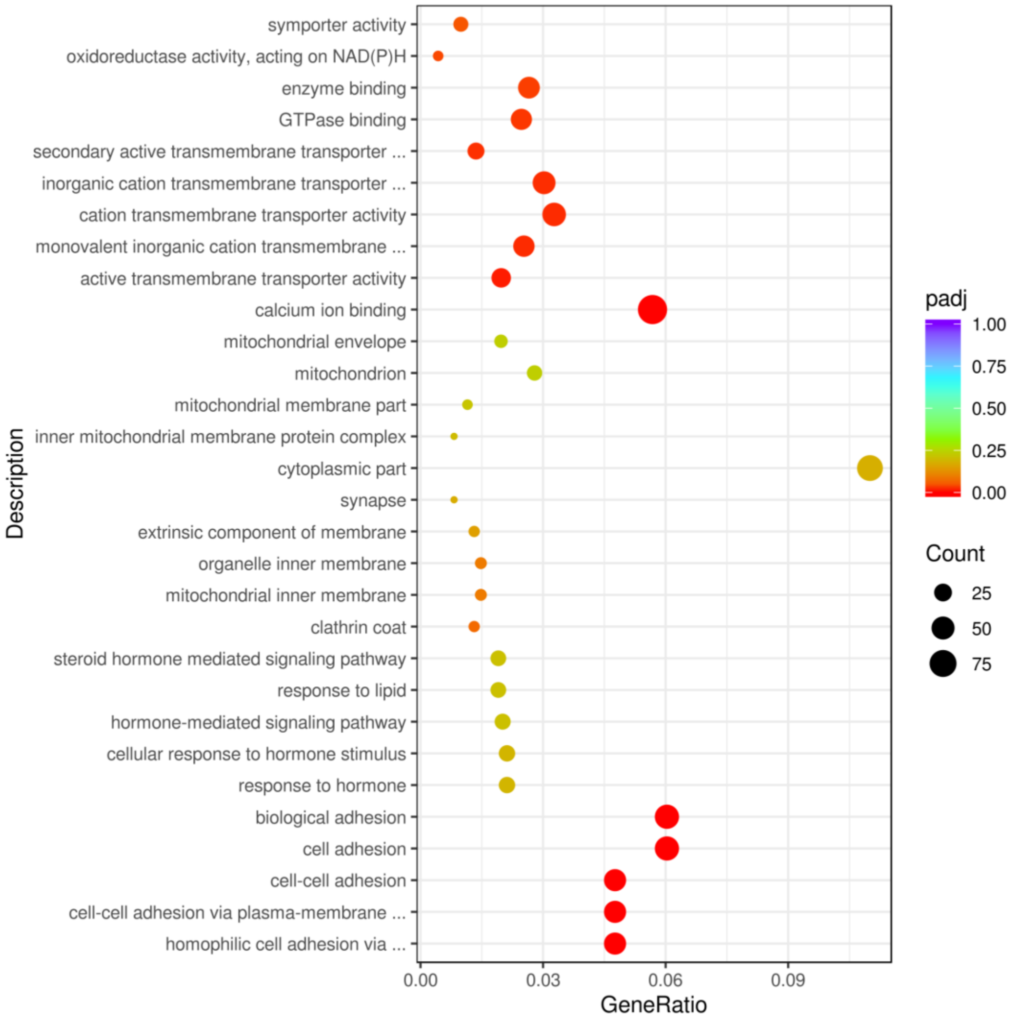


**Fig. S42. Gene Ontology (GO) terms enriched in genes differentially expressed in the brain between the Giant-HMPK and HMPK.**

Size of circles represent the number of genes, colors represent adjusted p-value from the differential expression analysis. G) categories are listed on the y-axis and Gene Ratio on the x-axis is the percentage of differentially expressed genes in the GO category.

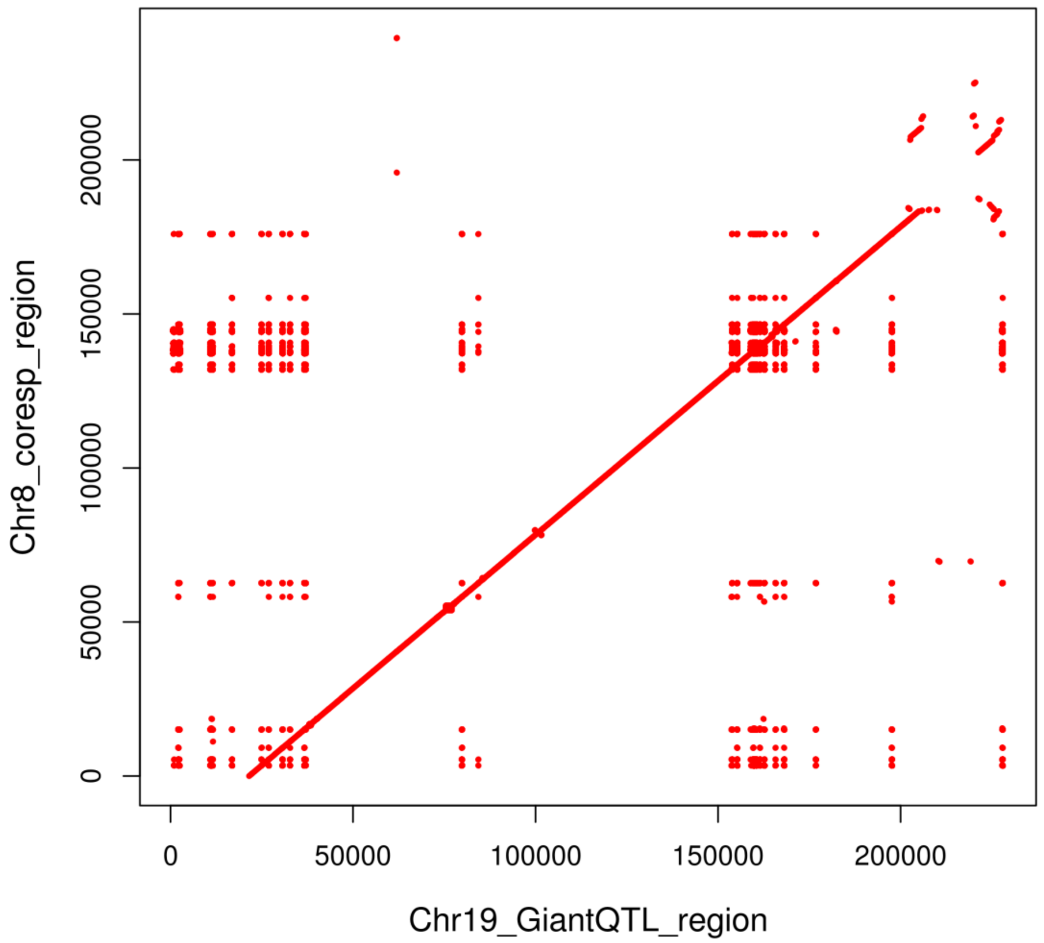


**Fig. S43. Synteny analysis of the locus associated with the Giant phenotype.**

The plot is a comparison between our genome assembly (x axis) of the Siamese fighting fish genome and a previously published assembly (fBetSpl5.3, GenBank accession: GCA_003650155.3).

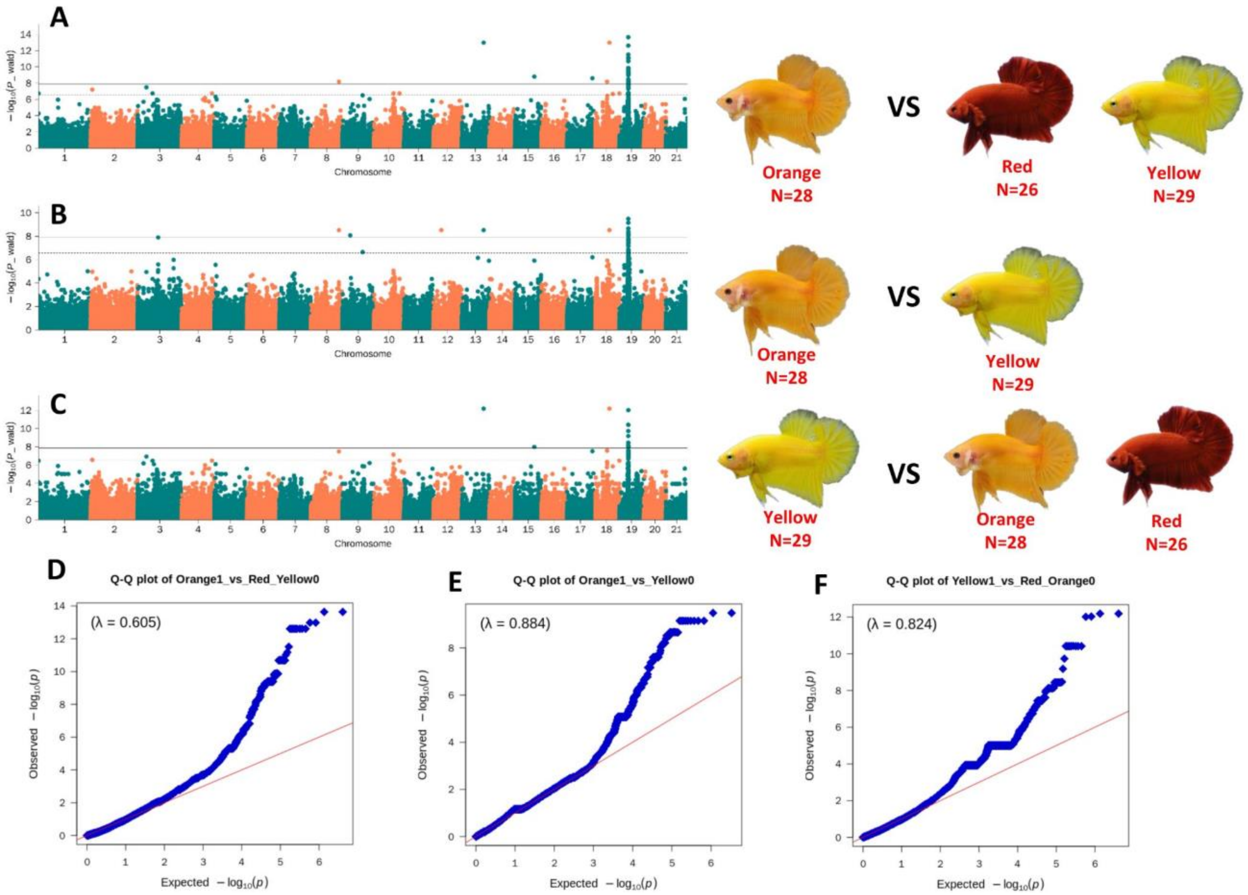


**Fig. S44. GWAS for red, yellow and orange color breeds in the Siamese fighting fish.**

(A, D) Manhattan and QQ plot for a case-control GWAS of Orange as cases compared to Red and Yellow as controls. (B, E) Manhattan and QQ plot for a case-control GWAS of Orange as cases compared to Yellow as controls. (C, F) Manhattan plot for GWAS using Yellow as cases compared to Orange and Red as controls. The solid and dashed lines in the Manhattan plots represent genome-wide (0.05/Nsnp) and chromosomal level (1/Nsnp) significance thresholds, respectively. Red line in the QQ plots shows expected P-values and blue dots show observed P-values.

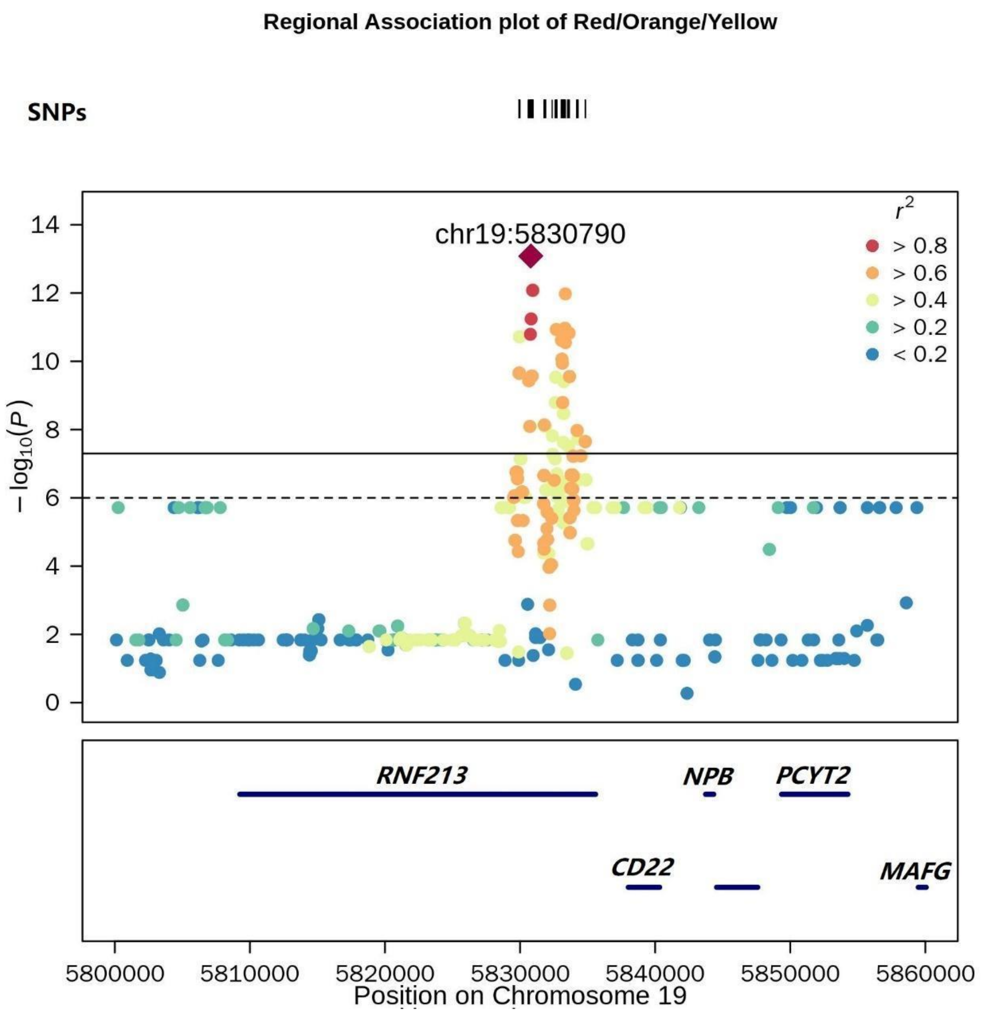


**Fig. S45. Regional plot of the locus with the strongest association in a GWAS between Orange, Yellow and Red breeds in the Siamese fighting fish.**

The solid and dashed lines represent genome-wide (0.05/Nsnp) and chromosomal level (1/Nsnp) significance thresholds, respectively. Colors represent linkage disequilibrium with the top variant, indicated by a diamond symbol. Bottom box shows the location of annotated genes in this region.

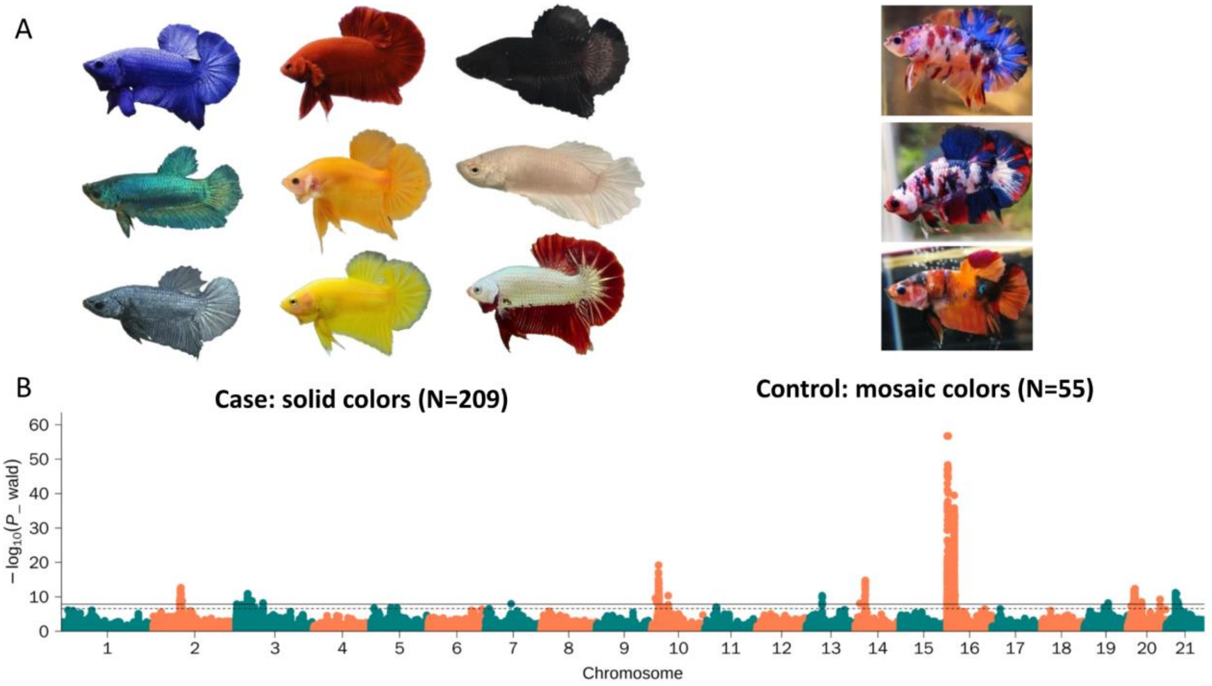


**Fig. S46. GWAS for solid color and mosaic breeds in the Siamese fighting fish.**

(A)Illustration of solid color breeds (n=209) and mosaic color breeds (n=55). (B) Manhattan plot. The solid and dashed lines represent genome-wide (0.05/Nsnp) and chromosomal level (1/Nsnp) significance thresholds in the Manhattan plot, respectively.

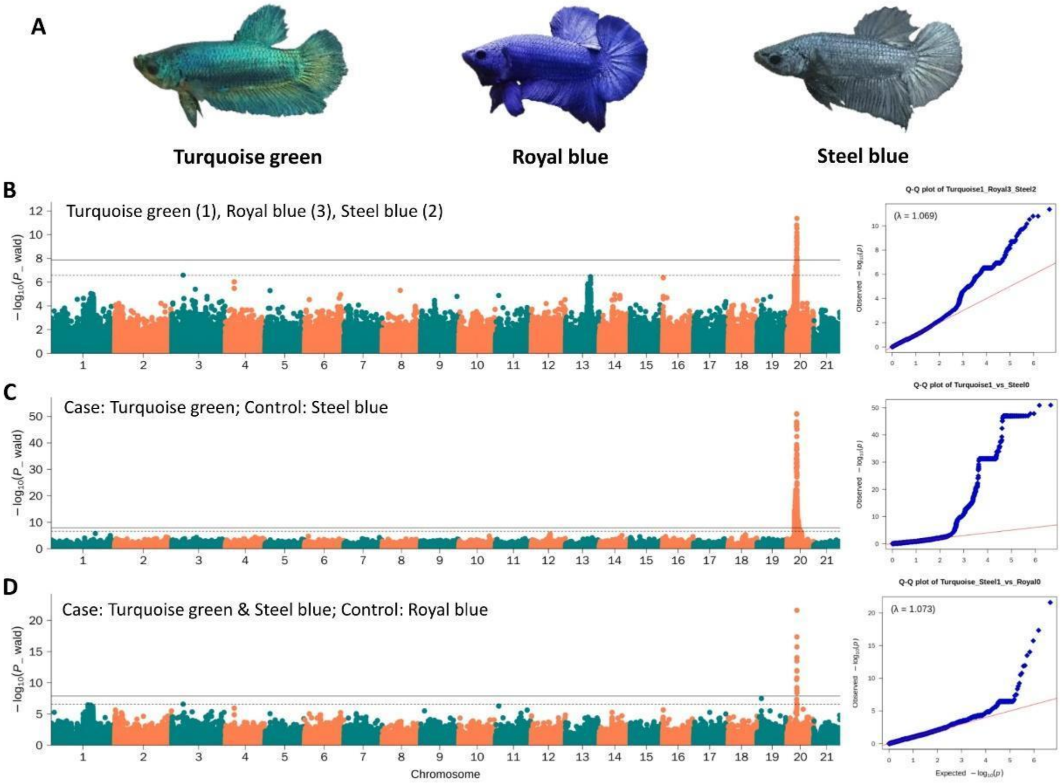


**Fig. S47. GWAS for Turquoise green, Royal blue and Steel blue color breeds in the Siamese fighting fish.**

(A) Illustration of Turquoise-green (N=26), Royal-blue (N=28) and Steel-blue (N=35) breeds used in this study. (B) Manhattan and QQ plot for treating the three colors as discrete phenotypes in GWAS. (C) Manhattan and QQ plot for using Turquoise-green as cases and Steel-blue as controls in GWAS. (D) Manhattan and QQ plot for using Turquoise-green and Steel-blue as cases and Royal-blue as controls in GWAS. The solid and dashed lines in the Manhattan plots represent genome-wide (0.05/Nsnp) and chromosomal level (1/Nsnp) significance thresholds, respectively. Red line in the QQ plots shows expected P-values and blue dots show observed P-values.

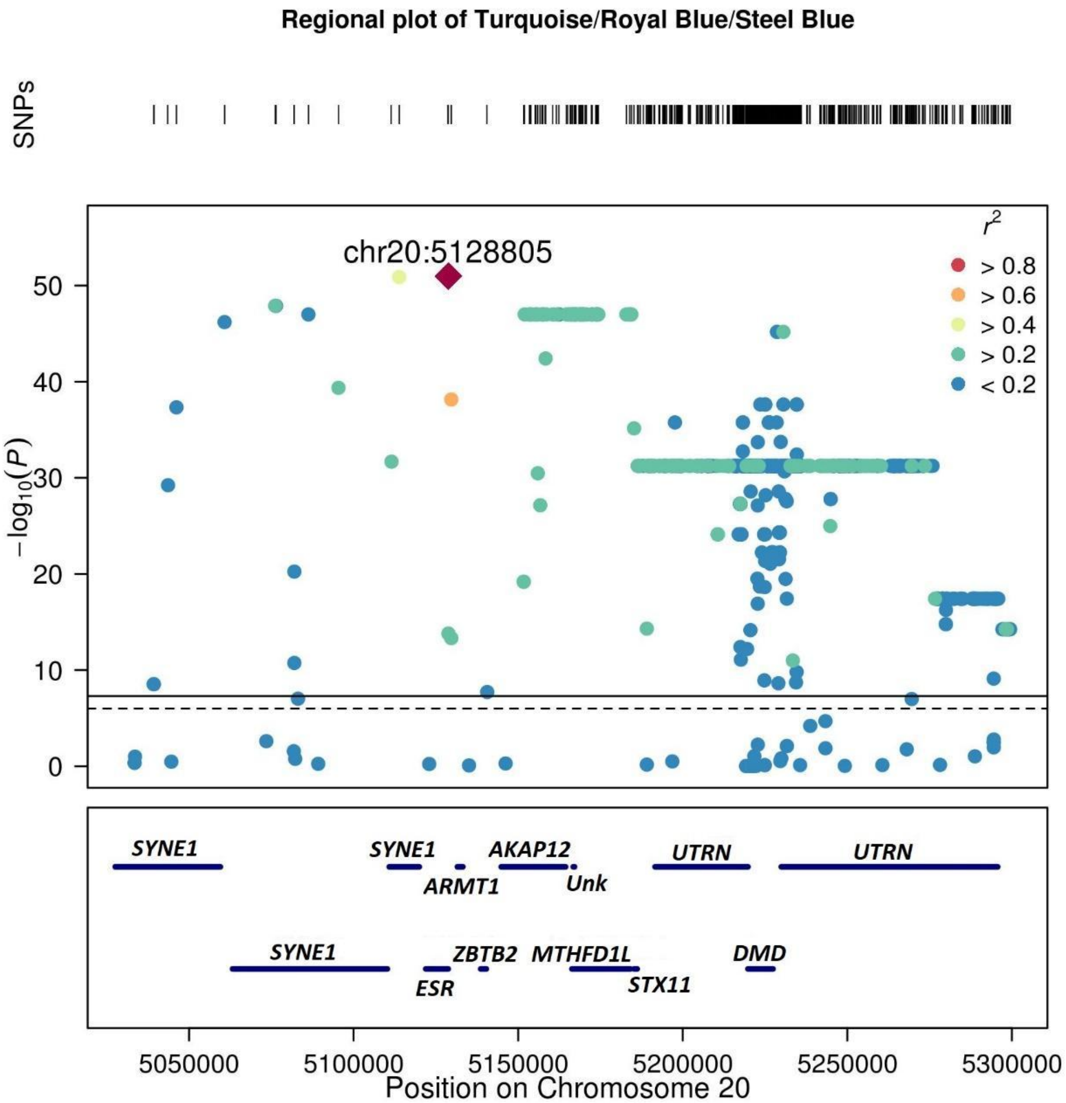


**Fig. S48. Regional plot of the locus with the strongest association in a GWAS between Turquoise green, Royal blue and Steel blue breeds in the Siamese fighting fish.**

The solid and dashed lines represent genome-wide (0.05/Nsnp) and chromosomal level (1/Nsnp) significance thresholds, respectively. Colors represent linkage disequilibrium with the top variant, indicated by a diamond symbol. Bottom box shows the location of annotated genes in this region.

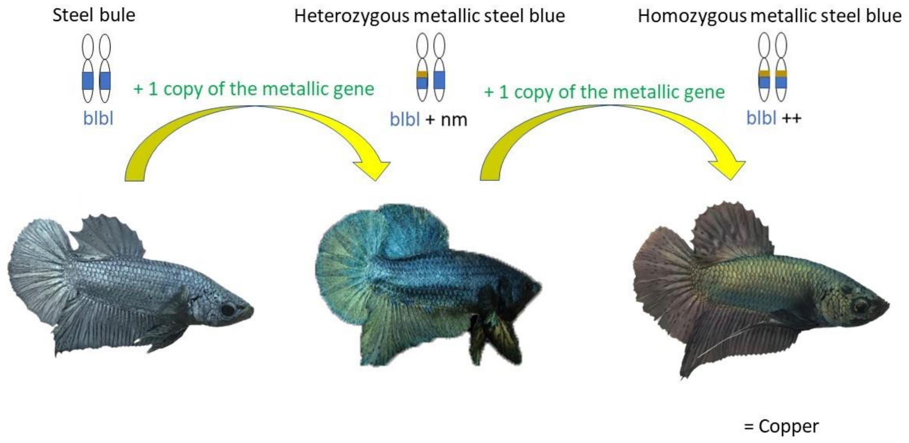


**Fig. S49. A schematic of the breeding of Copper from Steel blue.**

The Copper phenotype is derived from two copies of a presumed “metallic” gene on a Steel-blue genetic background.

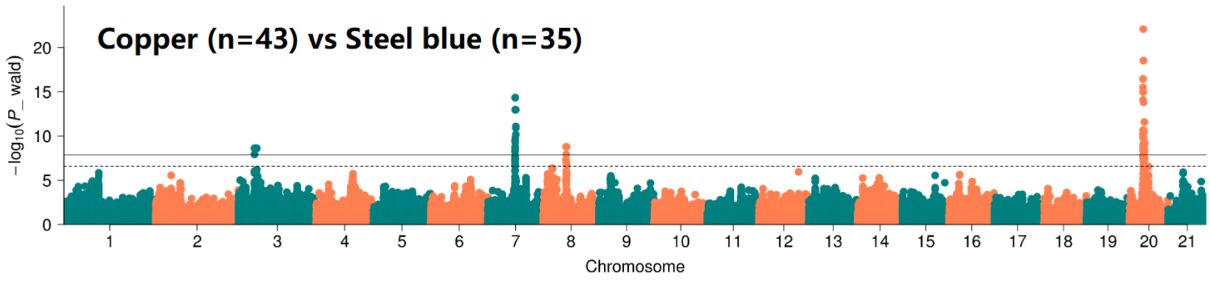


**Fig. S50. GWAS of Copper versus Steel blue breeds.**

Two loci are associated with Copper vs. Steel blue phenotypes. The locus on chromosome 20 is also associated with the Turquoise-green, Royal-blue and Steel-blue variants, as discussed in Supplementary Text 10. The solid and dashed lines represent genome-wide (0.05/Nsnp) and chromosomal level (1/Nsnp) significance thresholds, respectively.

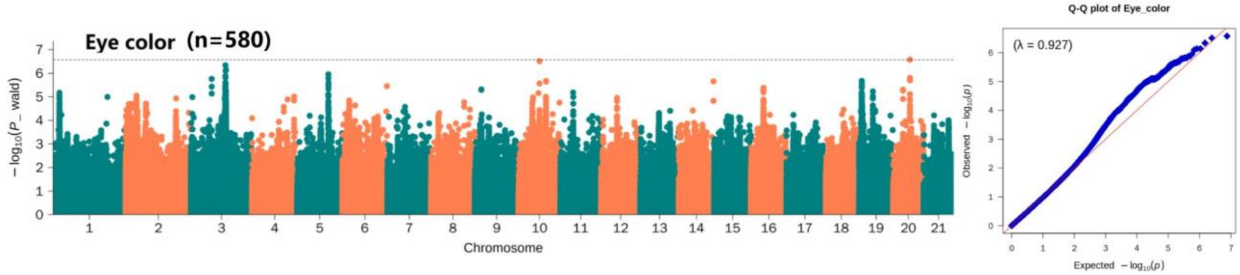


**Fig. S51. GWAS results of eye color in the Siamese fighting fish.**

No variant significantly associated with eye color in *Betta splendens* was found. The dashed line in the Manhattan plot represents the chromosomal level (1/Nsnp) significance threshold. Red line in the QQ plots shows expected P-values and blue dots show observed P-values.

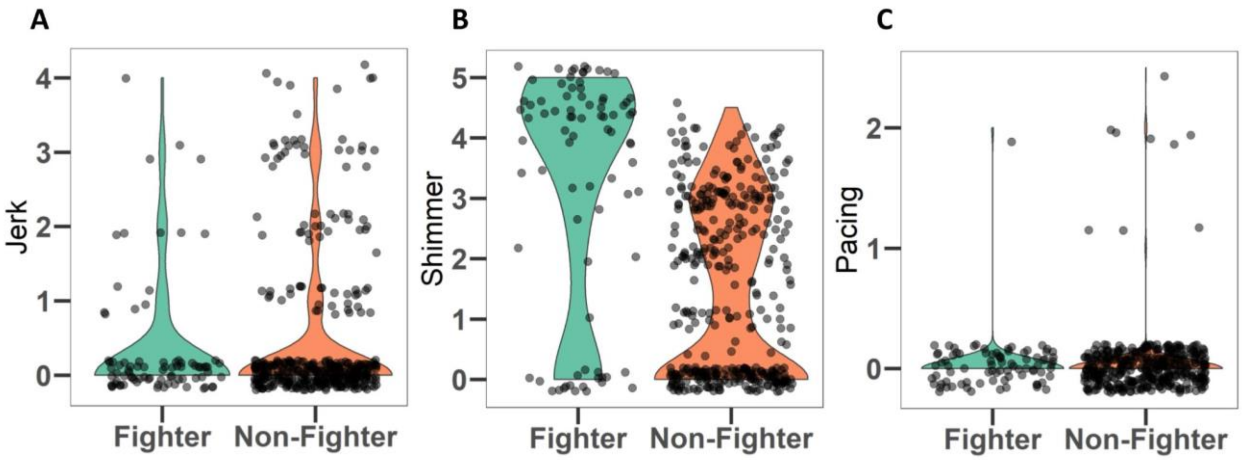


**Fig. S52: Boxplot for aggression related phenotypes composing the aggression index.**

Other indices of aggression are shown in Fig. 4C-J. The description of each phenotype and their contribution to the aggression index is given in Supplementary text 4.

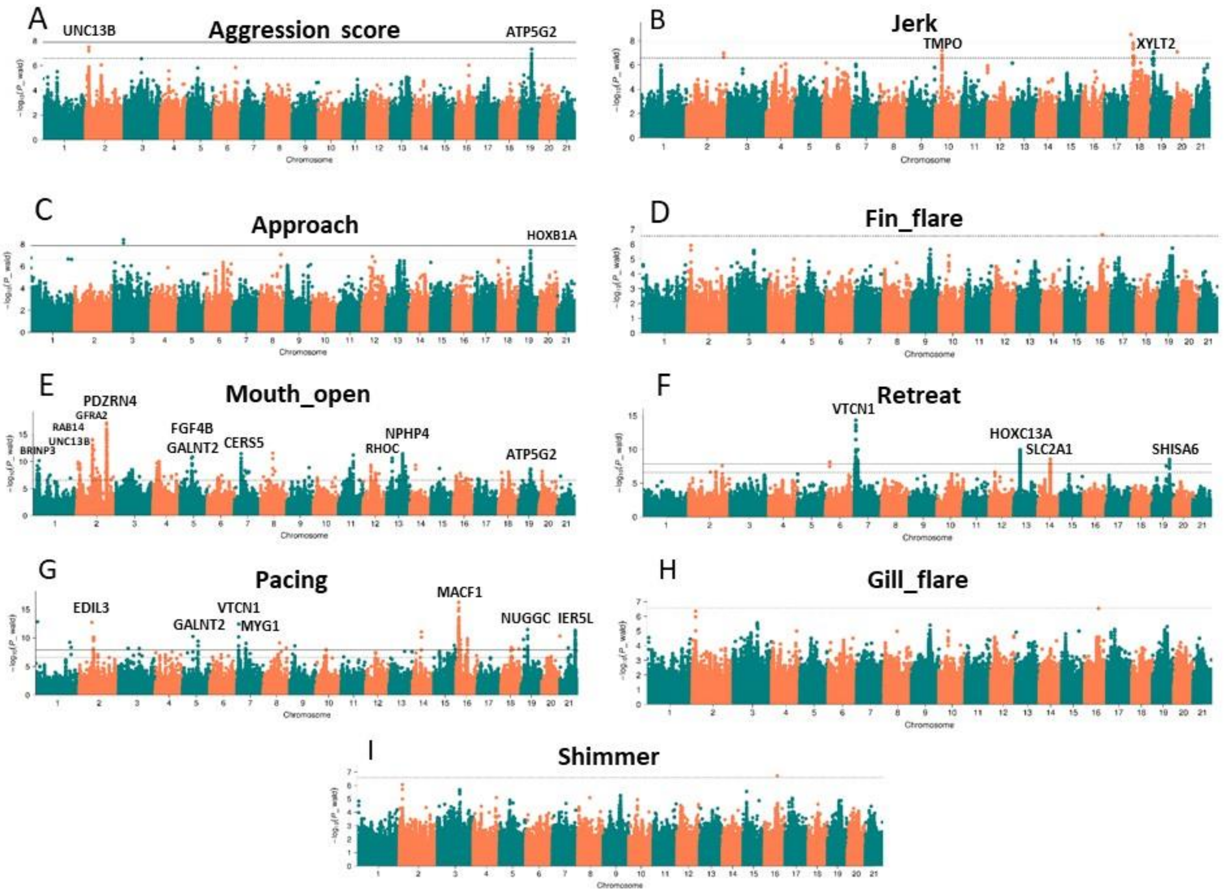


**Fig. S53. Manhattan plot of GWAS for aggression score and the phenotypes composing it.**

A definition of each aggressive phenotype and of the aggression score is given in Supplementary Text 4. The solid and dashed lines in the Manhattan plots represent genome-wide (0.05/Nsnp) and chromosomal level (1/Nsnp) significance thresholds, respectively. Genes significantly associated with each phenotype are annotated.

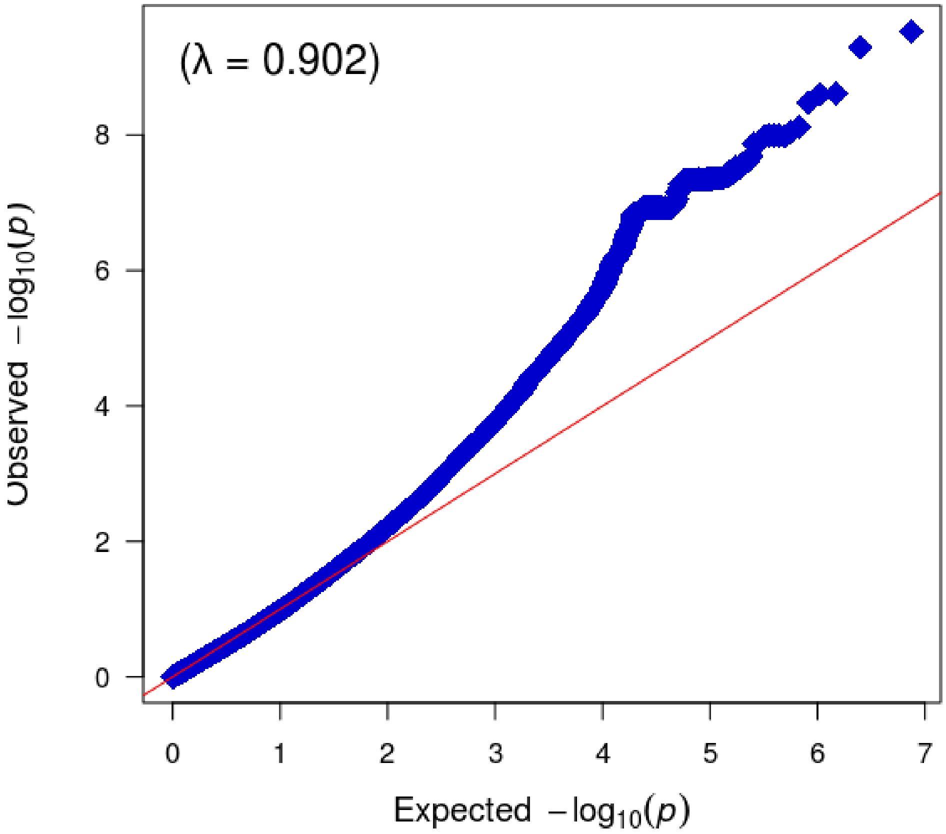


**Fig. S54. Quantile-quantile (QQ) plot for GWAS of times of breathing air in one minute in the Siamese fighting fish.**

Red line shows expected P-values and blue dots show observed P-values. The Manhattan plot for the corresponding GWAS is in Fig. 4L in the Main Text.

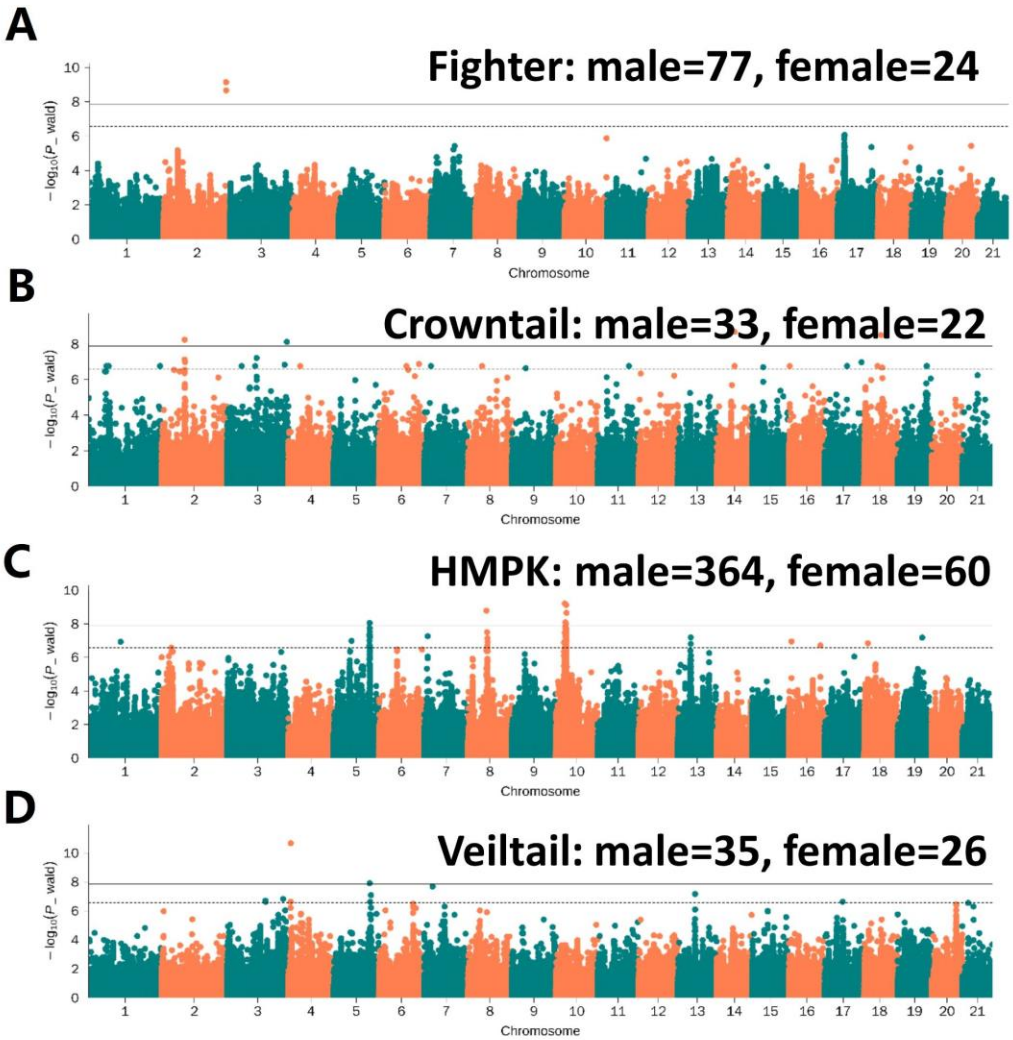


**Fig. S55. GWAS for sex conditioning on the top SNP associated with sex in the four domesticated breeds of *B. splendens* analysed.**

Manhattan plots of GWAS for sex conditioning on the top SNP associated with sex (chr2:2839325). The solid and dashed lines represent genome-wide (0.05/Nsnp) and chromosomal level (1/Nsnp) significance thresholds, respectively.

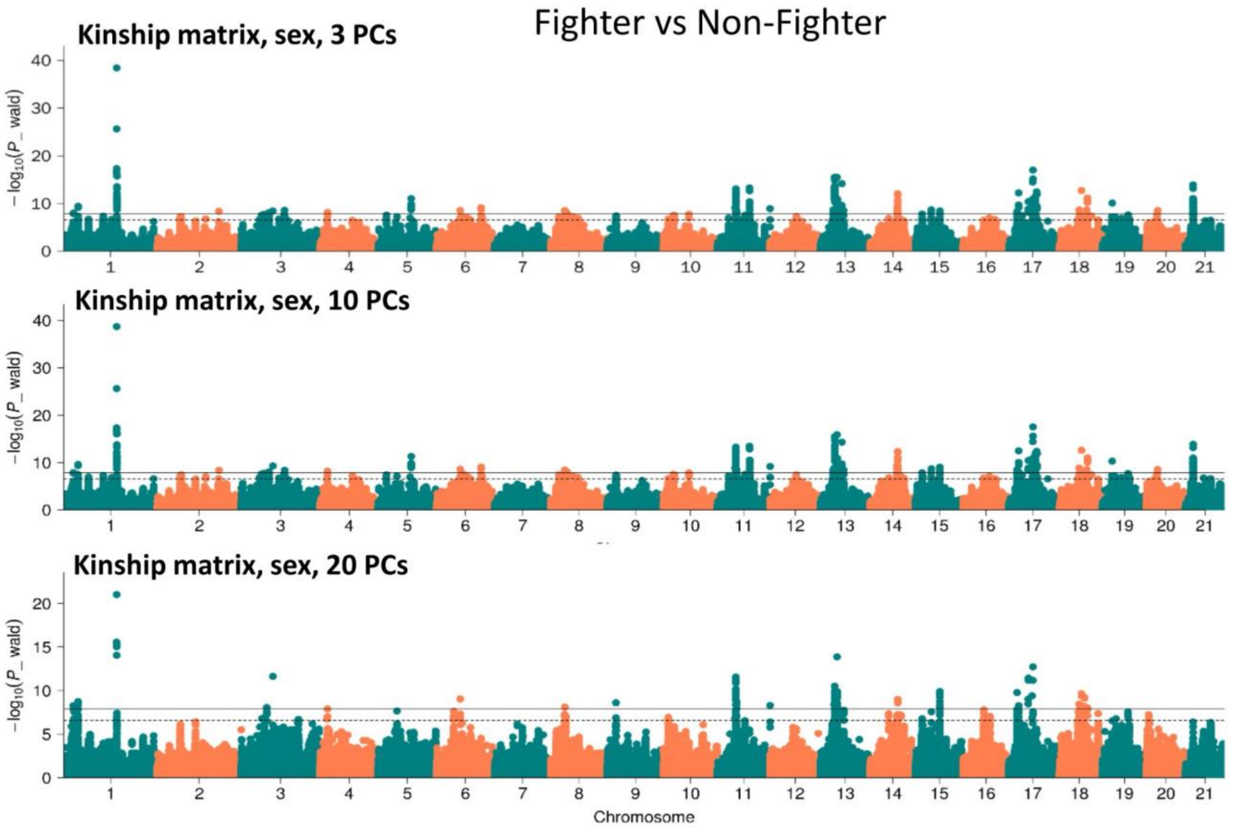


**Fig. S56. GWAS for Fighter vs Non-Fighter results using different numbers of PCs for correction.**

The solid and dashed lines represent genome-wide (0.05/Nsnp) and chromosomal level (1/Nsnp) significance thresholds, respectively.

### Supplementary Tables

#### Table S1. Statistics of three Siamese fighting fish genome assemblies.

| | ASM365015v1[1] (male: XY) | fBetSpl5.3[2] (male: XY) | BettaSRF1.0[3] (female: XX) |
|---|---|---|---|
| **Assembly Statistics** | | | |
| Sequencing platforms (Depth) | Illumina (114×)<br>Hi-C (75×) | PacBio (48×)<br>Illumina (83×)<br>10X Genomics (183×)<br>BioNano | PacBio (109×)<br>Illumina (109×)<br>10X Genomics (235×)<br>BioNano (319×)<br>Hi-C (107×) |
| Total sequence length (Mb) | 456.232 | 441.388 | 451.294 |
| Total ungapped length (Mb) | 412.702 | 441.359 | 448.889 |
| Total Gap length (Mb) | 43.53 (9.54%) | 0.029 (0.0065%) | 2.405 (0.53%) |
| Number of scaffolds | 12,333 | 70 | 318 |
| Scaffold N50 (Mb) | 19.754 | 20.129 | 19.630 |
| Scaffold L50 | 10 | 9 | 9 |
| Number of contigs | 35,779 | 398 | 526 |
| Contig N50 (bp) | 38,053 | 2,497,747 | 4,077,672 |
| Contig L50 | 3,417 | 48 | - |
| GC content | 40.04% | 45.10% | 45.08% |
| Unplaced | | | |
| Assembled pseudochromosomes | 21 | 21 | 21 |
| **Genome annotation** | | | |
| Gene number | 23,981 | - | 25,104 |
| Average transcript length (bp) | 6,421.71 | - | 7,695.22 |
| Average CDS length (bp) | 1,626.98 | - | 1,500.39 |
| Average exon per gene | 8.9 | - | 8.9 |
| Average exon length (bp) | 182.65 | - | 168.67 |
| Average intron length (bp) | 606.37 | - | 784.59 |
| **BUSCO genome assessment** | | | |
| Complete | 95.4% (4,375/4,584) | - | 97.5% (2,522/2,586) |
| Single-copy complete | 92.3% (4,232/4,584) | - | 96.1 (2,486/2,586) |
| Duplicated complete | 3.1% (142/4,584) | - | 1.4% (36/2,586) |
| Fragmented | 2.8% (128/4,584) | - | 1.4% (36/2,586) |
| Missing | 1.8% (82/4,584) | - | 1.1% (28/2,586) |

Reference 1 and 2 are assemblies generated by BGI (*56*) and Oxford (*57*). Reference 3 is the assembly built by this study. BUSCO for ASM365015v1 used Actinopterygii_odb9 as the reference lineage. BUSCO in this study used vertebrata_odb9 as the reference lineage. CEGMA assessment assembled 239 (238 complete and 1 partial) out of 248 CEGs (core eukaryotic genes) in the Siamese fighting fish lineage.

#### Table S2. Sequencing platforms and data output

| Platform | | Insertion (bp) | Data (Gb) | Depth |
|---|---|---|---|---|
| PacBio | SMRTbell | 20 kb | 53.27 | 109.18× |
| Illumina | HiSeq X Ten | 350/150 | 53.55 | 109.76× |
| 10X genomics | | 350/150 | 114.98 | 235.68× |
| BioNano | | -- | 155.66 | 319.05× |
| Hi-C | | 350/150 | 52.58 | 107.77× |

**Table S3. Repeat annotation of the Siamese fighting fish genome**

| Type | Repeat Size (bp) | % of genome |
|---|---|---|
| **Trf** | 34,495,406 | 7.64 |
| **Repeatmasker** | 111,386,726 | 24.68 |
| **Proteinmask** | 21,751,452 | 4.82 |
| **Total** | 119,389,200 | 26.45 |

**Table S4. Transposable element annotation of the Siamese fighting fish genome**

| | Denovo+Repbase Length(bp) | % in Genome | TE proteins Length(bp) | % in Genome | Combined TEs Length(bp) | % in Genome |
|---|---|---|---|---|---|---|
| **DNA** | 26,631,246 | 5.90 | 1,739,683 | 0.39 | 27,825,348 | 6.17 |
| **LINE** | 36,099,475 | 8.00 | 14,184,228 | 3.14 | 39,930,260 | 8.85 |
| **SINE** | 3,488,471 | 0.77 | 0 | 0 | 3,488,471 | 0.77 |
| **LTR** | 34,288,688 | 7.60 | 5,868,612 | 1.30 | 35,407,679 | 7.85 |
| **Simple repeat** | 19,853,701 | 4.40 | 0 | 0 | 19,853,701 | 4.40 |
| **Unknown** | 2,152,019 | 0.48 | 0 | 0 | 2,152,019 | 0.48 |
| **Total** | 111,386,726 | 24.68 | 21,751,452 | 4.82 | 113,804,129 | 25.22 |

#### Table S5. Gene annotation of the Siamese fighting fish genome

| | Gene set | Number | Average transcript length (bp) | Average CDS length (bp) | Average exons per gene | Average exon length (bp) | Average intron length (bp) |
|---|---|---|---|---|---|---|---|
| De novo | **Augustus** | 24,391 | 6,165.89 | 1,448.60 | 8.23 | 176.11 | 652.86 |
| | **GlimmerHMM** | 116,146 | 3,153.40 | 606.86 | 3.27 | 185.51 | 1,121.14 |
| | **SNAP** | 45,506 | 4,805.18 | 914.46 | 5.73 | 159.54 | 822.22 |
| | **Geneid** | 25,185 | 11,273.55 | 1,357.08 | 8.36 | 162.41 | 1,348.09 |
| | **Genscan** | 25,854 | 12,073.09 | 1,744.03 | 9.65 | 180.82 | 1,194.78 |
| | **Cca** | 22,661 | 5,271.45 | 1,300.62 | 6.83 | 190.44 | 681.14 |
| | **Cse** | 20,221 | 7,410.23 | 1,633.96 | 8.82 | 185.36 | 739.10 |
| | **Dre** | 20,403 | 6,869.51 | 1,560.61 | 8.30 | 187.99 | 727.10 |
| | **Gac** | 23,876 | 5,723.22 | 1,329.39 | 7.41 | 179.29 | 684.96 |
| | **Ipu** | 22,203 | 6,522.39 | 1,465.08 | 7.79 | 188.15 | 745.17 |
| | **Loc** | 20,327 | 6,875.17 | 1,474.38 | 8.02 | 183.88 | 769.55 |
| Homolog | **Ola** | 21,838 | 6,915.09 | 1,641.71 | 8.51 | 192.91 | 702.16 |
| | **Oni** | 21,398 | 7,214.29 | 1,656.79 | 8.81 | 188.14 | 711.93 |
| | **Pki** | 21,771 | 6,590.64 | 1,478.89 | 7.92 | 186.79 | 738.98 |
| | **Srh** | 21,101 | 6,682.92 | 1,513.33 | 8.03 | 188.55 | 735.77 |
| | **Ssa** | 24,121 | 5,977.32 | 1,427.03 | 7.40 | 192.74 | 710.57 |
| | **Tru** | 20,568 | 7,037.84 | 1,582.11 | 8.73 | 181.13 | 705.36 |
| | **Xma** | 21,285 | 7,431.05 | 1,689.68 | 8.91 | 189.67 | 725.98 |
| RNAseq | **PASA** | 183,242 | 5,808.30 | 1,149.68 | 7.11 | 161.63 | 762.08 |
| | **Cufflinks** | 70,695 | 13,208.26 | 3,448.98 | 10.88 | 317.01 | 987.81 |
| **EVM** | | 28,224 | 7,098.17 | 1,403.07 | 8.15 | 172.09 | 796.16 |
| **Pasa-update** | | 28,064 | 7,146.26 | 1,414.99 | 8.23 | 171.92 | 792.65 |
| **Final set** | | 25,104 | 7,695.22 | 1,500.39 | 8.90 | 168.67 | 784.59 |

#### Table S6. Non-coding RNA annotation of the Siamese fighting fish genome

| | Type | Copy(w*) | Average length(bp) | Total length(bp) | % of genome |
|---|---|---|---|---|---|
| **miRNA** | | 1,818 | 88.74 | 161,327 | 0.035748 |
| **tRNA** | | 6,214 | 75.02 | 466,163 | 0.10 |
| **rRNA** | **rRNA** | 1,439 | 129.18 | 185,888 | 0.041190 |
| | **18S** | 21 | 537.81 | 11,294 | 0.002503 |
| | **28S** | 110 | 237.87 | 26,166 | 0.005798 |
| | **5.8S** | 7 | 157 | 1,099 | 0.000244 |
| | **5S** | 1,301 | 113.24 | 147,329 | 0.032646 |
| **snRNA** | **snRNA** | 498 | 127.66 | 63,573 | 0.014087 |
| | **CD-box** | 87 | 92.36 | 8,035 | 0.001780 |
| | **HACA-box** | 76 | 156.71 | 11,910 | 0.002639 |
| | **splicing** | 319 | 126.80 | 40,449 | 0.008963 |

**Table S7. Statistics of functional annotation of structural genes in the Siamese fighting fish genome**

| | Number | Percent (%) |
|---|---|---|
| Total | 25,104 | - |
| Swissprot | 21,132 | 84.20 |
| Nr | 13,459 | 53.60 |
| KEGG | 19,127 | 76.20 |
| InterPro | 21,506 | 85.70 |
| GO | 16,079 | 64.00 |
| Pfam | 19,521 | 77.80 |
| Annotated | 22,788 | 90.80 |
| Unannotated | 2,316 | 9.20 |

#### Table S8. Description of all samples included in this study

| Breed | Abbreviation | Brief description | Num. | Illustration |
|---|---|---|---|---|
| Wild Siamese fighting fish | | Wild caught *Betta splendens* | 11 | Figure S5s |
| *Betta imbellis* | | Wild caught *Betta imbellis* | 9 | Figure S5t |
| *Betta siamorientalis* | | Wild caught *Betta siamorientalis* | 9 | Figure S5u |
| *Betta mahachaiensis* | | Wild caught *Betta mahachaiensis* | 11 | Figure S5v |
| *Betta smaragdina* | | Wild caught *Betta smaragdina* | 11 | Figure S5w |
| *Betta smaragdina guitar* | | Wild caught *Betta smaragdina guitar* | 4 | Figure S5x |
| *Betta stiktos* | | Wild caught *Betta stiktos* | 4 | Figure S5y |
| Fighter Siamese fighting fish | Fighter | Short fin, selected for more aggressive performances over centuries | 101 | Figure S5r |
| Veiltail Siamese fighting fish | Veiltail | Long fin, Veil-shaped caudal fin | 61 | Figure S5q |
| Crowntail Siamese fighting fish | Crowntail | Long fin, Crown-like appearance of the caudal fin, reduction in webbing/tissue between all fin rays | 55 | Figure S5p |
| Halfmoon Siamese fighting fish | Halfmoon | Long fin, symmetrical D-shaped caudal fin | 85 | Figure S5o |
| Halfmoon Dumbo Siamese fighting fish | Halfmoon_Dumbo | Long fin, symmetrical D-shaped caudal fin and enlarged pectoral fins | 38 | Figure S5n |
| Halfmoon Plakat Siamese fighting fish | HMPK | Short fin, symmetrical D-shaped caudal fin | 424 | Figure S5a-m |
| Giant Siamese fighting fish | Giant | Variation with a larger body size than normal ones; SL: male > 4.5 cm, female > 4 cm | 102 | Figure S5k |
| HMPK Dumbo Siamese fighting fish | HMPK-Dumbo | Enlarged and extended paired pectoral fins | 64 | Figure S5m |
| Solid Red Siamese fighting fish | Red | Solid red on the fish appearance | 26 | Figure S5a |
| Solid Yellow Siamese fighting fish | Yellow | Solid yellow on the fish appearance | 29 | Figure S5b |
| Solid Orange Siamese fighting fish | Orange | Solid orange on the fish appearance | 28 | Figure S5c |
| Solid Steel Blue Siamese fighting fish | Steel | Solid Steel Blue on the fish appearance | 35 | Figure S5f |
| Solid Royal Blue Siamese fighting fish | Royal | Solid Royal Blue on the fish appearance | 28 | Figure S5e |
| Solid Turquoise green Siamese fighting fish | Turquoise | Solid Turquoise green on the fish appearance | 26 | Figure S5d |
| Solid Black Siamese fighting fish | Black | Solid Black on the fish appearance | 20 | Figure S5g |
| Solid Opaque Siamese fighting fish | Opaque | Solid Opaque on the fish appearance | 17 | Figure S5h |
| Copper Siamese fighting fish | Copper | Solid Copper (homozygous metallic steel blue) on the fish appearance | 43 | Figure S5i |
| Dragon Siamese fighting fish | Dragon | Solid Dragon (black, red, orange) on the fish appearance | 28 | Figure S5j |
| Koi/Marble Siamese fighting fish | Koi/Marble | Different color exhibits a mosaic pattern on the fish appearance | 55 | Figure S5l |

**Table S9. ABBA BABA tests for introgression between B. mahachaiensis and Turquoise, Steel blue, Royal blue and Copper breeds of *Betta splendens.*** The tests are in the form of D(W, X; mahachaiensis, outgroup) where the outgroup is a *B. smaragdina* individual and W are Turquoise, Steel blue, Royal blue and Copper individuals and X are individuals of all other breeds.

| Tests | Z scores |
|---|---|
| D(Turquoise, X; mahachaiensis, outgroup) | range: -28 / 7.4, median: -15 |
| D(Royal, X; mahachaiensis, outgroup) | range: -26 / 10, median: -13 |
| D(Steel, X; mahachaiensis, outgroup) | range: -27 / 5, median: -16 |
| D(Copper, X; mahachaiensis, outgroup) | range: -26 / 12, median: -11 |

**Table S10. Genotype frequencies at top sex-associated SNP (chr2:2839325) in males and females**

| | Domesticated | | Wild | | | | | | |
|---|---|---|---|---|---|---|---|---|---|
| | Male | Female | BSP | BIM | BSS | BMA | BSM | BST | BSG |
| | 590 | 137 | 11 | 9 (3♂, 6♀) | 9 (4♂, 5♀) | 11 (5♂, 6♀) | 11 (6♂, 5♀) | 4 (2♂, 2♀) | 4 (2♂, 2♀) |
| CC | 8 | 0 | 0 | 9 | 9 | 11 | 11 | 4 | 4 |
| CT | 530 | 7 | 3♂ | 0 | 0 | 0 | 0 | 0 | 0 |
| TT | 52 | 130 | 8♀ | 0 | 0 | 0 | 0 | 0 | 0 |

BSP, wild betta splendens; BIM, betta imbellis; BSS, betta siamorientalis; BMA, betta mahachaiensis; BSM, betta smaragdina; BST, betta stiktos; BSG, betta smaragdina guitar.

**Table S12. GWAS summary for Mosaic pattern**

| Chr | TopSnp position | Top Snp P Value | Range | SNP numbers | Gene numbers | Candidate Genes |
|---|---|---|---|---|---|---|
| 2 | 10192063 | 2.31E-13 | chr2:9826395-10688853 | 189 | 39 | SLC2A11, Slc8b1, AIDB |
| 3 | 4276186 | 1.25E-11 | chr3:4153497-5856276 | 71 | 18 | BCO1, SLC22A13, DYTN |
| 10 | 2442406 | 6.56E-20 | chr10:1661045-2640841 | 66 | 13 | HCEA, SLC23A2 |
| 13 | 5799294 | 4.64E-11 | chr13:5794871-5837251 | 33 | 5 | Amigo1, Gpr61 |
| 14 | 3649208 | 1.78E-15 | chr14:3527165-3687639 | 50 | 9 | tmem151b, RPAP1 |
| 16_1 | Multiple | 1.93E-57 | chr16:22641-941275 | 1517 | 39 | Slc39a7, CHS3, CHS8, COL11A2, tubb |
| 16_2 | 2643502 | 3.39E-40 | chr16:1762170-2981446 | 1182 | 47 | PLEC, EPPK1, SLC17A5, SLC52A2 |
| 20 | 2720181 | 4.19E-13 | chr20:2331820-2721015 | 187 | 20 | TRIM67, esco2 |
| 21 | 3481675 | 6.31E-12 | chr21:3477277-4558869 | 47 | 18 | CAMKK2, slc20a1b |

**Table S13. Predicted miRNA binding to *KCNJ15* using RNAhybrid and miRanda**

| Target::miRNA | RNAhybrid | miRanda |
|---|---|---|
| KCNJ15::dre-let-7b | 1 | 0 |
| KCNJ15::dre-miR-138-3p | 1 | 0 |
| KCNJ15::dre-miR-1388-3p | 1 | 0 |
| KCNJ15::dre-miR-143 | 0 | 1 |
| KCNJ15::dre-miR-205-3p | 0 | 1 |
| KCNJ15::dre-miR-21 | 1 | 0 |
| KCNJ15::dre-miR-222a-3p | 1 | 0 |
| KCNJ15::dre-miR-22a-3p | 1 | 0 |
| KCNJ15::dre-miR-23a-3p | 0 | 1 |
| KCNJ15::dre-miR-23b | 0 | 1 |
| KCNJ15::dre-miR-34a | 1 | 0 |
| KCNJ15::novel_11 | 1 | 0 |
| KCNJ15::novel_149 | 1 | 0 |
| KCNJ15::novel_159 | 0 | 1 |
| KCNJ15::novel_236 | 1 | 0 |
| KCNJ15::novel_240 | 0 | 1 |
| KCNJ15::novel_243 | 1 | 0 |
| KCNJ15::novel_273 | 1 | 0 |
| KCNJ15::novel_343 | 0 | 1 |
| KCNJ15::novel_368 | 1 | 0 |
| KCNJ15::novel_372 | 0 | 1 |
| KCNJ15::novel_90 | 0 | 1 |

### Supplementary Text

#### Supplementary Text 1. Background on Betta fish

The common name Betta fish, refers to the wild and domesticated fish in the genus *Betta* distributed in Southeast Asia, especially in Thailand, Cambodia, Vietnam, Malaysia and Indonesia. The Siamese fighting fish, *Betta splendens*, is primarily located in Central Thailand and the lower Mekong, is well-known for its aggressive behavior and various domesticated forms (*58*). This fish was traditionally used in gambling matches and has been domesticated for at least 300 years (*58*). Today, multiple varieties have been raised differing in aggressiveness, fin shape, color morphology and body size. Already in the 1930s and 1940s, the inheritance of several traits such as sex-determination, long-fin vs short-fin, and coloration of Turquoise green, Royal blue and Steel blue colors, were explored (*59*-*63*). However, there have been relatively few modern studies on the genetics of *B. splendens* domestication traits, until a recent study of the double tail, elephant ear, albino and fin spot phenotypes (*64*). In addition to the interest in its phenotypic diversity, the Siamese fighting fish is also a popular model for toxicological and behavioral research.

Although initially domesticated for its aggressive behavior, numerous other breeds have been domesticated for ornamental purposes. In total, we have sampled five major caudal fin morphology types: the Fighters and four types of non-Fighters, namely Crowntail, Veiltail, Halfmoon and Halfmoon Plakat (HMPK). HMPK has a characteristic shorter tail, while Crowntail, Veiltail and Halfmoon all have distinctive long tails. Here we have collected 13 breeds of HMPK differing mainly in color, but also in size (the Giant breed being larger than the others) and pectoral fin morphology (the Dumbo breed having enlarged pectoral fins) (Figs. S5a-m).

We note here that although we are using the term “breed” in this section and throughout the paper, these are not breeds in the traditional sense of inbred lines, since these morphotypes are often interbred. One evidence of that is the Royal blue phenotype, which when bred to itself produces Turquoise green and Steel blue offspring (*60*), showing that a segregating locus underlies all those phenotypes. Another example is the Dumbo phenotype, which exists in both Betta fish with HMPK and Halfmoon tail types.

Being aware that what we are calling “breeds” are akin to morphotypes, we have sought to assign individuals to breeds not only based on phenotypic similarity, but also considering the potential for a shared genetic basis for the phenotype based on past breeding experiments.

**Supplementary Text 2: Genome assembly and genome annotation results**

Genome assembly

In this section we describe the results of our genome assembly. For a description of the methods please see Materials and Methods.

A female solid red HMPK individual was selected for genome assembly. The genome size was estimated to be 487.89 Mb using Jellyfish (v2.0)(*65*) with a heterozygosity of 0.27% determined by k-mer (k = 17) frequency analysis using 43.51 Gb Illumina sequencing data. The genome was assembled using 53.27Gb (~110x) PacBio Sequel SMRTbell data, augmented with 10X genomics, BioNano optimal map and Hi-C data for scaffolding and anchoring to chromosomes (see Materials and Methods). The final genome assembly is 451.29 Mb with a scaffold N50 of 19.63 Mb (Table S1), showing substantial improvements in continuity compared to previously published *B. splendens* genomes (Table S1). To evaluate the quality of our assembly, we mapped the Illumina short reads to the assembled genome using BWA (*38*), and 97.51% of reads were successfully mapped. 99.87% of the assembly was covered, indicating the near-completion of the genome. Filtered variants called from Illumina data yielded 198,849 heterozygous SNPs (0.0936%) and 532 homozygous SNPs (0.0003%), supporting high base level accuracy of the genome assembly.

Genome annotation

The final assembly was annotated for protein coding (Fig. S1, Tables S5, S7) and non-coding RNA (Fig. S1, Table S6), repeat sequences and transposable elements (Fig. S2, Tables S3 and S4).

We annotated structural genes, repeat sequences, and non-coding RNAs in the assembled genome, which are provided as resources for future research. A list of 25,104 structural gene models was predicted, and 22,788 (90.77%) of those were functionally annotated using a customized pipeline including both *de novo* and reference guided transcriptome assembly, and similarity based prediction of function (Supplementary Text 2 and Materials and methods). The Siamese fighting fish genome has 119.38 Mb repeat sequence (26.45% of the genome), which is lower than the zebrafish (52%) (*66*)and medaka (33%) (*67*). The repeat sequences consist of 19.85 Mb tandem repeats and 100.51 Mb transposable elements (TE) (Fig. S2 and Table S4). Class I (Retrotransposons) and classII (DNA transposons) TEs compose 17.47% and 6.17% of the assembly, respectively (Table S4). The long interspersed nuclear elements (LINE) retrotransposons are the most abundant of TEs (8.85% of the genome) (Table S4). We also identified sequences of 11,890 non-coding RNA elements across the genome, including 1,818 microRNAs, 6,214 transfer RNAs, 2,878 ribosomal RNAs and 980 small-nuclear RNAs (Fig. S1 and Table S6).

**Supplementary Text 3: Phylogeny and expanded and contracted gene families.**

Phylogeny for *B. splendens*

A maximum-likelihood phylogeny estimated using 465 commonly shared single-copy orthologs among 14 teleost fishes support the hypothesis that the Siamese fighting fish (*Betta splendens*) is a sister clade to Perciformes, including platyfish, medaka, Nile tilapia, puffer and stickleback (Fig. S3). The time-calibrated phylogeny is overall consistent with the one described in (*68*), with *Betta splendens* diverging ~109.6 million years ago (MYA) from Perciformes.

The gene-family expansion/contraction analyses in CAFE (*37*) on the *B. splendens* lineage identified 75 gene families consisting of 384 genes (1.53% of the total genes) expanding and 79 gene families consisting of 101 genes (0.40% of the total genes) contracting (Figs. S3, S4).

Expanded gene families related to muscular, neuronal and solute carrier functions

Several genes in the expanded families in the *B. splendens* lineage are generally involved in a variety of cytoskeleton related activities, including muscle fiber components, neuromuscular junction and muscle contraction activities (Fig. S4, Supplementary Data 3).

Among the gene families encoding components of muscle fiber are *MYO* and *TTN*. *MYOs* produces the thick filament (myosin), which connects with elastic filament (titin) encoded by *TTN*, and functions together with thick filament (actin) in a muscle fiber to generate muscle contraction [(*69*). *DNAH* (dynein axonemal heavy chain, 10 paralogs), *ANK* (ankyrin, 8 paralogs), *PLEC* (plectin, 8 paralogs), *MACF* (microtubule actin crosslinking factor, 7 paralogs), *SYNE* (spectrin repeat containing nuclear envelope protein, 11 paralogs) families (Supplementary Data 3) also encode components of muscle fiber. *DNAH* genes encode a group of proteins that move along microtubules (*70*). *ANKs* encode proteins which are involved in the attachment of integral membrane proteins to the spectrin-actin based membrane cytoskeleton (*71*). Additionally, the ankyrin 3 gene (*ANK3*) is also involved in neuronal activity. *ANK3* has been linked to bipolar disorder and schizophrenia and showed association with externalizing behavior in humans (*72*). Another gene related to neuronal activity is *NRXN1*, which encodes a cell membrane protein that regulates calcium, triggers neurotransmitter release and has also been associated with schizophrenia as well as developmental disorders (*73*).

In addition, 116 genes among the expanded families are involved in processes related to the neuromuscular junction, such as *MACF1* (microtubule actin crosslinking factor 1), *FAT1* (FAT atypical cadherin 1) and *SCN8A* (sodium voltage-gated channel alpha subunit 8) (Supplementary Data 3).

One of the most noticeable activities performed by the cytoskeleton is muscle contraction. An expansion of gene families related to cytoskeleton and muscular function in the Siamese fighting fish could be driven by the fact that these fish regularly exhibit a high muscular tension state when facing opponents, resulting in sustained energy consumption. When facing an opponent, the Siamese fighting fish will display its distinctive aggressiveness by tightening its muscles and wiggling its body frequently, possibly followed by direct fighting. The over-representation of genes related to cytoskeleton and muscular biological processes among the expanded gene families may indicate the underlying genetic basis for the aggressive behavior of the Siamese fighting fish relative to other species.

Another type of expanded gene family in *B. splendens* is a solute carrier. We mentioned in the main text that *SLC4A* (solute carrier family 4 anion exchanger) families have experienced remarkable expansions (Supplementary Data 3) in the Siamese fighting fish. *SLC4As* encode proteins that act as bicarbonate transporters that move $HCO3^-$ across the plasma membrane and regulate intracellular pH and transepithelial movement of acid-base equivalents (*69*). These 18 SLC4A paralogs are demonstrated to play an important physiological role in maintaining acid-base homeostasis at the cellular and systemic levels (*69*). We postulate three types of selective pressures that could drive the multiplication of *SLC4A* genes. First, as one of the Anabantoids, the Siamese fighting fish is a flagship species that inhabits peat swamp waters, where pH could be as low as 3-4. The acidic water causes the penetration of $H^+$ and loss of $HCO_3^-$ from the gill and kidney of the Siamese fighting fish. Second, excessive fighting would increase anaerobic metabolism, which produces intermediate components such as lactate, which reacts with bicarbonate. Third, many species in Anabantoids including the *B. splendens* are facultative air breathers, since they do not have sufficient gill function to fully support their oxygen needs. Thus, digestion of the acid-base equivalents under these circumstances would sustain its survival in natural habitats.

Gene loss related to ion exchange pathway

The SLC6A (solute carrier 6 alpha) is the gene family showing the strongest contraction in the Siamese fighting fish. This family is widely engaged in the transmembrane carriage of serotonin, dopamine, GABA, norepinephrine and amino acids (*74*). The transport of neurotransmitters is potentially related to the characteristic aggressive behaviour of *B. splendens*. However, the relationship between the function of this and other contracted gene families and potential selective pressures on the Siamese fighting fish (such as aggressive behaviour and adaptation to hypoxic and acidic black water conditions) remains to be investigated.

**Supplementary Text 4: Phenotyping**

Morphology

Body Size: The giant and non-giant individuals were identified visually. Body weight was measured with a digital scale and body height, standard length, total length and tail length of each fish were measured with digital callipers.

Color and sex: The body color, eye color, fin color, fin shape and sex were phenotyped visually based on photographs taken for each fish, and the photos of each fish were deposited in Dropbox (https://www.dropbox.com/sh/kr3jub2b7wiplbe/AAD-1ipXdvI5ovMFThJVKuRGa?dl=0).

Fin rays: Three phenotypes of fin rays for dorsal and caudal fins were obtained by visual examination: (1) number of rays in fins; (2) maximum number of branches of all fin rays; (3) Unanimous branching, *i.e.* whether all the rays in dorsal and caudal fins share the same number of branchings or not (coded as a binary trait).

Aggressive behaviours

To quantify aggressiveness, we set up a simulative fighting experiment and recorded the behaviours of the test fish with a sports camera for one minute. The experiment is performed in a fish tank equally divided into halves using a transparent glass plate (Fig. 4B). An opponent fish is put on one side of the tank, and the test fish is put on the other side of the tank. Before simulative fighting, an opaque plate was inserted between the two fishes, and all fishes were acclimated to the new tank for at least 10 and up to 20 minutes. We excluded those individuals which did not seem to be acclimated to the new environment, based on observation of their behaviour, *i.e.* if the fishes were standing still or behaved nervously in the new tank. Then, the opaque plate was removed and the behaviours of the test fish were recorded for one minute. In total, 467 individuals were recorded by video. Based on these videos, the aggressiveness of each fish is scored on a 0-5 range on nine indices: charge, mouth open, gill flare, fin flare, shimmer, jerk, approach, pacing, and retreat, which are explained in detail below. The number of times air-breathing within one minute was also counted for each fish. Based on the nine behaviours recorded, we assign an overall aggressiveness rating named “aggression index”, with the equation: (Charge×5 + Mouth open×5 + Gill flare×4 + Fin flare×3 + Shimmer×2 + Jerk×2 + Approach×2 + Pacing×1 - Retreat×1). The nine indices composing the aggression index are defined as:

Charge: swims towards transparent barrier rapidly and repeatedly (Fig. 4H)

Mouth open: opens mouth and locks jaw not for eating purposes (Fig. 4C)

Gill flare: operculum flare under extension (Fig. 4D)

Fin flare: raises the dorsal fin (Fig. 4E)

Shimmer: displays one side of the body with shimmering colour within one inch of the opponent and (Fig. S42B)

Jerk: rapid twitching movement resulting in direction change (Fig. S42A)

Approach: swims in the direction of the opponent (Fig. 4G)

Pacing: circular swimming around the edge of the tank (Fig. S42C)

Retreat: swims in the direction opposite to the opponent (Fig. 4F)

**Supplementary Text 5: An overview of GWAS setups**

Body size

Two types of GWAS for body size were performed: (1) case-control, giant (n=101) versus non-giant breeds (n=508) (Fig. 2C, S33A); (2) quantitative phenotypes including body weight (Fig. S32D, S33D), body height (Fig. S32E, S33E), standard length (Fig. 2C, S32C, S33C), total length (Fig. S32B, S33B) and tail length of each fish (n=609), Body mass index (BMI), Total length/Height, Weight/Height (n=608) (Fig. S49).

We performed GWAS for body size both including and excluding the Fighter breed, as body size is highly variable even only within non-fighter breeds. GWAS signal was weaker when including the Fighter breed (Fig. S54), but the same loci that were significantly associated with body size in the non-fighter GWAS showed weak signal when including the Fighters. We hypothesize that different loci might be involved in the genetic basis of body size in Fighters and non-fighters, and we focus on the significant signal found in non-fighters.

Color

For color, we performed (1) case-control GWAS using different combinations of colors as cases and controls, as well as (2) quantitative GWAS coding color as a discrete variable with multiple levels.

For the Yellow, Orange and Red color breeds, we performed three sets of case-control GWAS (Fig. S33), using each color as cases against the other two as controls: Orange vs (Red+Yellow), Yellow vs (Red+Orange), Red vs (Yellow+Orange). We also did one case-control GWAS of Orange vs Yellow (Fig. S33).

For the Turquoise green, Steel blue and Royal blue color breeds, we used the following setups for case-control GWAS: (Turquoise green+Steel blue) vs Royal blue, Turquoise green vs Steel blue (Fig. S35). Additional case-control GWAS were done for the following color patterns: Copper vs Non-Copper, Copper in HMPK vs Non-Copper in HMPK, Dragon vs Non-Dragon, Dragon in HMPK vs Non-Dragon in HMPK, Steel blue vs Copper (Fig. S50), Solid color (Red, Yellow, Orange, Turquoise green, Royal blue, Steel blue, Black, Opaque, Copper and Dragon) vs Mosaic color (Koi), Iridescent (Steel blue, Royal blue, Turquoise green, Opaque) vs Non-Iridescent (Red, Yellow, Orange, Black).

Finally, we performed GWAS coding colors as discrete quantitative variables (Fig. 2B, S34, S36). The Royal-blue is a hybrid between Turquoise-green and Steel-blue, and Orange is a hybrid between Red and Yellow. Therefore, we coded the hybrid phenotypes Royal blue and Orange as 2, and the two parental phenotypes from each hybrid as 1 and 3.

In addition to the body color, we also performed GWAS for eye color. We classified eye color into six categories: black, white, yellowish-brown, yellow, light blue and brown and performed GWAS coding these phenotypes as categorical variables.

Fins

For fin-related phenotypes, we also performed both case-control and quantitative GWAS. The case-control setups for tail type were: Short tail (HMPK+Fighter) vs Long tail (Veiltail+Halfmoon+Crowntail) (Figs. 3A, S23, S24), Halfmoon vs Crowntail (Figs. 3H, S27), Halfmoon vs Veiltail (Figs. 3G, S26), Crowntail vs Veiltail (Figs. 3I, S28).

We also performed a case-control GWAS for fin color, constructed using (body and fin sharing the same color) vs (body and fin showing different colors).

For fin rays, we did two case-control GWAS: one comparing fish for which all the rays in the dorsal fins share the same number of branching to fish where they do not (Fig. S32E), and a second one for the same trait in the caudal fin (Fig. S32F). We also did the following quantitative GWAS for fin rays: 1) Number of dorsal (Fig. S32A) and 2) caudal fin rays (Fig. S32B), maximum number of branches of 3) dorsal (Fig. S32C) and 4) caudal fin rays (Fig. S32D).

Finally, we performed conditional GWAS, in which the genotype of the most significant variant for each phenotype was treated as covariate, i.e. a conditional GWAS to find associations controlling for the effect of the lead SNP (Fig. S36, S55).

**Supplementary Text 6. Population genetics of *Betta splendens***

As described in the Methods, we estimated a Neighboring-Joining (NJ) phylogeny (Figure 1B) using concatenated sequences. We note that, because of the coalescent process, different regions of a genome will have different trees. The NJ tree can be considered an estimate of the average tree across the genome. While it is important to avoid confusion of the interpretation of the branch lengths in such a tree, it is useful for elucidating individuals, and groups, that in average are more closely related to each other genetically than other such groups. In our NJ tree, all individuals from *Betta splendens* species form a clade relative to other wild species, providing genetic evidence for a single origin of all domesticated Siamese fighting fish from the wild *B. splendens*, despite their extraordinarily high phenotypic diversity. This result is also compatible with the PCA in which all *B. splendens* individuals form a cluster (Fig 1C). The fighter individuals are located in multiple different clades at the basal position of the tree, suggesting that the domesticated *B. splendens* was first selected for fighting, and other breeds featured for color or fin morphology diversified later during domestication.

The population structure of the domesticated *B. splendens* is largely driven by several breeds that carry substantial breed-specific genetic drift, including the Black, Orange and Yellow, the clade consisting of Steel-blue, Royal-blue and Turquoise-green, and Opaque (Figure 1B-D). These breeds have long breed-specific branch lengths on the NJ tree (Fig. 1B) and show distinctive components in the *ADMIXTURE* analysis (Fig 1D). The most noticeable is the group consisting of Steel-blue, Royal-blue and Turquoise-green, as they form an independent cluster in the PCA analysis (Figure 1C). This breed-specific genetic drift is likely due to the strong bottleneck effect imposed by local breeders.

The Dumbo Halfmoon breed is featured for both its distinctive enlarged pectoral fin (the dumbo phenotype) and its semicircular tail that is similar to the Halfmoon breed, but its genetic origin remains elusive. In the genome-wide average phylogeny (Figure 1B) it clusters with Dumbo-HMPK rather than Halfmoon, which is also supported by the *ADMIXTURE* result as the two Dumbo breeds share the same genetic components that is distinctive from the Halfmoon breed (Figs. 1D, S17). These results suggest that the Dumbo Halfmoon obtained the long caudal-fin phenotype through introgression from the Halfmoon breed.

Treemix analyses and potential gene-flow

Threemix analyses (Figs. S7-S15) suggest extensive gene-flow between wild species and domesticated breeds of *B. splendens*. Allowing one admixture edge (Fig. S8) infers an admixture event from the ancestor of the (*B. splendens, B. imbellis, B. siamorentalis*) group into wild *B. splendens*. Allowing two edges (Fig. S9) infers an additional edge from the ancestor of B. *splendens* into the Giant breed. Allowing 3 edges (Fig. S10) adds an admixture event into the Steel blue from *B. Mahachaiensis.* With four admixture events allowed, a new migration edge from the ancestor of *B. siamorientalis* and *B. imbellis* into the Dragon breed is added. We find the possibility of gene-flow between *B. Mahachaiensis* and some *B. splendens* breeds particularly interesting because of the iridescent scales on *B. Mahachaiensis* shared with some domesticated *B. splendens* breeds such as the Steel blue breed. We, therefore, investigate this hypothesis further using ABBA-BABA tests.

To explore gene flow between *B. mahachaiensis* and Siamese fighting fish, we exhaustively tested all possible combinations of D-statistics in the form of D(H1, H2; mahachaiensis, outgroup) where H1 and H2 are all possible combinations of different Siamese fighting fish breeds. If there were no gene flow between *B. mahachaiensis* and any of the Siamese fighting fish breeds, we would expect D ~ 0. However, the tests strongly deviate from the no-gene-flow expectation (Fig. S16), suggesting frequent gene flow between them. Moreover, the most significant gene flow signals are found in Turquoise green, royal blue, steel blue and copper. which all have excess allele sharing with *B. mahachaiensis* when compared with other breeds (Fig. S16). These results are highly consistent with the TreeMix analyses, providing strong evidences for gene flow between *B. mahachaiensis* and the Turquoise green, Royal blue, Steel blue and Copper breeds, with the TreeMix analyses suggesting that the direction of gene flow is mainly from the wild *B. mahachaiensis* into the domesticated species.

**Supplementary Text 7: GWAS of sex determination**

In the GWAS of sex determination, we detected a single locus on chromosome 2 that is highly associated with the phenotype (Fig. 2A). The locus was replicated when dividing the whole cohort into different breeds, including Fighter (77 males vs 24 females), Crowntail (33 males vs 22 females), HMPK (364 males vs 60 females) and Veiltail (35 males and 26 females) to conduct GWAS, suggesting all breeds share the same sex-determining locus (Fig. S55).

To track down the causal mutations, we first ask whether there is structural variation involved, as structural variations often occur around the sex-determining region of fish, like *DMY* in medaka (*75*) and *AMY* in Nile tilapia (*76*), initiating the process of differentiation of sex chromosomes. We first leveraged the published high quality genome assembly of a male which has XY chromosome and compared it with our genome assembly which is from an XX female. When aligned our female-based sex-determining (SD) region (chr2:2.8-2.9 Mb) to the corresponding region (chr9:28.8-28.9 Mb) in another male-based assembly using LastZ (*77*)(Fig. S20), a complete syntenic block is observed synteny between the males and females at the sex determination locus. This result suggests that there is no structural variation underlying the sex determination, but it is also possible that an alternative haplotype with structural variation is missed during genome assembly. To further investigate this, we aligned the sequencing reads of male and female individuals separately to the female-based (XX) assembly and calculated the read depth within this region (Figs. S21C and D). No difference was observed in the average sequencing depth between males and females, with GC content along this region showing no disparity (Fig. S21). These results collectively suggest that there is no structural variation in the SD region between males and females in the Siamese fighting fish.

Zooming in on the associated locus revealed several genes including *DMRT1* (Fig. 2A). Among these genes, *DMRT1*, *DMRT3* and *CFAP157* are differentially expressed between the adult testis and ovary (Fig. 2A). A cluster of the most significantly associated variants reside in introns of *DMRT1*, with the lead SNP located on the 4th intron of *DMRT1* (Fig. 2A). Taken together, our results strongly support that the underlying causal mutation for sex determination in the Siamese fighting fish is SNPs rather than large structural variations or gene duplications. Further experiments are needed to pinpoint causative mutations, and reveal the full picture of the molecular mechanism for sex-determination in *B. splendens*.

In the main text, we mention that 52 of 590 males have the homogametic genotypes typical of females (Table S10), and that this suggests non-genetic factors may have an effect on sex-determination. In the 11 wild *B. splendens*, the genotype of the lead SNP is perfectly correlated with their sex, suggesting that the wild *B. splendens* and the domesticated breeds share the same XY sex determining locus. However, all 22 male and 26 female samples from the other wild species in the *B. splendens* complex have the same homozygous genotype at this SNP (CC) (Table S10), which suggests that they either have a different sex determination system or that the lead SNP in our study is not causal. However, the *Betta* species may be an interesting study system for rapid evolution of sex determination systems similarly to medaka.

We also compared the genes homologous to *DMRT1* in both *B. splendens* and medaka (Fig. S22). There are five paralogs in *B. splendens* and three paralogs in medaka. The phylogeny involving these genes (Supplementary Figure 24) show that the most related homolog of *DMRT1* in medaka is *DMY* which is the sex-determination gene in medaka. It is surprising that the *DMRT1* orthologs is also co-opted for sex determination, albeit the divergence time between *Betta splendens* and medaka is ~100 million years.

**Supplementary Text 8: Candidate loci in GWAS of fin morphology**

We have explored the genetic basis of various fin morphology traits in the Siamese fighting fish using GWAS (Supplementary Text 5). The non-fighter breeds can generally be classified into two categories, the HMPK breeds with short caudal fin, and the long-caudal-fin breeds including Veiltail, Crowntail and Halfmoon. We first performed GWAS on the long-fin and short-fin phenotype and identified a strong gene candidate, *KCNJ15*. Among the three long-fin breeds, Veiltail, Crowntail and Halfmoon also show great diversity on the shape which we also investigated with GWAS. The 'Dumbo' breed carries a mutated form of the pectoral fin that is enlarged, and we also mapped the genomic loci underlying this phenotype.

The *KCNJ15* gene associated with fin length

The GWAS of long-fin vs. short-fin pointed to three associated variants located in the vicinity of the gene *KCNJ15* (Fig. 3B). A GWAS using caudal fin (tail) length as a quantitative trait identified a significant association at the same locus. *KCNJ15* encodes a potassium channel, and fin overgrowth has been shown to be caused by other genes related to potassium transport, such as *KCNK5B*, *KCNH2A*, *KCNJ13*, *KCC4A* in zebrafish (*78-81*)and *KCNQ5* in goldfish (*82*). Gain-of-function mutations in *KCNK5B*, a gene also engaged as a K+ channel, affect ionic conduction and lead to hyperpolarization of the cell (*78*). Overexpression of *KCNK5B* is sufficient to cause fin overgrowth (*78*). The RNA-seq reads alignment on this region in IGV shows that there were no intact transcripts identified in caudal fins in HMPK (a short-fin breed) (Fig. S25). Instead, in Halfmoon (a long-fin breed), a consensus transcript longer than the annotated structure of *KCNJ15* was detected in all individuals (Fig. S25). The top associated mutation (chr17:9,596,738, G>A) is located in the 3'UTR of the long transcript of *KCNJ15*. To explore a potential mechanism underlying the GWAS signal at this variant, we sequenced caudal fin tissues by miRNA-Seq and compared the miRNA binding sites genome-wide between long-fin and short-fin breeds. There is no strong evidence of microRNA binding sites within or near the *KCNJ15* locus (Table S13), indicating there might be other mechanisms involved. Nevertheless, the *KCNJ15* is a promising candidate gene for fin length variation, and deserves further experimental validation, e.g., phenotypic rescue experiments of the long-fin phenotype using RNA injection.

We also conducted GWAS for fin ray number, maximum number of fin ray branchings and unanimous branching pattern (Fig. S32). No significant association was detected for numbers of dorsal and caudal fin rays and unanimous branching (Fig. S32A, B, E, F). But we did identify a locus at 9.51 - 9.65 Mb on chromosome 20 significantly associated with the maximum number of branchings at both the dorsal and caudal fins (Fig. S32C, D). The top SNP is an exonic mutation (C/T) in *BCL11B*, a zinc finger protein transcription factor with important roles for differentiation and development of various neuronal subtypes in the central nervous system (*83*).

Genomic loci associated with Veiltail, Crowntail and Halfmoon caudal fin type

Through GWAS, we mapped the long-fin versus short-fin locus to a single base mutation in 3'UTR of *KCNJ15*. However, there is much additional phenotypic variation among the long-fin breeds including the Veiltail, Crowntail and Halfmoon phenotypes as some of the most distinctive ones. To study the genetic basis underlying these variations, we performed GWAS on the three long-fin types (see GWAS setup in Supplementary Text 5), and found three independent loci (Fig. 3G-I).

Phenotypically, the most important difference in the caudal fin morphology between Crowntail and Halfmoon breeds is the amount of webbing tissue between fin rays. GWAS using the two breeds as case and control revealed a significant signal at 12.21-12.27 Mb on chromosome 3 (Figs. 3H, S27), A cluster of significantly associated variants is located in the intergenic region between *A0ZSK3* (neoverrucotoxin subunit alpha) and *CEP70* (centrosomal protein 70) (Fig. S27). *A0ZSK3* encodes a potentially lethal secreted toxin with hemolytic activities in nemerteans and fish(*84*). *CEP70* organises both preexisting and nascent microtubules in interphase cells(*85*). The previously described functions of both of these genes do not immediately suggest any clear functional relationship to the phenotypic difference between Crowntail and Halfmoon.

Comparing the Crowntail and Veiltail breeds revealed a locus (8.34-8.39 Mb) on chromosome 14 containing 36 associated variants and eight genes (Figs. 3I, S28). The lead SNP (chr14:8346226) is located near a gene named *HIVEP2*, which encodes a protein binding to the enhancer elements of numerous viral promoters (*85*). One gene in this region, *FRMD6*, is a promising candidate as it is involved in actomyosin structure organization and loss of expression in this gene leads to epithelial-to-mesenchymal transition features (*86*).

GWAS for phenotypic difference between Veiltail and Halfmoon identified a 300-kb long locus on chromosome 8, harboring 29 associated SNPs and four protein-coding genes (Figs. 3G, S26). This region includes four zinc finger encoding genes (*ZNF407*, *ZADH2*, *TSHZ1* and *ZNF516*). Zinc finger proteins are widely involved in transcriptional regulation and cellular functions (*87*). 20 out of the 29 associated variants in this locus that show the strongest association are intergenic (Fig. S26).

Associations with the "Dumbo" phenotype

Performing GWAS on the "Dumbo" phenotype, i.e., the outgrown pectoral fin that is observed in the Dumbo HMPK and Dumbo Halfmoon breeds, we identified two highly significant signals on chromosome 16 (Fig. S29.) and chromosome 18 (Fig. S30) respectively. The signal on chromosome 16 is in the same region that is also mapped by Wang et al. using an $F_{ST}$ scan in a recent study (*64*) (note that our Chr. 16 is labeled Chr. 9 in their study), but the peak identified here in our analysis is narrower and does not contain the gene Wang et al. (*64*)suggested as causal using arguments based on expression patterns. We also leveraged our RNA-Seq dataset from the pectoral fins of five Dumbo HMPK and five HMPK individuals to examine the three candidate genes proposed in Wang et al. (*64*), i.e. *KCNH8*, *EVX1*, *COL16A1*, and *KCNH8* and *EVX1*, did not find evidence of differential expression (Fig. S31), and thus cannot confirm the results in Wang et al.(*64*). However, in the region we identify, there are eight genes of the *HOXA* family, including *Hoxa3a*, *Hoxa4*, *Hoxa5*, *Hoxa7*, *Hoxa9*, *Hoxa10*, *Hoxa11a* and *Hoxa13a*. HOXA family genes are reported to be involved in axial patterning of many structures in vertebrates, including fins and limbs(*88*), and are thus potentially functionally relevant to the Dumbo phenotype.

The lead SNP on chromosome 18 is located at 3'UTR of *FBXL15* (F-box and leucine-rich repeat protein 15) (Fig. S30). The *FBXL15* gene is a substrate recognition component of a SCF (SKP1-CUL1-F-box protein) E3 ubiquitin-protein ligase complex, which involved in the ubiquitination and subsequent proteasomal degradation of SMURF1 (*89*) which performs a crucial role in the regulation of the bone morphogenetic protein (BMP) signaling pathway in both embryonic development and bone remodeling (*89*). Thus, we consider *FBXL15* to be a strong candidate gene as well.

**Supplementary Text 9: Candidate loci in GWAS of body size**

The Giant mutant shows remarkable enlargement in body size dimensions compared to other breeds of the Siamese fighting fish (Fig. S34). By conducting GWAS using a case-control design and quantitative measurements (Supplementary Text 5), we mapped the locus to (lead SNP: chr19:2089265) at position 2.08 - 2.23 Mb on chromosome 19 and one unplaced scaffold (Oscaffold_143) (Supplementary Data 1), possibly due to an assembly error. The lead SNP on Chr. 19 explains 7.83-16.34% of the phenotypic variance (Fig. 2C). When conditioning on the top SNP chr19:2089265, all the significant associations to different body size measurements disappeared, indicating a shared mutation underlying all these phenotypes (Fig. S36). Synteny analysis of this locus with the corresponding sequence in another assembly (Accession: GCA_003650155.3) shows possible assembly errors (Fig. S43). This locus also represents one of the most gene-abundant regions across the genome (Fig. S1).

A total of 185 variants within or near 17 genes on chromosome 19 and 42 variants within or near 7 genes on Oscaffold_143 were identified (P-value < 8.46). There are 14 exonic variants in seven genes in the chromosome 19 locus (*ATP2A1*, *CRAMP1L*, *EHD1*, *MAPK8IP3*, *SLC9A3R1*, *SPSB3* and *XYLT2*) (Fig. S47) and six exonic variants in four genes in the locus on Oscaffold_143 (*ALYREF-A*, *CDR2L*, *PPP1R27*, *RECQL5*) (Supplementary Data 1). The lead SNP is in *CRAMP1L* (cramped chromatin regulator homolog 1 like), and other associated SNPs are in *MRPS34* and *SPSB3* in this locus, which are associated with human body height in GWAS catalog (https://www.ebi.ac.uk/gwas/home).

To further identify the candidate genes related to body size enlargement in the Siamese fighting fish, we explored the mRNA expression profiles of the associated genes in brain and muscle tissues between the Giant HMPK and normal HMPK (Fig. S40). We extracted the expression values of the annotated genes in the locus (Fig. S40). Significant differences in expression (padj<0.05, fold change>2) between Giant HMPK and HMPK were observed in the brain for genes *znf598*, *dhrs7b*, *ATP6V0C* and *Recql5*. In the muscle, significant differences (padj<0.05, fold change>2) were observed for genes *DRG2*, *Gfer*, *HN1L*, *MAPK8I3* and *ATP6v0C.* Intracellular non-membrane-bounded organelle, non-membrane-bound organelle and cytoplasmic part GO terms were significantly enriched (|log2foldchange|>0, padj<0.05) in the differentially expressed genes in muscle between the Giant-HMPK and HMPK (Fig. S41). Cytoplasmic part was also the top enriched GO terms (|log2foldchange|>0, padj<0.05) based on the DEGs in the brain between the Giant-HMPK and HMPK (Fig. S42).

We mapped another three body-size related traits, BMI, Total length/Height and Weight/Height (Fig. S38). GWAS result of Total Length/Height replicates the *KCNJ15* long-fin locus on chromosome 17 (Fig. 3A), and GWAS results of BMI and Weight/Height replicate the body size locus on chromosome 19 (Fig. S35). The Giant variety in the Siamese fighting fish provides an excellent model for studying the Mendelian genetics of body size, which is usually a polygenic based phenotype.

#### Supplementary Text 10: Candidate loci in GWAS of coloration

GWAS on the solid red, solid yellow and solid orange color (Supplementary Text 5) identified a strong signal at 5.83 Mb on chromosome 19, and GWAS schemes involving different comparisons can replicate this signal (Fig 2B, S44, S45). In the peak region, there are a total of 93 variants located on or in the vicinity of *RNF213* gene, consisting of 58 exonic and 35 intronic SNPs (Fig. S45). The *RNF213,* known as mysterin, encodes a large cytoplasmic protein as long as 5207 amino acids that contains two AAA+ ATPase modules and a RING finger ubiquitin ligase domain, involved in angiogenesis and the non-canonical Wnt signaling pathway in vascular development (*90*). There is no obvious direct connection between the function of *RNF213* and the pigmentation phenotype. However, these variants may alter the ATPase modules or RING finger domain of the *RNF213* gene related to the non-canonical Wnt signaling pathway, possibly leading to transport, deposition and metabolism of the erythrophores and xanthophores.

Hybridization between Turquoise-green and Steel-blue breeds produces F1s with Royal-blue color, an inheritance pattern that was first documented in the 1930s (*60*). We successfully mapped this mendelian locus to 5.03-5.26 Mb on chromosome 20 using a case-control GWAS (Fig 2B). There are 13 protein coding genes in this locus including *MTHFD1L* (methylenetetrahydrofolate dehydrogenase 1 like) gene (Fig. S48). A previous study reports that the *MTHFD1L* gene is involved in the synthesis of tetrahydrofolate in the mitochondrion (*91*). Tetrahydrofolate is engaged in the de novo synthesis of purines, a key component of iridophores (*92*). As the Royal-blue, Steel-blue, and Turquoise-green are structural colors, *MTHFD1L* gene is a promising candidate in the formation of the three colors. More experiments including chromatography identification and sections are needed in the future.

The Copper breed is derived by introducing two copies of the "metallic gene" into the genetic background of the Steel-blue breed (Fig. S49), and the "metallic gene" locus is independent of the locus (5.03-5.26 Mb on chromosome 20) underlying the Turquoise-green, Royal-blue and Steel-blue variation. Based on this knowledge, we performed a case-control GWAS on the Steel-blue and Copper breeds (Fig. S50). Surprisingly, we identified two significantly associated loci, one is the same locus underlying the Royal-blue, Steel-blue and Turquoise-green color variation, the other is a novel one on chromosome 7. This result confirms the observation suggesting an independent "metallic gene" underlies the distinctive Copper phenotype.

Further analysis is needed to pinpoint the “metallic gene” within the locus on chromosome 7.

The Siamese fighting fish also displays considerable diversity in eye color. To map the genetic basis of this phenotypic variation, we classified them into six colors including black, white, yellowish-brown, yellow, light blue and brown and conducted GWAS using the eye color as categorical phenotypes. However, no significant loci were detected (Fig. S51), perhaps suggesting a complex genetic architecture or environmental cues underlying eye color variation.

The Mosaic color pattern is a spectacular phenotype in the Siamese fighting fish; many commercial names are present for such a pattern, including koi, candy, galaxy, lemon and marble. Several hypotheses are proposed for the Mosaic color pattern, including the “jumping genes” hypothesis(*93*). Although the mosaic color pattern was also recorded in koi carp(*94*) and medaka (*95*), genetic studies on this phenotype are scarce. By classifying the solid colors as controls and the mosaic color as cases, we performed a case-control GWAS and revealed a polygenic basis underlying this phenotype (Fig. 2B, S46). Nine loci on eight chromosomes are identified, with two adjacent major loci on chromosome 16 (Supplementary Data 1, Table S12). The most significant locus (22.64-941.2 Kb) on chromosome 16 with the lead SNP (chr16:74988) contains 1,517 associated variants (P-value $< 8.46$) and 39 genes. *SLC39A7*, *CHS3*, *CHS8*, *COL11A2* and *TUBB* in this region are candidate genes related to pigmentation. *TUBB* is related to intracellular pigment mobilization (*96*) making it a promising candidate for the mosaic phenotype. The adjacent locus at 1.76-2.98 Mb contains 1,182 associated variants (P-value $< 8.46$) and 47 genes. *PLEC*, *EPPK1*, *SLC17A5* and *SLC52A2* are candidates worthy of further investigation. Surprisingly, there are eight *PLEC* gene copies in the region, and there are 346 variants (P-value $< 8.46$) within/near the eight copies, with 62 exonic and 245 intronic. *PLEC* (plectin) plays a role as plakins or cytolinkers, which link different elements of the cytoskeleton. It was proposed to be the top hub gene associated with the “Pink-dark green” color alteration in a gene coexpression network analysis in the cavefish (*97*). Reduced expression of *PLEC* causes increased melanosome uptake by keratinocytes and skin hyperpigmentation after UVA exposure in humans (*98*). Given the functional relevance, we speculate that the multiple *PLEC* copies could be the main driver leading to the mosaic pattern in the Siamese fighting fish. Two other genes in this region, *SLC17A5* and *SLC52A2*, are also considered pigment related genes described in (*99*, *100*). The region (9.82-10.68 Mb) on chromosome 2 contains 189 associated variants (P-value < 8.46) and 39 protein coding genes. *SLC2A11*, *SLC8B1* and *AIDB* are potential candidate genes. *SLC2A11* is related to xanthophore differentiation in goldfish(*101*). There are 66 associated variants between 1.66-2.64 Mb on chromosome 10. Within this region, *SLC23A2* is a candidate pigment-determining gene in the yellow-feathered chickens(*102*). Our GWAS results comparing the solid color breeds and the mosaic color breeds illustrate a polygenic picture of the mosaic pattern. This analysis represents a first attempt to map the mosaic pattern in fish, and also helps unveil the genetic mechanism of this intriguing phenotype.

**Supplementary Text 11: Candidate loci in GWAS of aggression behaviours**

Aggression is a complex behavioral trait with neurophysiological, endocrinological, physiological and psychological bases. The Siamese fighting fish was intensely selected for aggression for centuries during domestication. It is one of the few animals selected for more aggressiveness rather than tameness, thus providing an excellent model for studying this complex trait. Here we performed an in-depth phenotypic investigation of aggressiveness and characterized the genetic architecture of aggression in the Siamese fighting fish.

GWAS signals for Fighter vs non-Fighter breeds

GWAS using the aggressive fighter breeds and non-fighter in a case-control design identified multiple loci across the genome significantly associated (Fig. 4A). We discuss possible candidate genes on these loci below: On the 19.32 Mb position of chromosome 1, 15 significantly associated SNPs (P-value < 8.46) are located in the intron of *CACNB2* (calcium channel, voltage-dependent, beta 2 subunit) gene which controls the calcium channel by altering the voltage dependencies and modulating G protein inhibition in the axon terminal(*103*). Calcium entering through this channel is essential for several cellular processes, like hormone release, neurotransmitter release, pacemaker activity and cardiac muscle contraction (*104*). Variants in *CACNB2* are shown to be associated with several psychiatric disorders in humans and mice, such as autism spectrum disorder, major depressive disorder, bipolar disorder and schizophrenia (*103, 105, 106*). *CACNB2* was reported to have the most splice variants among the *CACNB* genes, and intronic variants in *CACNB2* are associated with working memory and brain activity (*107*).

Another locus at around 5.16 Mb on chromosome 13 contains 105 associated variants (15 coding and 90 non-coding, P-value < 8.46). A gene within this locus, *AVPR2* (arginine vasopressin receptor 2), encodes the receptor for arginine vasopressin, which reabsorbs water from urea and contributes to maintenance of water homeostasis(*108*). A paralogous gene of *AVPR2*, *AVPR1A* (arginine vasopressin receptor 1a), is known to contribute to several behavior diseases in humans, including autism (*109*), stress management (*110*) and territorial aggression (*111*). *AVPR1A* knockout mice display reduced anxiety, impaired reciprocal social interaction and decreased social recognition (*112*, *113*). There is no previous study that relates *AVRP2* to behavioral related traits yet.

Another signal spanning from 8.37 to 8.65 Mb on chromosome 17 contains three genes named *RYR2* (ryanodine receptor 2, three intronic SNPs), *NFRKB* (nuclear factor related to kappa-B binding protein, one exonic variant) and *NRXN2* (neurexin-2, two exonic and eight intronic SNPs). *RYR2* encodes ryanodine receptors, whose function is to form channels to transport charged calcium atoms within myocytes. The routine control of calcium ions within myocytes governs the cycle of muscle contraction and relaxation(*114*). We detected higher expression levels of the *RYR2* in the brain and muscle of the Fighters than Non-Fighters (Supplementary Data 4), which might be related to an increased physiological capacity for fighting behaviours such as biting, active motion, and longer fighting duration. *NFRKB* is functionally similar to *NFKB1* (nuclear factor kappa B subunit 1), which is a candidate gene associated with anger(*115*) and antisocial behavior(*116*). *NRXN2* is engaged in cell recognition and adhesion activities by encoding neuronal cell surface proteins. *NRXN2A* KO mice displayed deficits in sociability and social memory and an anxiety-like phenotype(*117*). All three genes remain plausible candidate genes and further functional analyses are needed to determine which variants in these genes might be causally related to fighting behavior.

The associated locus on chromosome 11 from 6.68 to 6.93 Mb contains a gene named *PCDH15* (protocadherin 15). PCDH15 is one of the cadherin superfamily and is associated with several psychiatric disorders (*118*). Other genes, including *BFSP1*, *PLEK* and *PROKR2* in this region are involved in coding the components of cytoskeleton, potentially suggesting that variations related to cytoskeleton have contributed to the aggressiveness of the Fighters.

Another locus on chromosome 11 from 11.68 to 11.72 Mb contains *DISC1* (disrupted in schizophrenia 1). This gene was identified as a causal gene responsible for schizophrenia, bipolar disorder, and recurrent major depression in a large Scottish family(*119*).

The associated locus around 2.24 Mb on chromosome 2 contains *CALD1*, which played an important role in hypoxic adaptation by mediating blood vessel development, energy metabolism through angiogenesis, vascular smooth muscle contraction, and various hypoxia-related signaling pathways during embryonic development in Tibetan chicken(*120*). This gene might be related to the hypoxia endurance in the Fighters.

On the associated peak region on chromosome 5, 3 intronic and 6 surrounding variants (P-value < 8.46) for *GALNT2* (polypeptide N-acetylgalactosaminyltransferase 2) are found to be significantly associated with the Fighter phenotype. This gene plays an important role in initiation of mucin-type protein O-glycosylation(*121*). Other genes, including, *FGF18*, *MAPK3*, *MAP6* and *STX2* are also candidate genes reported in previous studies(*122*)worthy of further investigation for aggression in the Fighter breeds.

GWAS on the quantitative indices of aggression

To obtain a quantitative insight into the aggressiveness of the Siamese fighting fish, we designed an experiment to quantify the aggressiveness of each fish using 10 indices (Supplementary Text 4). In a GWAS on these measurements, we identified 32 loci (P-value < 8.46) on 16 chromosomes associated with eight aggressiveness phenotypes (Supplementary Data 1). Among these results, one locus at 12.28 Mb on chromosome 2 is associated with Charge score and Mouth open score simultaneously (Fig. 4K, 53E). One gene named *GFRA2* within this locus regulates the neuron survival and differentiation(*123*), would be a candiage gene. Another locus overlapped in Charge score and Mouth open points to a region on chromosome 1 near a gene named *BRINP3*. This gene is a member of membrane attack complex superfamily and genetic variation in its paralogs (i.e., *BRINP2*, *BRINP1*) exhibit behavior reminiscent of autism spectrum disorder and attention deficit hyperactivity disorder(*124*). The top SNP (chr16:2116016, P-value = 6.01E-16) on chromosome 16 locates in introns of *MACF1* (microtubule-actin cross-linking factor 1). *MACF1* is engaged in multiple neural processes and mutations or dysregulation of this gene is associated with schizophrenia and Parkinson's disease(*124*). More candidate genes worthy of refined investigations are marked in Fig. S53. Although we observed few associations overlapped between GWAS results of case-control and quantitative traits for aggression, they are complementary to support each other for the fact that most associated genes relate to human psychiatric disorders, suggesting a similar molecular network between human psychiatric disorders and the domestication of aggressiveness in the Siamese fighting fish.

### Supplementary Data

**Supplementary Data 1 Results of genome-wide association studies of all the recorded traits in the Siamese fighting fish.**

**Supplementary Data 2 Results of ABBA-BABA tests.**

**Supplementary Data 3 Expanded and contracted gene families in the Siamese fighting fish.**

**Supplementary Data 4 Differentially expressed genes in the brain between Fighter and Non-Fighter.**

**Supplementary Data 5 Functional enrichment of the identified genes by GWAS between the Fighter and Non-Fighter breeds.**
